# Disinhibition of the orbitofrontal cortex biases decision making in obesity

**DOI:** 10.1101/2020.04.02.022681

**Authors:** Lauren T. Seabrook, Lindsay Naef, Corey Baimel, Allap K. Judge, Tyra Kenney, Madelyn Ellis, Temoor Tayyab, Mataea Armstrong, Min Qiao, Stan B Floresco, Stephanie L. Borgland

## Abstract

The lateral orbitofrontal cortex (lOFC) receives sensory information about food and integrates these signals with expected outcomes to guide future actions, and thus may play a key role in a distributed network of neural circuits that regulate feeding behaviour. Here, we reveal a novel role for the lOFC in the cognitive control of behaviour in obesity. Food-seeking behaviour is biased in obesity such that in male obese mice, behaviours are less flexible to changes in the perceived value of the outcome. Obesity is associated with reduced lOFC inhibitory drive, and chemogenetic reduction in GABAergic neurotransmission in the lOFC induces obesity-like impairments in goal-directed behaviour. Conversely, pharmacological or optogenetic restoration of inhibitory neurotransmission in the lOFC of obese mice reinstates flexible behaviour. Our results indicate that obesity-induced disinhibition of the lOFC leads to a failure to update changes in the value of food with satiety, which in turn may influence how individuals make decisions in an obesogenic environment.

## Introduction

Diet-induced obesity; along with metabolic disease, is a major health concern for many individuals around the world. Obesity is currently defined as having a body mass index of greater than 30 kg/m^2^ and is associated with multiple comorbid diseases including type 2 diabetes, stroke, cancer and depression^1^. Numerous forces drive an increase in obesity rates, including complex environmental and societal changes^2^. Moreover, the transition from consuming traditional nutritional foods to highly marketed, inexpensive, energy dense, palatable foods has exacerbated overeating^3^. These pre-packaged foods are often consumed despite already fulfilled energy requirements^4^ as they entice our innate likings of sugars, salts, and fats^5^. Quantitative models have been used to calculate the efficiency of obesity prevention efforts, including the impact of individual behaviours, public health interventions, and government policies. It is incumbent on the individual to “make personal healthy food and activity choices”^6^, yet the complexity of the modern food environment and the choices it offers biases decision-making to influence food choices. Prior literature proposes that readily available palatable and energy dense foods usurps an individual’s ability to make decisions and control their caloric intake, leading to overconsumption and obesity^7^. How access to obesogenic food alters neural circuits to bias behaviour towards eating beyond satiety remains unclear.

During goal-directed behaviour, decisions and behavioural strategies are flexible as they track the relationship between actions and outcomes. Individuals will adjust their behaviour upon a change in the value of an outcome previously associated with the action. For example, a food reward can be devalued with satiety or by pairing the food with cues predicting sickness, as in conditioned taste avoidance. The development of feeding behaviour that is insensitive to changing reward values has been implicated in obesity, such that humans with obesity^8^ and obese rodents^9^ demonstrate deficits in outcome devaluation.

The lateral OFC (lOFC) is anatomically and functionally situated to influence food intake as it is reciprocally connected with sensory^10^, motor ^11–13^, and limbic ^14,15^ brain regions and is thought to integrate information from these regions to guide decision-making. Several lines of evidence suggest that an intact lOFC is required for goal-directed behaviour, as the disruption of the lOFC by lesions or inactivation impairs devaluation by satiety^11^, sickness^16^ and contingency degradation^17^. A current hypothesis is that value representations of food and decision-making mechanisms to control food-intake are disrupted in obesity^18^, however, it is unknown if obesity influences the function of the lOFC and how this occurs. Here, we tested the hypothesis that diet-induced obesity impairs goal-directed behaviour by disrupting lOFC function.

## Results

### Obesity impaired devaluation

To determine if goal-directed value representations of food are impaired in obesity, we adapted three devaluation paradigms commonly used to examine changes in goal-directed behaviour^19^. Long-term exposure to a high-fat diet led to the development of diet-induced obesity noted by increased body weight, hyperglycemia, as well as decreased glucose clearance reflective of impaired insulin signalling (**Extended Data Figure 1a-d**). Because some of our behavioural paradigms require mild food restriction, we examined these measures after food restriction to 15% of body weight. The same pattern was observed.

We first examined the impact of diet-induced obesity on devaluation by satiety. Lean and obese mice were trained to lever press for liquid sucrose rewards on a random ratio (RR) 20 schedule of reinforcement, whereby sucrose delivery followed on average the 20^th^ lever press (**Extended Data Figure 2a,b**). Once mice displayed stable levels of responding, we devalued the reward outcome (sucrose) by pre-feeding them to satiety with the same sucrose solution, and then tested them in a non-reinforced instrumental test session (**Extended Data Figure 2c,d**). Lean mice were sensitive to devaluation (**Extended Data Figure 2e**), indexed by reduced responding when pre-fed with sucrose in the devalued condition, relative to non-prefed (valued) conditions. In contrast, obese mice were impervious to the effects of the change in value of sucrose, as they had comparable levels of responding under both valued and devalued conditions (**Extended Data Figure 2e**). This effect was observed when we computed the revaluation index (lever presses valued state – lever presses devalued state) / (lever presses valued state + lever presses devalued state) which indicates the degree of goal-directedness during outcome revaluation, such that a positive revaluation index reveals the strength of devaluation (**Extended Data Figure 2f).** Total lever presses were not different between groups (**Extended Data Figure 2g**). We confirmed this by reanalyzing our devaluation data in lean and obese mice that had matched total lever presses on the last day of training. Even under these stricter inclusion criteria, lean mice display devaluation, whereas obese mice do not (**Figure 1a-d**). This suggests when controlled for learning, actions of lean mice track the reward value, whereas obese lever press regardless of the reward value.

**Figure 1:**
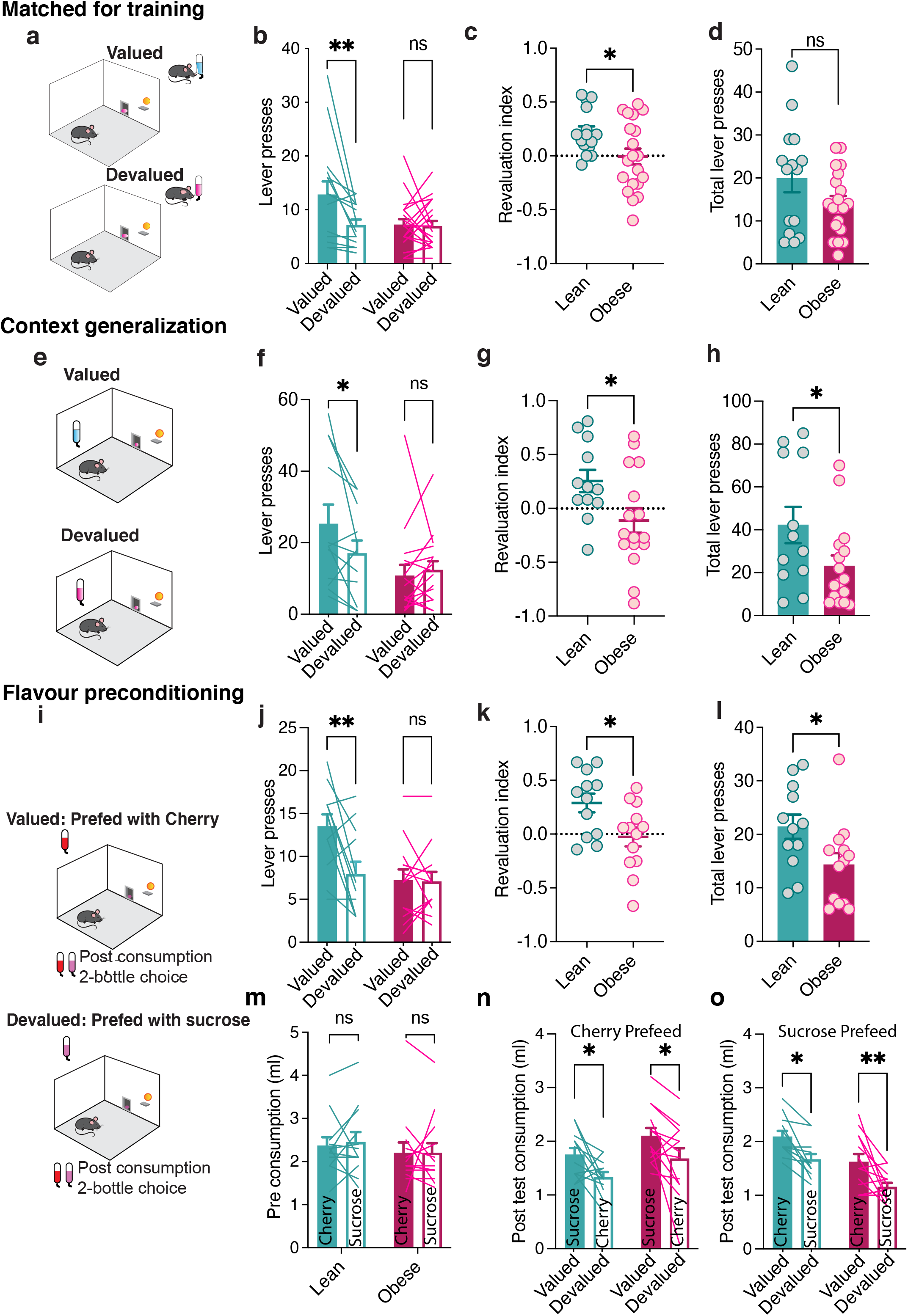
Diet-induced obesity induces deficits in satiety-induced devaluation. a) Satiety-induced devaluation procedure in lean and obese mice matched for lever pressing. b) Lean (n=15) mice displayed devaluation as indicated by decreased lever presses in the devalued compared to the valued condition. Obese mice (n=20) did not display devaluation as indicated by comparable lever presses in the valued and devalued conditions. RM two-way ANOVA: Devaluation effect, F (1,33) = 7.098, p=0.0118*, Diet effect F (1,33) = 2.919, p=0.0970, Diet x devaluation interaction, F (1, 33) = 5.949, p=0.0203*, Sidak’s multiple comparisons test, lean: p= 0.0038**, obese: p = 0.9816. Data are presented as mean values +/− SEM. c) Obesity decreased the revaluation index ((lever presses valued – lever presses devalued)/ (lever presses valued + lever presses devalued)). Unpaired t-test: lean (n=15), obese (n=20), t_(33)_ = 2.327, p=0.0262*. Data are presented as mean values +/− SEM. d) Matched number of lever presses in lean mice (n=15) and obese mice (n=20) based on last training day of RR20, excluded mice that pressed <30 and >100. Unpaired t-test: t_(33)_ = 1.708, p=0.0970. Data are presented as mean values +/− SEM. e) Context generalization procedure for devaluation. f) Lean (n=12) mice displayed devaluation when sucrose pre-consumption test was in same environment as test, obese mice (n=16) did not. RM two-way ANOVA: Devaluation effect, F (1, 26) = 4.239, p=0.0497*, Diet effect F (1,26) = 2.399, p=0.1335, Diet x devaluation interaction, F (1, 26) = 5.330, p=0.0292*, Sidak’s multiple comparisons test, lean: p = 0.0336*, obese: p = 0.8123. Data are presented as mean values +/− SEM. g) Obesity decreased the revaluation index. Unpaired t-test: lean (n=12), obese (n=16). Unpaired t-test t_(26)_ = 2.293, p=0.0302*. h) Total number of lever presses (valued + devalued) of lean (n=12) and obese (n=16) during the devaluation task. Unpaired t-test: t_(26)_ = 2.059, p=0.0497*. i) Calorie matched flavour induced devaluation and post consumption test procedure. j) Lean (n=12) but not obese mice (n=13) display devaluation as indicated by increased lever presses in the valued and devalued conditions. RM two-way ANOVA: Devaluation effect F (1, 23) = 8.642, p=0.0074**, Diet effect F (1,23) = 5.066, p=0.0343*, Diet x devaluation interaction, F (1, 23) = 7.740, p=0.0106 *, Sidak’s multiple comparisons test, lean: p = 0.0012**, obese: p = 0.9920. k) Obesity decreased the revaluation index. Unpaired t-test: lean (n=12), obese (n=13). Unpaired t-test t_(23)_ = 2.562, p=0.0174* l) Total number of lever presses of lean (n=12) and obese (n=13) during the devaluation task. Unpaired t-test: t_(23)_ = 2.251, p= 0.0343*. m) Lean (n=12) and obese mice (n=13) pre-test consumption. RM two-way ANOVA: Consumption effect F (1,23) = 0.08699, p=0.7707, Diet effect F (1,23) = 0.5220, p=0.4773 Diet x consumption interaction, F (1,23) = 0.08699, p=0.7707. n) Both lean (n=12) and obese mice (n=13) drink less cherry sucrose compared to plain sucrose when pre-fed with cherry. RM two-way ANOVA: Devaluation effect F (1,23) = 12.37, p=0.0018**, Diet effect F (1, 23) = 4.028, p=0.0566, Diet x devaluation interaction, F (1, 23) = 0.0007210, p=0.9788, Sidaks multiple comparisons test, lean: p = 0.0471*, obese: p = 0.0349*. o) Both lean (n=12) and obese mice (n=13) drink less plain sucrose compared to cherry sucrose when pre-fed with sucrose. RM two-way ANOVA: Value effect F (1, 23) = 21.94, p=0.0001***, Diet effect F (1, 23) = 15.72, p=0.0006***, Diet x devaluation interaction F (1, 23) = 0.05368, p=0.8188, Sidaks multiple comparisons test, lean: p = 0.0104*, obese: p = 0.0034**.

To determine if obese mice are unable to update the response outcome association because they could not recall that they were pre-fed sucrose as it was given in a different context, we next tested if obese mice were generalizing the devaluation setting. To test this, mice were pre-fed with sucrose or water in the operant chambers where they are tested for devaluation (**Extended Data Figure 2h,i)**. Under these conditions, lean mice again devalued sucrose whereas obese mice did not; evident by decreased lever pressing in the devalued state as well as a positive revaluation index (**Figure 1e-h)**. Additionally, when responses during the devaluation session were reinforced, lean but not obese mice adjusted their lever pressing in the devalued condition (**Extended Data Figure 3a-d)**, suggesting that even when obese mice are reminded of the pre-sated reward, they continue to lever press. This difference in responding is not due to motivational state because when tested on a progressive ratio schedule of reinforcement, lean and obese mice displayed comparable levels of responding, sucrose consumption and breakpoints for sucrose (**Extended Data Figure 3e-h**). Together this suggests that regardless of motivational state or when obese mice are not required to remember the devaluation setting or flavour, they continue to show comparable responding in valued and devalued states.

We then examined the impact of diet-induced obesity on a contingency change task, which measures the relationship between action and outcome. Mice were trained on a positive contingency whereby increased lever presses yielded increased sucrose delivery (nondegraded, ND). During testing, we degraded the lever contingency (CD) so that sucrose delivery was not contingent on lever pressing and delivered in a variable interval 20 schedule, whereby the reward was administered approximately every 20 seconds regardless of whether the mice pressed the lever. Lean mice significantly decreased their lever pressing to the degraded contingency, whereas this did not reach statistical significance in obese mice (**Extended Data Figure 3i**).

The satiety induced devaluation procedure in Figure 1a-d confounds general satiety with sensory-specific devaluation of the outcome. Therefore, to determine if satiety influences devaluation differently in lean and obese mice, we next tested devaluation using a sensory-specific satiety devaluation procedure, whereby mice were pre-fed with either an isocaloric cherry-flavoured solution (valued) or unflavoured sucrose (devalued) and then lever pressed for the expected sucrose outcome. Lean mice devalue when pre-fed with a cherry solution, showing sensory specific satiety that influences goal-directed behaviour to attenuate responding for a specific tastant. In contrast, obese mice showed comparable lever pressing when the outcome was valued vs. when it was devalued (**Figure 1i-l)**. This is not because of flavour preference, as both lean and obese equally consumed cherry and sucrose solutions in a pre-consumption test (**Figure 1m**). Moreover, in a two-bottle choice post-consumption test, both lean and obese mice drank less of the flavour they were pre-fed with (**Figure 1n,o)** indicating that lean and obese mice were satiated to a similar extent during the prefeed. Taken together, these data indicate that while lean mice reduce their actions for the expected outcome when pre-sated with the same flavour, obese mice display impairments in this aspect of goal-directed behaviour.

We next examined the impact of diet-induced obesity on sickness-induced devaluation. Prior to dietary manipulation, mice were conditioned over three sessions to associate either grape or orange flavoured gelatine with lithium chloride (LiCl)-induced malaise (**Figure 2a-c**). Pre-diet exposure, both groups consumed significantly less of the LiCl-paired gelatine flavour indicated by a decreased consumption of the gelatine flavour that was paired with sickness and a positive revaluation index (**Figure 2c-f**), suggesting effective pairing of LiCl-malaise and flavour. In contrast, after 3 months exposure to low or high fat diets, lean mice maintained conditioned taste avoidance to the LiCl-paired flavour, whereas obese mice consumed comparable amounts of the valued and devalued gelatine (**Figure 2g-i**). However, while we did not observe memory impairments in our other experiments, these data do not exclude the possibility that obese mice forget the original association after diet exposure. In summary, obese mice displayed impaired sickness-induced devaluation by conditioned taste avoidance.

**Figure 2:**
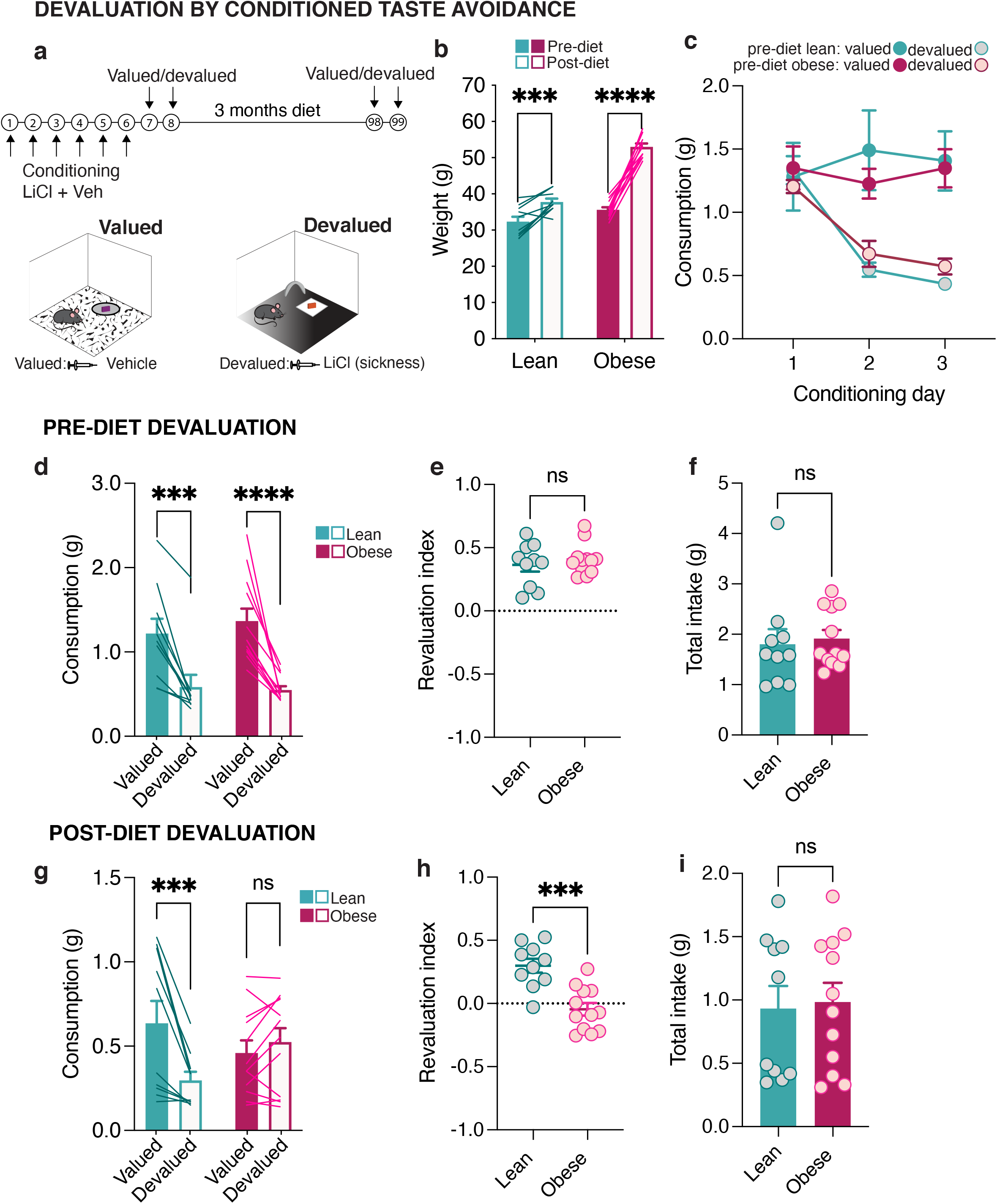
Diet-induced obesity induces deficits in conditioned taste avoidance. a) Conditioned taste avoidance devaluation procedure. b) Weights of mice in the CTA experiment during conditioning (pre-diet) and following exposure to either the low-fat (n=10) or high-fat diet (n=12). RM Two-way ANOVA: Diet effect F (1,20) =75.01, P<0.0001, Time effect F (1,20) =179.3, P<0.0001****, Diet x time interaction F (1,20) =49.20, p<0.0001****, Sidak’s multiple comparisons test: pre-diet lean vs. post-diet lean p= 0.0007*** pre-diet obese vs post-diet obese p<0.0001****. Data are presented as mean values +/− SEM. c) Consumption (g) of the valued and devalued gelatine during the three days of taste avoidance conditioning (pre-diet) of lean (n=10) and obese mice (n=12). Three-Way ANOVA: Day x devaluation interaction: F (1.391,27.82) = 12.33, p=0.0006, Day effect: F (1.832,36.63) = 10.84, p=0.0003, Valued vs. devalued effect: F (1,20) = 24.42, p<0.0001, diet effect: F (1,20) = 0.02218, p=0.8831. Tukey’s multiple comparisons test: Pairing Day 3 lean valued vs. devalued: p = 0.0379*, Pairing Day 3 obese valued vs. devalued: p = 0.0015**. Data are presented as mean values +/− SEM. d) Consumption (g) of the valued and devalued gelatine during the CTA test prior to exposure to the diets (lean n=10, obese n=12). RM Two-way ANOVA: Diet x devaluation interaction: F (1,20) = 0.9552, p = 0.3401. Devaluation effect F (1,20) = 60.98, p<0.0001****, diet effect: F (1,20) = 0.1227, p=0.7297, Sidak’s multiple comparisons lean: p=0.0003***, obese: p<0.0001****. Data are presented as mean values +/− SEM. e) Revaluation index of pre-diet lean (n=10) and pre-diet obese (n=12) mice. Unpaired t-test: t _(20)_ =0.6773, p=0.5060. Data are presented as mean values +/− SEM. f) Total intake (valued + devalued) of gelatine in pre-diet lean (n=10) and pre-diet obese (n=12). Unpaired t-test: t_(20)_ =0.3503, p=0.7297. Data are presented as mean values +/− SEM. g) Lean (n=10) mice displayed devaluation as indicated by a decreased consumption of devalued compared to valued gelatine after exposure to the diets. Obese (n=12) mice were insensitive to devaluation as indicated by comparable valued and devalued gelatine consumption. RM two-way ANOVA, devaluation effect: F (1, 20) =8.210, p=0.0096**, diet effect: diet x devaluation interaction: F (1, 20) =17.92 p=0.0004***. Sidak’s multiple comparison test, lean: p=0.0002***, obese: p=0.5410. Data are presented as mean values +/− SEM. h) Obesity decreased the revaluation index. Unpaired t-test: lean (n=10), obese (n=12) mice, t_(20)_ =4.681, p=0.0001***. Data are presented as mean values +/− SEM. i) Total consumption (valued + devalued) of gelatine in post-diet lean (n=10) and post-diet obese (n=12) mice. Unpaired t-test: t_(20)_ =0.2268, p=0.8229. Data are presented as mean values +/− SEM.

We next examined if impairments in devaluation in obese mice were due either to recent exposure to the energy-dense diet or to obesity. To test this, both lean and obese mice were switched to the control diet for 7 days during training and prior to testing for satiety-induced devaluation. Obese mice maintained their significant weight difference from lean mice during the low-fat diet (**Extended Data Figure 4a,b**). After 7 days low-fat diet exposure, when pre-fed with sucrose (**Extended Data Figure 4c)** lean mice devalued to the expected outcome, whereas obese mice did not (**Extended data Figure 4d-f)**. In summary, these data demonstrate clear discrepancies in the value attributed to food rewards and related actions by lean and obese mice. While lean mice change their behaviour depending on internal state and prior experiences, obese mice show marked impairments in behavioural adjustment to the current value of food rewards, lasting beyond the duration of obesogenic diet exposure.

### Obesity alters the function of lOFC neurons

The lateral OFC (lOFC) has emerged as a hub for assessing information about rewarding outcomes and orchestrating flexible, goal-directed behaviour^11,20^. Our previous work demonstrated that obese rats fed a cafeteria diet have changes in inhibitory and excitatory transmission in the lOFC^21,22^. Furthermore, obesogenic diets can also alter the intrinsic excitability of neurons, as demonstrated in the nucleus accumbens^23^ or dorsal raphe^24^. To examine the effects of obesity on the lOFC, we performed whole-cell electrophysiology recordings in brain slices containing the lOFC from lean and obese mice and measured the number of action potentials in response to current steps of increasing amplitude and calculated the frequency-current (F-I) distribution as well as an excitability slope as a measure of the relative excitability of layer II/III lOFC pyramidal neurons. lOFC pyramidal neurons from obese mice were more excitable than those of lean mice (**Figure 3a-c**). This effect persisted when both lean and obese mice were food restricted (**Extended data Figure 5a-e**). An increase in excitability could be due to several mechanisms; a change in intrinsic properties, an increase in excitatory inputs, a decrease in presynaptic GABA release probability, or of tonic GABA regulation of lOFC pyramidal neurons. We ruled out changes in excitatory inputs as unlike layer V pyramidal neurons, which receive inputs from subcortical regions, layer II/III pyramidal neurons are primarily innervated by intracortical projections at distal dendrites and are more readily controlled by the coordinated action of local inhibitory interneurons^25^. Furthermore, our previous work demonstrated that an obesogenic diet decreased presynaptic glutamate release onto OFC neurons via mGlur2/3 activation^21^, which would be unlikely to increase firing. Therefore, we first assessed if obese mice had altered passive membrane properties. There were no differences in the cell capacitance (**Extended Data Figure 6a**), resting membrane potential (**Extended Data Figure 6b**), action potential height and width (**Extended Data Figure 6e,f**), threshold to fire (**Extended Data Figure 6g**), or after hyperpolarization potential height (**Extended Data Figure 6h**) in lean or obese mice in the presence or absence of the GABA_A_ inhibitor, picrotoxin. However, we did observe an increase in input resistance (**Extended Data Figure 6c**), decrease in rheobase (**Extended Data Figure 6d)** and a decrease in AHP width (**Extended Data Figure 6i**) of obese OFC neurons. We next tested if GABAergic disinhibition underlies the enhanced excitability of pyramidal neurons. Picrotoxin-induced inhibition of GABAergic transmission increased the excitability of pyramidal neurons from lean, but not obese mice (**Figure 3b,c**), suggesting that increased neuronal excitability of obese mice may be due to decreased inhibition. Further, the increase in input resistance, decrease in rheobase and AHP width was blocked by picrotoxin, suggesting that these changes are secondary to action at GABA_A_ receptors. We next isolated and quantified miniature inhibitory postsynaptic currents (mIPSCs) onto lOFC pyramidal neurons. The frequency, but not the amplitude of mIPSCs were decreased in obese relative to lean mice in lOFC pyramidal neurons (**Figure 3d-f**). We also observed a paired pulse facilitation (**Figure 3g,h**), suggesting a decrease in presynaptic GABA release probability onto lOFC pyramidal neurons of obese mice, consistent with previous findings from our group^21,22^.

**Figure 3.**
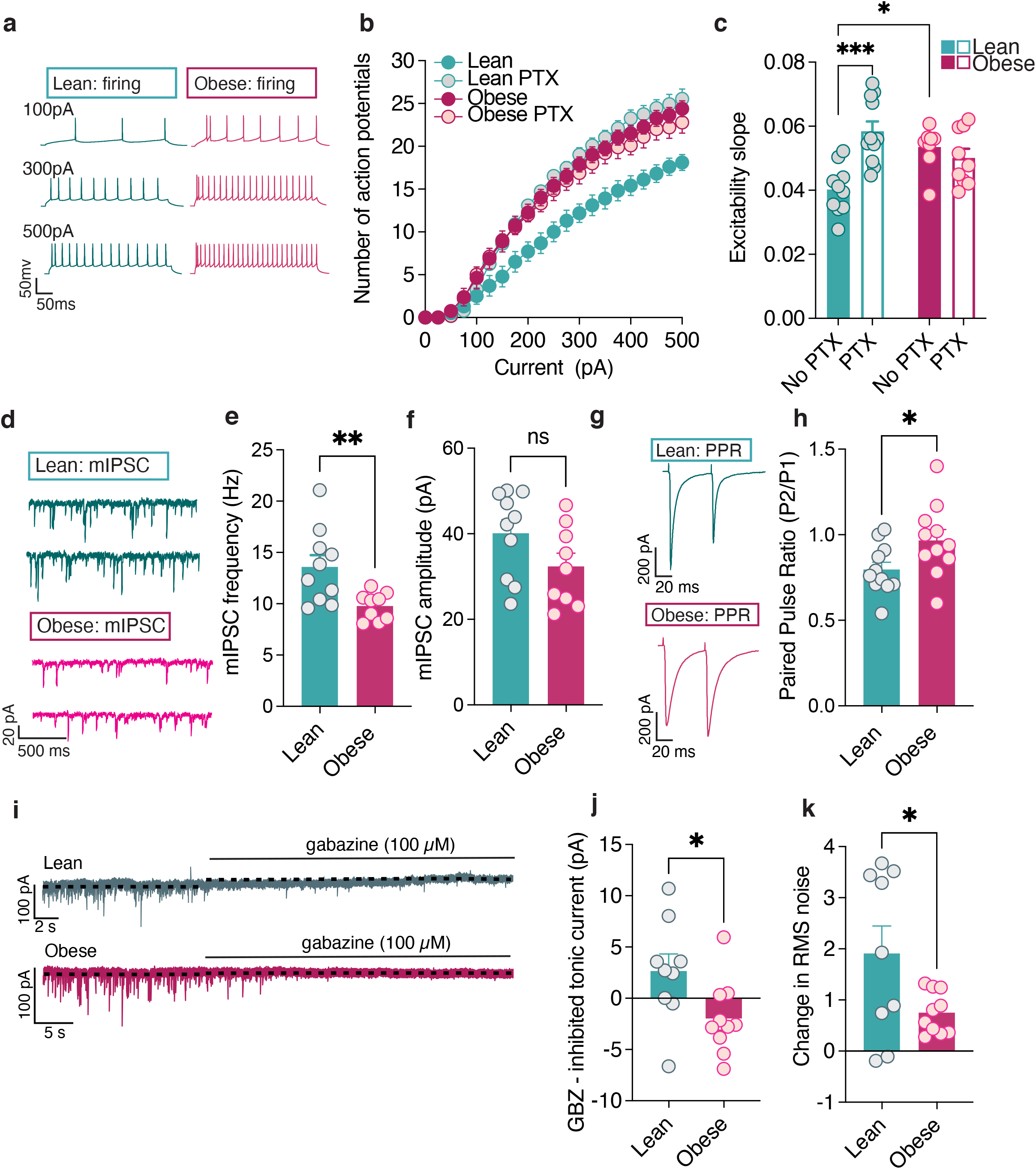
Obesity reduces inhibitory tone in the lOFC and disinhibits principal output neurons. a) Representative recordings of action potentials observed at 100pA, 300pA and 500pA current steps from lOFC pyramidal neurons of lean and obese mice. b) Diet-induced obesity increased the excitability of lOFC pyramidal neurons as indicated by frequency-current (F-I) plots of number of action potentials at current injections from 0 pA to 500pA of lean (n=10 cells/3 animals) and obese (n=8 cells/3 animals) mice. In the presence of picrotoxin, obese mice (n=9/5) excitability but not lean mice (n=11/3) were altered. RM 3-way ANOVA: current step effect F (20.00, 360.0) = 779.8 p<0.0001****, PTX vs vehicle effect F (0.09746, 1.754) = 7.656 p=0.0653, Diet effect: F (1, 18) = 4.325 p= 0.0521, current step x drug interaction F (2.261, 35.94) = 5.372, p= 0.0071**, Current step x diet interaction F (20, 360) = 1.406 p= 0.1156, Drug x diet interaction F (1, 18) = 12.56, p= 0.0023**, Current x Drug x Diet interaction F (20, 318) = 9.668 p= <0.0001****. Data are presented as mean values +/− SEM. c) Diet-induced obesity increased the excitability of lOFC pyramidal neurons and picrotoxin-induced disinhibition only changed the firing of pyramidal neurons from lean mice. Mean excitability slope (slope of linear regression from individual cells, x=current step y=number of action potentials) of lOFC pyramidal neurons of lean in aCSF (n=10 cells/3 mice), lean in picrotoxin (n=11 cells/3 mice), obese in aCSF (n=8 cells/3 mice), obese in picrotoxin (n=9/5 mice). Two-way ANOVA: picrotoxin effect F (1,34) = 7.156 p=0.0114*, picrotoxin x diet interaction F (1, 34) = 15.14 p=0.0004***. Sidaks multiple comparisons, lean: aCSF vs picrotoxin p <0.0001**** and obese aCSF vs picrotoxin p= 0.6627, lean aCSF vs obese aCSF p= 0.0139*. Data are presented as mean values +/− SEM. d) Representative mIPSC recordings of lOFC pyramidal neurons of lean and obese mice. e) Diet-induced obesity (lean n=10 cells/6 mice vs. obese n=9 cells/3 mice) increased the frequency of mIPSCs onto pyramidal neurons. Unpaired t-test: t _(17)_ = 2.94, p= 0.0091**. Data are presented as mean values +/− SEM. f) Diet-induced obesity did not alter the amplitude of mIPSCs recorded from pyramidal neurons of lean (n=10 cells/6 mice) or obese (n=9 cells/3 mice) mice. Unpaired t-test: t_(17)_ = 1.754, p= 0.0975. Data are presented as mean values +/− SEM. g) Representative PPR recordings of lOFC pyramidal neurons of lean and obese mice. h) Diet induced obesity (obese=11 cells/4 mice, lean n=11 cells/4 mice) increased the ratio of P2/P1 in inhibitory paired pulse ratio. Unpaired t-test: t_(20)_ = 2.229, p=0.0375*. Data are presented as mean values +/− SEM. i) Representative tonic GABA recordings of lOFC pyramidal neurons of lean and obese mice. j) Diet induced obesity (lean n=9cells/4 mice, obese=10 cells/4mice) decreased tonic GABA holding current. Unpaired t-test: t_(17)_ = 2.348, p=0.0313*. Data are presented as mean values +/− SEM. k) Diet induced obesity (lean n=9 cells/4 mice, obese=10 cells/4mice) decreased the RMS noise. Unpaired t-test: t_(17)_ = 2.199, p=0.0420*. Data are presented as mean values +/− SEM.

Tonic inhibition is characterized by a persistent inhibitory tone that dampens excitability through distinct GABA_A_ receptors largely located extrasynaptically^26^. Tonic activation of GABA_A_ receptors can modulate neuronal excitability by increasing rheobase^27,28^. Thus, given that we observed a decrease in rheobase in obese mice, a reduction in tonic GABA may be another potential mechanism driving excitability of lOFC neurons of obese mice. To test this, we isolated sIPSCs and washed on gabazine, a selective and competitive GABA_A_ antagonist. In lean mice, application of gabazine induced a significant change in both holding current and root mean square noise, a proxy for tonically open GABA_A_ receptors^29^, consistent with previous a previous report in layer 2/3 of the frontoparietal cortex^30^. However, the tonic current was absent in lOFC neurons of obese mice (**Figure 3i-k**). Thus, diet-induced obesity reduces both synaptic and tonic inhibitory drive onto lOFC pyramidal neurons, suggesting that the enhanced excitability of lOFC pyramidal neurons is through disinhibition.

### LOFC GABAergic tone is necessary for outcome devaluation

Thus far, we have demonstrated that diet-induced obesity induces deficits in devaluation and reduces inhibitory control of lOFC principal output neurons. To causally link these synaptic changes with behaviour, we tested two hypotheses. First, that lOFC GABAergic neurotransmission is necessary for devaluation, and second, that enhancing lOFC GABAergic neurotransmission in obese mice is sufficient to restore the activity of pyramidal neurons and impairments in behavioural performance. To test our first hypothesis, we targeted lOFC inhibitory neurons using vesicular GABA transporter-cre (VGAT^cre^) mice and reduced their activity with local infusion of a cre-dependent inhibitory Designer Receptor Exclusively Activated by Designer Drug (DREADD; hM4D(Gi)), which is activated by the inert ligand clozapine n-oxide (CNO; **Figure 4a,b**). In *ex vivo* slices, we confirmed that CNO decreases the firing rate of lOFC GABAergic neurons (**Figure 4c,d**). We then examined whether disinhibition of the lOFC influenced devaluation. Lean mice expressing a control reporter in lOFC GABAergic neurons were trained to lever press (**Extended Data Figure 7a, b)** and exhibited satiety-induced devaluation in response to vehicle and CNO (**Figure 4e-g**). However, in mice expressing hM4D(Gi) in lOFC GABAergic neurons, administration of CNO impaired satiety-induced devaluation indicated by similar lever pressing in both valued and devalued states and a decrease in the revaluation index. (**Figure 4e-g**). CNO did not alter the amount of sucrose consumed during the pre-feed (**Extended Data Figure 7c**). Consistent with these effects, disinhibition of the lOFC also impaired LiCl-induced devaluation (**Figure 4h-J, Extended Data Figure 7d,e**). Taken together, these data indicate that lOFC GABAergic transmission is necessary for goal-directed behaviour.

**Figure 4:**
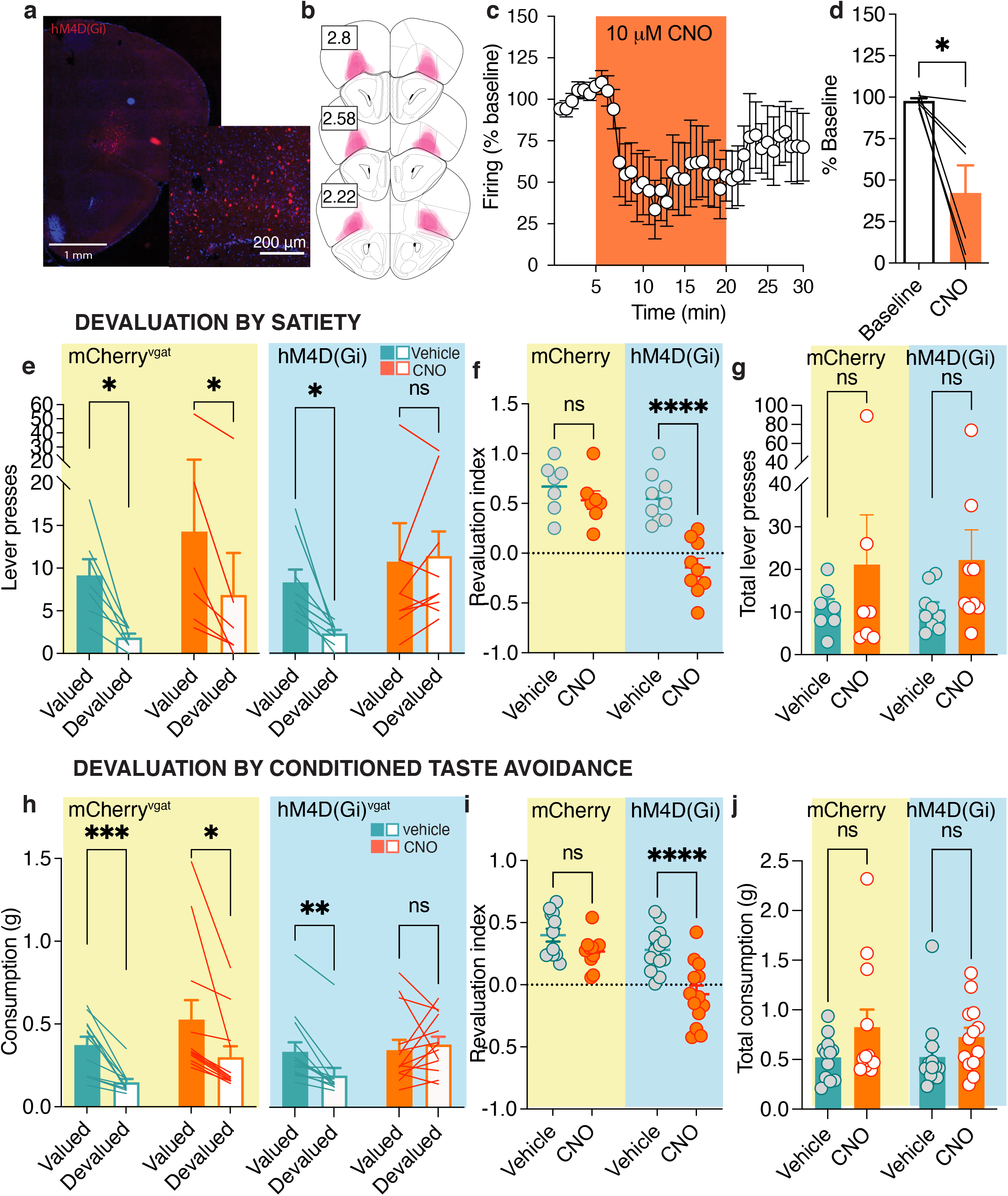
Disinhibition of the lOFC impairs devaluation. a) Representative image of hM4D(Gi) expression in the lOFC from n = 6 mice. b) Schematic representations of hM4D(Gi) expression in the lOFC. Numbers correspond to anterior distance from Bregma (mm). c) CNO application (10 μM) reduced the firing in lOFC inhibitory neurons. n = 6 cells/2 mice. Data are presented as mean values +/− SEM. d) CNO application (10 μM) reduced the firing of LOFC inhibitory neurons averaged response before and after CNO. N = 6 cells/ 2 mice. Paired t-test: t_(5)_ = 3.418, p=0.0189*. e) VGAT^cre^ mice expressing mCherry (n=7) display devaluation following vehicle and CNO (2 mg/kg, i.p.) indicated by reduced lever presses in the devalued condition. VGAT^cre^ mice expressing hM4D(Gi) (n=9) display devaluation following injection of vehicle, but not CNO. RM three-way ANOVA: CNO x virus x devaluation interaction: F (1, 14) = 2.705, p=0.1223. CNO x devaluation interaction, F (1, 14) = 2.483, p=0.1374, virus x devaluation interaction: F (1, 14) = 3.933, p=0.0673, CNO x virus interaction: F (1, 14) = 0.01563, p=0.9023. CNO effect: F (1, 14) = 3.687, p=0.0754, virus effect: F (1, 14) = 0.002476, p=0.9610, devaluation effect: F (1, 14) = 17.96, p=0.0008***. Our a priori hypothesis was that the vehicle group, but not CNO group, would devalue. Holm-Sidak’s multiple comparisons test hM4D(Gi) vehicle p=0.0153*, hM4D(Gi) CNO p= 0.7508. mCherry vehicle p=0.0153*, mCherry CNO p= 0.0153*. Data are presented as mean values +/− SEM. f) Activation of the DREADD by CNO in VGAT^cre^ mice expressing hM4D(Gi) animals (n = 9) reduced the revaluation index during devaluation compared to mCherry expressing mice (n = 7). Two-way RM ANOVA: virus (mCherry vs. hM4D(Gi) effect: F (1, 14) = 18.07, p=0.0008***, drug (vehicle vs. CNO) effect: F (1, 14) = 22.30, p=0.0003***, virus x drug interaction: F (1, 14) = 9.984, p=0.0070 **. Sidak’s multiple comparisons test: mCherry vehicle vs. CNO p=0.5311, hM4D(Gi) vehicle vs. CNO p<0.0001****. Data are presented as mean values +/− SEM. g) There were no differences in total lever presses between groups (mCherry n=7 vs. hM4D(Gi) n=9) or following vehicle or CNO administration. Two-way RM ANOVA: virus x drug Interaction: F (1, 14) = 0.01563, p=0.9023, virus effect: F (1, 14) = 0.002476, p=0.9610, drug (vehicle vs. CNO) effect, F (1, 14) = 3.687, p=0.0754. Data are presented as mean values +/− SEM. h) VGAT^cre^ mice expressing mCherry (n=12) display devaluation following vehicle and CNO (2mg/kg) indicated by decreased gelatine consumption in the devalued condition. VGAT^cre^ mice expressing hM4D(Gi) (n=13) display devaluation following injection of vehicle, but not CNO. RM three-way ANOVA: drug x devaluation interaction: F (1.000, 23.00) = 4.533, p=0.0442*, drug x virus interaction: F (1,23) = 0.3585, p=0.5552, virus x devaluation interaction: F (1,23) = 7.653, p=0.0110*, drug x virus x devaluation interaction: F (1, 23) = 4.709, p=0.0406*. CNO effect: F (1.000, 23.00) = 7.989, p=0.0096**. Virus Effect: F (1,23) = 0.1232, p=0.7288, Devaluation effect: F (1.000, 23.00) = 20.85, p=0.0001***. Holm-Sidak’s multiple comparisons test mCherry vehicle valued vs devalued: p= 0.0009*** mCherry CNO valued vs devalued p= 0.0180* hM4D(Gi) vehicle valued vs devalued: p= 0.0019 ** hM4D(Gi) CNO valued vs devalued: p= 0.5752. Data are presented as mean values +/− SEM. i) CNO administration in animals expressing hM4D(Gi) show deficits in CTA. Revaluation index following vehicle or CNO administration in animals expressing mCherry (n = 12) or hM4D(Gi) (n = 13) in CTA. RM two-way ANOVA: We observed a significant virus effect (mCherry vs. hM4D(Gi)): F (1, 23) = 15.04, p=0.0008***, significant drug (vehicle vs. CNO) effect: F (1,23) = 26.44, p<0.0001***, and a virus x vehicle vs. CNO interaction: F (1,23) = 5.700, p=0.0256*. In Sidak’s multiple comparisons test, we only observed a significant effect (vehicle vs. CNO) in animals expressing hM4D(Gi) p<0.0001****. Data are presented as mean values +/− SEM. j) Total consumption during the CTA test (post-diet) in mCherry (n = 12) and hM4D(Gi) mice (n = 13) following either vehicle or CNO. Two-way RM ANOVA: virus x drug (vehicle or CNO) interaction: F (1,23) = 0.3585, p = 0.5552, virus (mCherry vs. hM4D(Gi)) effect: F (1,23) = 0.1232 p=0.7288, and drug (vehicle vs. CNO) effect: F (1,23) = 7.989 p=0.0096. Data are presented as mean values +/− SEM.

We then employed two strategies to test our second hypothesis, that enhancing lOFC GABAergic neurotransmission in obese mice would restore the activity of pyramidal neurons and behavioural performance. In the first strategy, we used a pharmacological approach to enhance lOFC GABAergic neurotransmission by administering the selective GAT-1 GABAergic transporter inhibitor, NNC-711, to the lOFC. Obese mice display increased excitability and decreased inhibitory transmission at OFC pyramidal neurons. We found that application of NNC-711 (10 μM) to lOFC slices restored excitability levels of pyramidal neurons in obese mice without significantly altering those of lean mice (**Figure 5a-c**). NNC-711 had no effect on the frequency, nor the amplitude of sIPSCs (**Figure 5d-f**), but reduced the amplitude of evoked IPSCs in lean but not obese mice (**Figure 5g-h**), again suggesting that obese mice have reduced GABA release pools. Finally, we tested if NNC-711 could restore tonic GABA currents of obese mice. Application of NNC-711 increased tonic GABA, evident by an increase in gabazine-induced change in holding current, and an increase in RMS noise in lean but not obese mice (**Figure 5J-I**). Taken together, blocking GABA reuptake decreases excitability of pyramidal neurons and increases tonic GABA current in obese mice.

**Figure 5.**
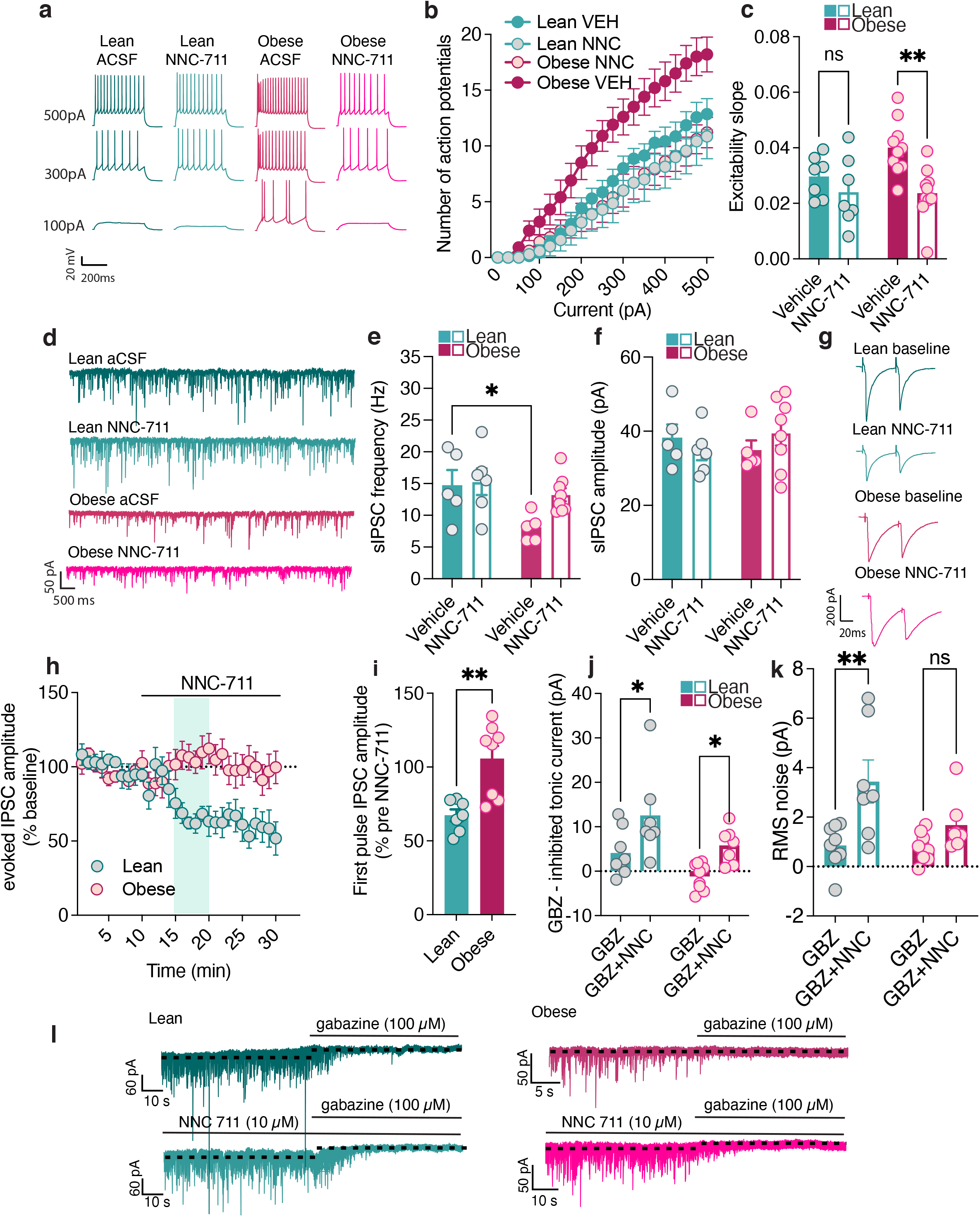
A GAT-1 transporter inhibitor reverses diet-induced impairment in inhibitory tone of obese mice. a) Representative recordings of action potentials observed at 100pA, 300pA and 500pA current steps from lOFC pyramidal neurons of lean and obese mice with ACSF or NNC-711 (a GAT-1 GABA reuptake inhibitor). b) Diet-induced obesity increased the excitability of lOFC pyramidal neurons as indicated by frequency-current (F-I) plots of action potential numbers at current injections from 0pA to 500pA of lean + ACSF (n=7 cells/4 mice), lean + NNC-711 (n=7cells/4 mice), obese + ACSF (n=10 cells/4 mice) and obese + NNC-711(n=10 cells/6 mice). RM 3-way ANOVA mixed effects analysis: current effect: F (2.154, 34.46) = 181.3 p<0.0001****, drug effect: F (1.000, 16.00) = 7.076 p= 0.0171*, Diet effect: F (1,278) = 3.272, p= 0.0715, current x drug interaction: F (1.192, 16.56) = 7.192, p= 0.0127*, Current x diet interaction: F (20, 278) = 1.816, p= 0.0190*, Drug x diet interaction F (1, 278) = 2.438, p= 0.1196 and current x drug x interaction: F (20, 278) = 1.756, p=0.0253*. Data are presented as mean values +/− SEM. c) Excitability slope of pyramidal neurons from the F-I plot of pyramidal neuronal activity from of lean + ACSF (n=7 cells/4 mice), lean + NNC-711 (n=7cells/4 mice), obese + ACSF (n=10 cells/4 mice) and obese + NNC-711(n=10 cells/6 mice). NNC-711 (10 μM) significantly reduced the firing of lOFC pyramidal neurons from obese mice compared to vehicle + obese. Two-way RM ANOVA mixed-effects model: drug effect: F (1,30) = 10.32, p= 0.0031**, Diet effect: F (1,30) = 2.126, p = 0.1552, diet x drug interaction: F (1, 30) = 2.447, p=0.1282. Sidaks multiple comparisons test: Lean: vehicle vs NNC-711 p=0.4977 and obese: vehicle vs. NNC-711 p= 0.0016**. Data are presented as mean values +/− SEM. d) Representative sIPSC recordings of lOFC pyramidal neurons of lean and obese mice with ACSF or NNC-711. e) Diet-induced obesity decreased the frequency of sIPSCs onto pyramidal neurons. lean + ACSF (n=5 cells/4 mice), lean + NNC-711 (n=6 cells/6 mice), obese + ACSF (n=5 cells/2 mice) and obese + NNC-711(n=8 cells/5 mice). Two-way RM ANOVA: drug effect: F (1, 20) = 2.944, p=0.1016, Diet effect: F (1, 20) = 6.743, p=0.0172*, diet x drug interaction: F (1, 20) = 1.898, p=0.1836. Based on previous data, we had an a priori hypothesis that diet would influence frequency of sIPSC. Sidaks multiple comparisons test: vehicle lean vs obese p= 0.0330*. Data are presented as mean values +/− SEM. f) Diet did not alter amplitude of sIPSCs onto pyramidal neurons. Lean + ACSF (n=5 cells/4 mice), lean + NNC-711 (n=6 cells/6 mice), obese + ACSF (n=5 cells/2 mice) and obese + NNC-711(n=8 cells/5 mice). Two-way RM ANOVA: NNC-711 effect: F (1,20) = 0.02718, p=0.8707, Diet effect: F (1, 20) = 0.05205, p=0.8219, Diet x Drug interaction: F (1, 20) = 1.602, p=0.2202. Data are presented as mean values +/− SEM. g) Representative evoked IPSC recordings of lOFC pyramidal neurons of lean and obese mice with ACSF or NNC-711. h) NNC-711(10 μM) application reduced eIPSC amplitude in lean (n = 7 cells/6 mice) but not obese mice (n=8 cells/6 mice). Shaded bar represents section analyzed for bar graph in i. Data are presented as mean values +/− SEM. i) NNC-711 produced a significant more decrease in amplitude in lean (n = 7 cells/6 mice) compared to obese mice (n=8 cells/6 mice). Unpaired t-test: t _(13)_ = 3.786, p= 0.002**. Data are presented as mean values +/− SEM. j) Tonic GABA current increased in the presence of NNC-711. Lean + ACSF (n=8 cells/4 mice), lean + NNC-711 (n=7 cells/4 mice), obese + ACSF (n=9 cells/5 mice) and obese + NNC-711(n=7 cells/4 mice). Two-way RM ANOVA: NNC-711 effect: F (1, 27) = 12.90, p=0.0013**, Diet effect: F (1,27) = 7.786, p=0.0095**, Diet x Drug interaction: F (1,27) = 0.1367, p=0.7144. Holms-Sidaks multiple comparisons test lean: p=0.0202*, Obese: p= 0.0288*. Data are presented as mean values +/− SEM. k) RMS noise in the presence of NNC-71 increased in lean but not obese mice. Lean + ACSF (n=8 cells/4 mice), lean + NNC-711 (n=7 cells/4 mice), obese + ACSF (n=9 cells/5 mice) and obese + NNC-711(n=7 cells/4 mice). Two-way RM ANOVA: NNC-711 effect: F (1,27) = 13.34, p=0.0011**, Diet effect: F (1, 27) = 3.721, p=0.0643, Diet x Drug interaction: F (1, 27) =3.095, p=0.0898. Holms-Sidaks multiple comparisons test lean: p=0.0016**, Obese: p=0.1860. Data are presented as mean values +/− SEM. l) Representative traces of tonic GABA current in lean and obese mice in the presence of NNC-711 and or gabazine.

We then assessed if enhancing lOFC GABAergic neurotransmission reinstates devaluation in obese mice, by infusing NNC-711 locally in the lOFC (**Figure 6a,b**). Lean and obese mice were trained to lever press for sucrose and then were pre-sated with sucrose immediately prior to the test (**Extended data Figure 8a-c)**. Lean mice devalued sucrose when either vehicle or NNC-711 was microinfused into the lOFC, evident by decreased lever pressing and a positive revaluation index (**Figure 6c,d**). Notably, there was an increase in proportion lean mice devaluing to the expected outcome after NNC-711 (veh: 66% mice devalued, NNC-711: 92% mice devalued). In contrast, obese mice receiving lOFC vehicle infusions failed to display devaluation, while intra-lOFC NNC-711 prior to testing restored devaluation (**Figure 6c,d**). There were no differences in the total number of lever presses between groups (**Figure 6e**) and NNC-711 did not alter locomotor activity (**Extended Data Figure 8e)**.

**Figure 6.**
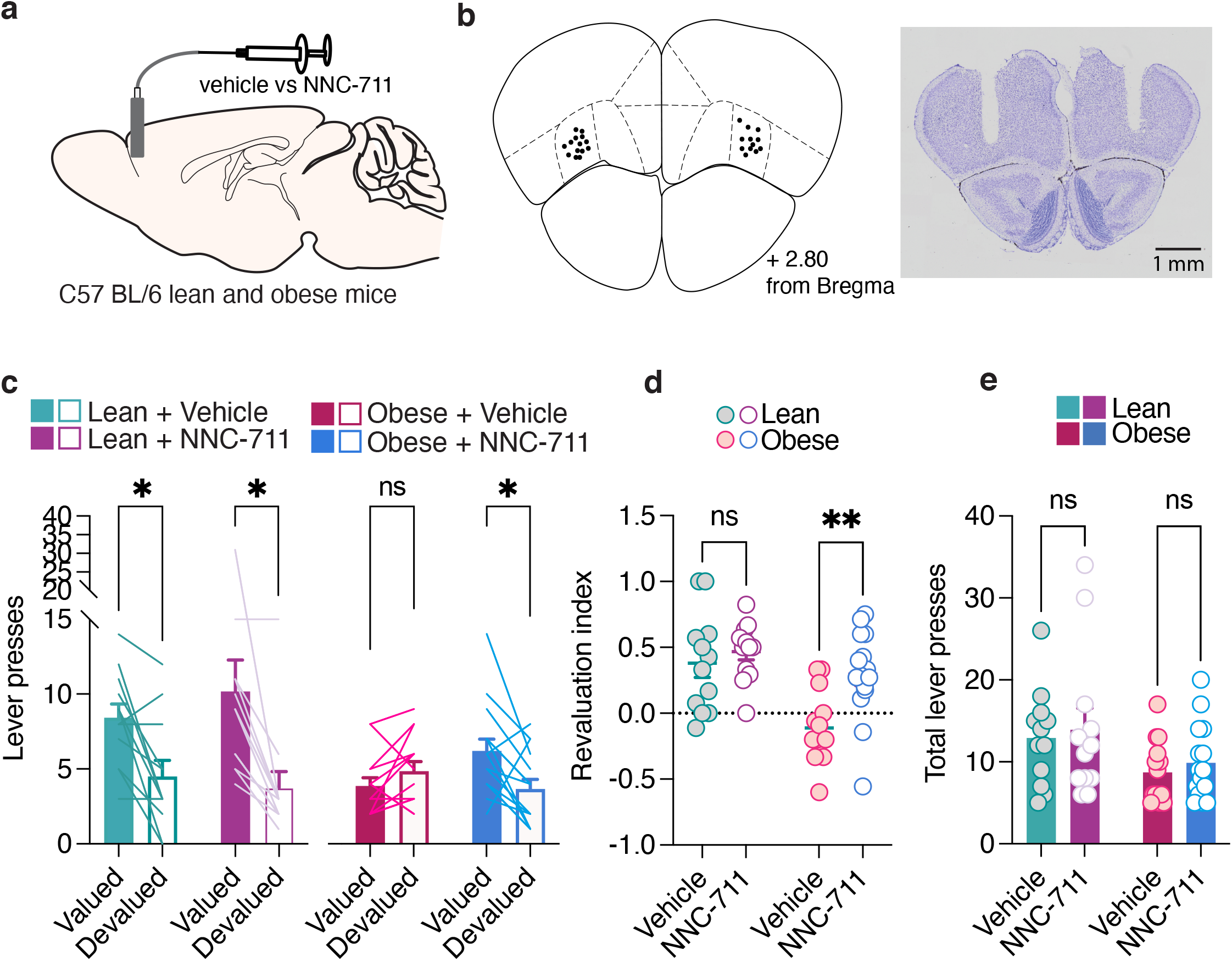
Increasing GABAergic tone in the lOFC of obese mice rescues goal-directed behaviour. a) Strategy for NNC-711 administration to the lOFC to increase inhibitory neurotransmission. b) Cannulae placements and representative Nissl staining of an lOFC slice. c) Lever presses of lean vehicle mice (n=12), lean NNC mice (n=12), and obese vehicle mice (n=14) and obese NNC mice (n=15) in the valued (closed bars) and devalued (open bars) conditions following bilateral lOFC infusions of either vehicle or NNC-711. Three-way ANOVA mixed effects model: Devaluation and diet Interaction: F (1, 14) = 14.34, p= 0.0020**, Devaluation and drug interaction: F (1.000, 14.00) = 6.739, p= 0.0211*, Diet and drug interaction: F (1, 14) = 0.008637, p= 0.9273, Devaluation and diet and drug interaction: F (1, 14) = 0.1977, p= 0.6634: Devaluation main effect F (1.000, 28.00) = 26.07, p= <0.0001****, Diet main effect: F (1, 14) = 4.241, p= 0.0586, Drug main effect F (1.000, 28.00) = 0.9240, p=0.3446. Our a priori hypothesis was lean but not obese mice would devalue. Holm-Sidak’s multiple comparison’s: Lean vehicle valued vs devalued p= 0.0142*, Lean NNC-711 valued vs devalued p= 0.0204*, Obese vehicle valued vs devalued p= 0.1309, obese NNC valued vs devalued p= 0.0201*. Data are presented as mean values +/− SEM. d) Revaluation index of lean vehicle mice (n=12), lean NNC-711 mice (n=12), obese vehicle mice (n=14) and obese NNC-711 mice (n=15) after lOFC infusion of either vehicle or NNC-711. Twoway RM ANOVA mixed effects model: diet effect: F (1,28) = 14.23, p= 0.0008***, NNC-711 effect: F (1, 21) = 10.02, p= 0.0047**, Diet and drug Interaction: F (1, 21) = 4.074 p=0.0565. Sidak’s multiple comparisons test: lean vehicle vs NNC-711: p=0.6948, obese vehicle vs NNC-711: p=0.0018**. Data are presented as mean values +/− SEM. e) Total number of lever presses of lean vehicle mice (n=12), lean NNC-711 mice (n=12), obese vehicle mice (n=14) and obese NNC-711 mice (n=15) mice during the devaluation task following bilateral lOFC infusions of either vehicle or NNC-711. Two-way RM ANOVA mixed effects model: diet x drug interaction: F (1,21) = 0.01049, p = 0.9194, drug (vehicle vs. NNC-711) effect: F (1, 21) = 1.074, p=0.3119, diet effect: F (1, 28) = 4.262, p=0.0484*. Data are presented as mean values +/− SEM.

The second strategy for enhancing GABAergic neurotransmission in the lOFC involved optogenetic activation of lOFC local inhibitory neurons. To test this, we expressed channelrhodopsin2 (ChR2) in GABAergic interneurons in the lOFC of lean and obese VGAT^cre^ mice. We confirmed that lOFC GABAergic interneurons expressing ChR2 reliably responded to blue light (**Extended Data Figure 9a,b)**. We then tested if optogenetic activation of inhibitory neurons in lOFC could restore excitability levels of pyramidal neurons in obese mice. As observed previously, lOFC pyramidal neurons from obese mice had increased excitability compared to those of lean mice (**Figure 7a-c**). Photostimulation of GABAergic inputs (5 x 1s 5Hz pulse trains at 4 mW) prior to the current-steps reduced the excitability of lOFC pyramidal neurons in these mice (**Figure 7a-c**). Photostimulation also decreased the excitability of pyramidal cells in lean mice, suggesting that although there are higher levels of baseline inhibition in lean mice, GABAergic inputs can be further engaged to reduce excitability. In line with our earlier data, obese mice had decreased frequency but not amplitude of sIPSCs, but the optical stimulation protocol had no effect on sIPSCs in lean or obese mice (**Figure 7d-f)**. To assess if the low frequency optical stimulation protocol altered evoked GABA release, we optically evoked inhibitory postsynaptic currents. Prior to the low frequency stimulation protocol, obese mice showed a paired pulse facilitation compared to lean mice (**Figure 7g,h**). 5 min after low frequency optical stimulation, the PPR was increased in lean mice, but not obese mice (**Figure 7g,h**). Optical stimulation reduced the first pulse amplitude in lOFC neurons of both lean and obese mice (**Figure 7i**), suggesting this protocol releases GABAergic vesicle stores. Taken together, low frequency activation of GABAergic interneurons decreases excitability of pyramidal neurons and releases GABAergic stores, which may underlie the decreased excitability after optical stimulation.

**Figure 7:**
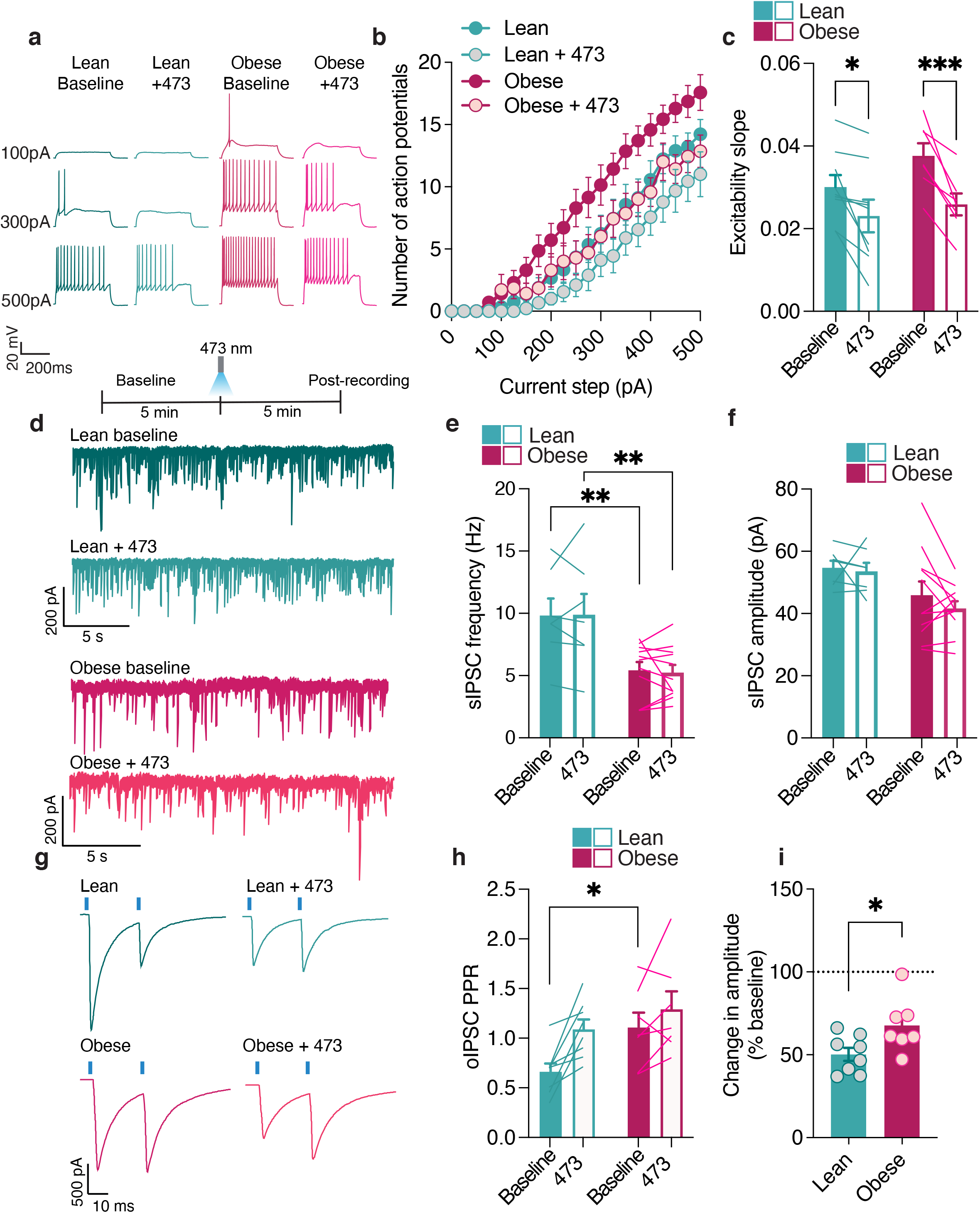
Increasing GABAergic drive via optogenetic stimulation of GABAergic interneurons restores GABAergic inhibition of lOFC pyramidal neurons. a) Representative recordings of action potentials observed at 100pA, 300pA and 500pA current steps from lOFC pyramidal neurons of lean and obese VGAT mice after a 5Hz optogenetic protocol. b) Diet-induced obesity increased the excitability of lOFC pyramidal neurons as indicated by frequency-current (F-I) plots of action potentials at current injections from 0pA to 500pA of lean (n=9 cells/4 mice), lean + 473 (n=9 cells/4 mice), obese (n=7 cells/4 mice) and obese + 473 (n=7 cells/4 mice). RM 3-way ANOVA: current step effect F (4.154, 58.16) = 102.5, p<0.0001****, stimulation effect: F (1.000, 14.00) = 19.88, p=0.0005***, Diet effect: F (1, 14) = 3.954, p=0.0667, current step x stimulation: F (2.535, 35.49) = 10.52, p<0.0001****, Current step x diet interaction: F (20, 280) = 1.505, p=0.0784, stimulation x diet interaction F (1, 14) = 1.011, p=0.3317, Current x Drug x Diet interaction: F (20, 280) = 0.9284, p=0.5513. Data are presented as mean values +/− SEM. c) Excitability slope of pyramidal neurons from the F-I plot of pyramidal neuronal activity from lean (n=9 cells/4 mice), lean + 473 (n=9 cells/4 mice), obese (n=7 cells/4 mice) and obese + 473 (n=7 cells/4 mice). Two-way RM ANOVA: Stim effect: F (1, 14) = 33.58, p<0.0001****. Diet effect: F (1, 14) = 1.396, p=0.2571, Diet x stim interaction: F (1, 14) = 2.141, p=0.1655. Sidaks multiple comparisons test: lean pre vs post stimulation p= 0.0110*, obese pre and post stimulation p= 0.0005***. Data are presented as mean values +/− SEM. d) Representative sIPSC recordings of lean and obese prior and post optogenetic photo stimulation protocol. e) Diet-induced obesity decreased the frequency of sIPSCs onto pyramidal neurons in obese mice (lean n=7 cells/4 mice, obese n=11 cells/5 mice). In both diets, optogenetic stimulation did not alter sIPSC frequency. Two-way RM ANOVA: Stim effect: F (1, 16) = 0.01998, p=0.8894. Diet effect: F (1, 16) = 10.66, p=0.0049**, Diet x stim interaction: F (1, 16) = 0.09190, p=0.7657. We had an a priori hypothesis that diet would alter frequency, therefore we did a Sidaks multiple comparisons test: baseline lean vs obese p= 0.0095**, post stimulation lean vs obese p=0.0060**. Data are presented as mean values +/− SEM. f) Stimulation protocol did not alter amplitude of sIPSCs onto pyramidal neurons in lean (n=7 cells/4 mice) or obese mice (n=11 cells/5 mice), Two-way RM ANOVA: Stimulation effect: F (1, 16) = 1.291, p=0.2726. F (1, 16) = 5.911, p=0.0272*, Diet x stimulation interaction: F (1, 16) = 0.4151, p=0.5285. Data are presented as mean values +/− SEM. g) Representative oPPR recordings of lOFC pyramidal neurons of lean and obese mice. h) Diet induced obesity (lean=8 cells/6 mice, obese n=7 cells/4 mice) increased the ratio of P2/P1 in optically evoked paired pulse ratio prior to optogenetic stimulation. Two-way RM ANOVA: Stim effect: F (1, 13) = 14.12, p=0.0024**, Diet effect: F (1, 13) = 3.819, p=0.0725, Diet x stimulation interaction: F (1, 13) = 2.211, p=0.1609. We had an a priori hypothesis that diet would alter oPPR. Sidaks multiple comparison of lean and obese mice prior to optogenetic photo stimulation p= 0.0462*. Data are presented as mean values +/− SEM. i) Optical stimulation (lean=8 cells/6 mice, obese n=7 cells/4 mice) decreased the amplitude of the first pulse in lean and obese mice. Unpaired t-test: t_(13)_ = 2.438, p=0.0299*. Data are presented as mean values +/− SEM.

Lastly, we accessed if optically enhancing lOFC GABAergic neurotransmission reinstates devaluation in obese mice. VGAT^cre^ mice were placed on a low or high fat diet and trained to lever press on an RR20 schedule (**Extended Data Figure 9c,d**). ChR2 was expressed in GABAergic interneurons and a dual fibre-optic cannula was implanted in layer 2/3 of the lOFC (**Figure 8a-c**). Lean and obese mice were prefed with sucrose (**Extended Data Figure 9e**) and then received either 589 or 473 nm optical stimulation (5x 1s 5Hz pulses) immediately prior to the test for satiety-induced devaluation. Lean mice displayed devaluation in the presence of non-ChR2 activating light (589 nM) or the ChR2 activating wavelength (473 nM; **Figure 8d,e**). In response to the non-activating wavelength, obese mice were insensitive to devaluation (**Figure 8d,e**). However, activation of lOFC inhibitory neurons rescued devaluation in obese mice evident by decreased lever pressing in the devalued state and an increase in the revaluation index (**Figure 8d,e**). Photostimulation did not alter total lever presses or locomotor activity in lean or obese mice (**Figure 8f, Extended data Figure 9f**). Taken together, restoring lOFC pyramidal neuron firing activity by either boosting GABAergic firing with optogenetics or GABAergic tone via a GAT-1 blocker can restore devaluation in obese mice.

**Figure 8:**
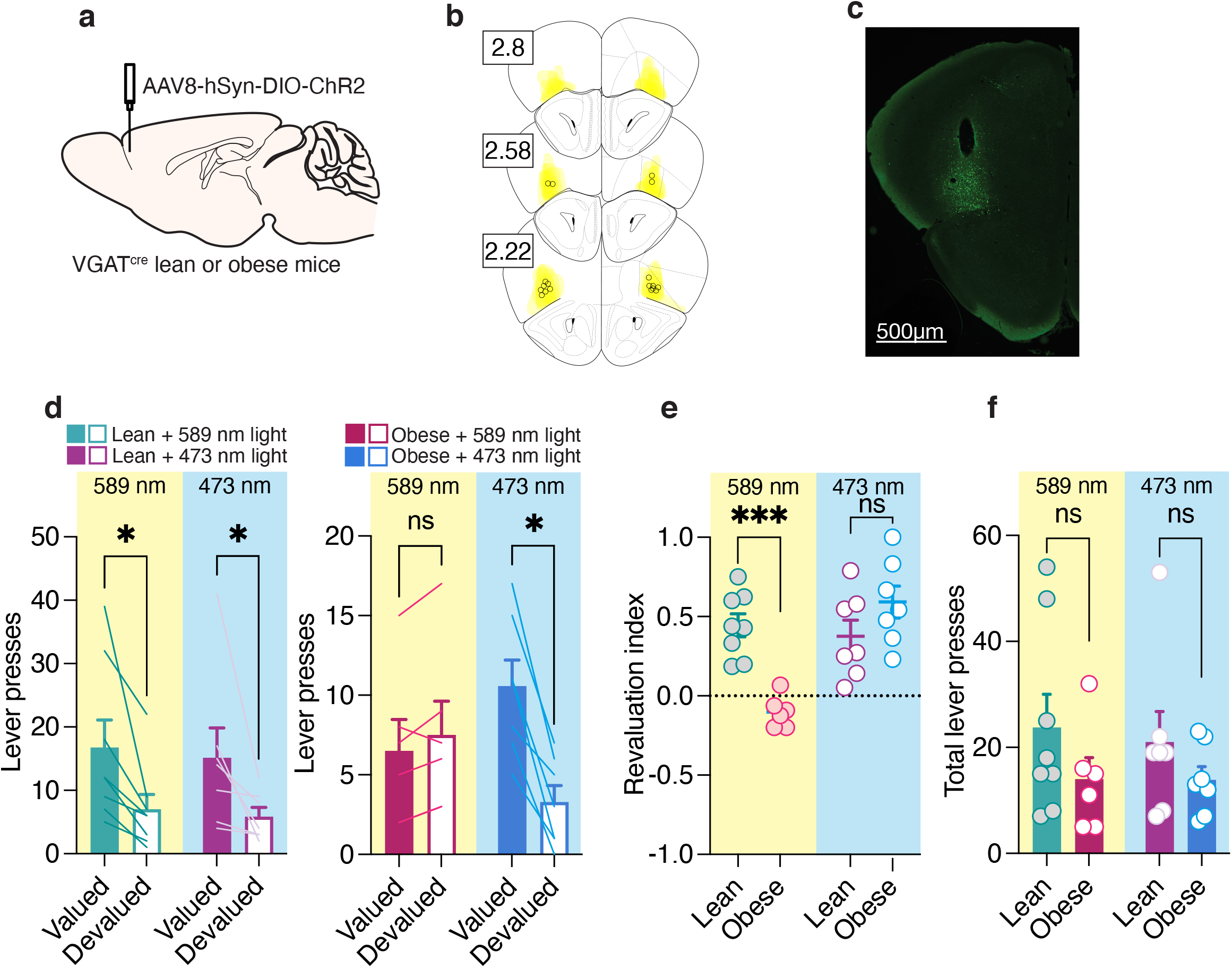
Optogenetically restoring inhibitory drive in the lOFC of obese mice rescues goal-directed behaviour. a) Viral strategy for optogenetic enhancement of lOFC inhibitory neurotransmission. b) Schematic representations of ChR2 expression in the lOFC for devaluation experiments. Numbers correspond to anterior distance from Bregma (mm). c) Representative image AAV2-EF1a-DIO-hChR2-EYFP expression (green) and optic fibre implantation site in the lOFC of VGAT^cre^ mice from n = 6 mice. d) Photo stimulation in lean mice (589nM (n = 8) vs. 473nM (n = 7)) did not alter devaluation. Lever presses in the valued or devalued state of obese mice following 589 nm (inactive, n = 6) and 473nm (active, n = 7) photostimulation. Three-way ANOVA mixed effects model: devaluation effect F(1,13)=21.95, p=0.0004***, photostimulation effect: F(1,13)=0.3920 p=0.5421 or diet effect: (F(1,9)=1.594, p=0.2348, devaluation x photo stimulation interaction: F(1,9)=2.094, p=0.1818, devaluation x diet interaction: F(1,9)=5.564, p=0.0427*, diet x photo stimulation interaction: (F(1,9)=0.4254, p=0.5306), devaluation x diet x photo stimulation interaction: (F(1,9)=2.621, p=0.1399). A Holm-Sidak’s multiple comparison test indicated that lean animals devalued at either photo stimulation wavelength 589nM: p = 0.0149*, 473nM: p = 0.0215*, obese mice devalued at 473nM p = 0.0474* but not 589nm p = 0.7385. Data are presented as mean values +/− SEM. e) During inactive light (lean 589 nM n=8, obese 589 nM n=6), significant group differences in revaluation index were observed with lean mice displaying a positive revaluation index and mice with obesity displaying a negative one. During active light (473 nM lean n=7, 473 nM obese n=7), the revaluation index of both diet groups was positive, and no group difference was observed. Two-way RM ANOVA mixed effects model: diet x stimulation interaction: F (1, 24) = 20.29, p=0.0001***, diet effect: F (1,24) = 3.829, p = 0.0621, photo stimulation effect: F (1,24) = 13.61, p = 0.0012**. Sidak’s multiple comparison test: 589 nm (inactive) light lean vs. obese p=0.0003***; 473nm (active) light lean vs. obese p=0.1586. Data are presented as mean values +/− SEM. f) Inactive light (589 nM lean n=8, 589 nM obese n=6) vs. active light (473 nm) did not alter the total number of lever presses performed during the devaluation test in lean or obese mice. Two-way RM ANOVA: Interaction: F (1,11) = 0.4254, p = 0.5277, photostimulation effect (589nm vs. 473nm): F (1, 11) = 0.3920, p=0.5440, diet effect: F (1, 13) = 1.594, p=0.2289. Data are presented as mean values +/− SEM.

## Discussion

These data demonstrate a causal role of lOFC GABAergic transmission in obesity-induced impairment in devaluation. Long-term exposure to a high-fat diet led to the development of diet-induced obesity and subsequent disruption of goal-directed behaviour in three different devaluation paradigms. In obese mice, GABAergic release probability as well as GABAergic tone onto pyramidal neurons was decreased, leading to hyperexcitability of principal output neurons. Relatedly, reducing GABAergic neurotransmission in lean mice disrupted goal-directed behaviour, further suggesting that lOFC GABAergic neurotransmission is necessary for devaluation induced by satiety and conditioned taste avoidance. Finally, increasing GABAergic neurotransmission in obese mice, by pharmacological or optogenetic methods, restored lOFC pyramidal neuron excitability and devaluation. Together, our results suggest that in obesity, lOFC dependent impairments in devaluation may change the ability to use the value of the outcome to guide behaviour, thus making one’s behaviour less flexible to changes in satiety.

### Obesity disinhibits principal output neurons

We observed neurophysiological changes in the lOFC of obese mice. The probability of GABAergic release onto lOFC pyramidal neurons was reduced in obese mice, consistent with previous work showing that obese rats given 24h access to a cafeteria diet had reduced GABAergic release probability onto layer II/III pyramidal neurons in the lOFC^22^ through an endocannabinoid mechanism^21^. In our recordings, obese mice from 3 different cohorts and 2 different genetic lines also demonstrated enhanced excitability of pyramidal neurons. We propose that the enhanced excitability of lOFC pyramidal neurons is due to disinhibition rather than a change in intrinsic excitability or altered excitatory inputs. While increased excitatory input could drive firing, our previous work demonstrated that extrasynaptic glutamate suppresses glutamate release via presynaptic mGluR2/3 receptors in obese rats^21^, which would be unlikely to increase firing rate. Surprisingly, we did not observe a change in RMP of obese mice, as found previously^22^. It is possible that our recordings sample from different populations of pyramidal neurons^31^ and that any obesity induced changes in RMP are washed out when recording from the total population. Other changes in intrinsic properties, such as decreased rheobase and AHP width in the lOFC of obese mice, were absent in the presence of the GABA_A_ inhibitor, picrotoxin, suggesting that these changes were secondary to inhibitory signaling. Furthermore, picrotoxin also significantly increased the excitability of lOFC neurons of lean mice, without altering that of obese mice, suggesting that GABAergic mechanisms controlling the excitability of lOFC pyramidal neurons are suppressed in obese mice. Consistent with this, we found that tonic GABA observed in lean mice was absent in obese mice. This suggests a reduction of tonic inhibitory control of OFC pyramidal neurons during obesity, which could underlie the increased firing observed in obese mice^32^. Previous work has shown that tonic inhibition primarily modulates excitability by increasing rheobase^27,28^. Notably, decreased rheobase was associated with the absence of tonic GABA in obese mice. In addition to reduced tonic GABA currents, we observed a reduction in synaptic GABAergic release probability in obese mice, demonstrated by decreased mIPSC frequency and paired pulse facilitation of evoked IPSCs. Finally, increasing GABAergic release by either using a GAT-1 blocker or optogenetically activating lOFC GABAergic neurons decreased the excitability of lOFC pyramidal neurons in obese mice. GABAergic interneurons form axo-somatic synapses, and are well positioned to coordinate principal output neuronal firing^25,33^. Thus, disruption of synaptic and tonic GABAergic release onto pyramidal neurons in obese mice, likely dysregulates the coordinated firing of principal neurons, ultimately leading to altered behavioural performance. Taken together, our data indicate that diet-induced obesity disinhibits lOFC neurons and this effect can occur across different strains of mice, species, and diet types.

### Diet-induced obesity disrupts goal-directed behaviour

We demonstrate that diet-induced obesity disrupts goal-directed behaviour in three different devaluation tasks. Two of these tasks, satiety and conditioned taste avoidance, involve devaluing the outcome whereas the third task, contingency change, involves devaluing the action to obtain the outcome. While impairment on satiety-induced devaluation tasks has been previously reported in obese rodents^9^, and humans^8^, it was unclear if this was due to a failure in associative learning, altered satiety processing, or inflexible behaviour.

Although there was no significant difference in total lever presses between lean and obese mice during the test, obese mice consistently made fewer responses throughout training. Previous work has also noted reduced instrumental actions of obese rodents^34–36^. The reduction in instrumental responding may be attributed to a downshift in the expected value of the reward compared to their home cage diet^34^. We tried to mitigate this effect by using a higher sucrose concentration during instrumental training, 30% compared to the 9% sucrose present in the high fat diet. An alternative explanation for decreased instrumental responding of obese mice may be due to general attenuation of locomotor and exploratory behaviour, as previously observed in obese mice^37^. Consistent with this, we observed an overall decrease in locomotion in obese compared to lean mice, congruent with other studies^37–39^.

Another interpretation of reduced lever pressing and failure to devalue in obese mice is that they are unable to engage in instrumental learning and performance. However, both lean and obese mice increase lever presses when the response requirement ratio increases, such that there are significant increases in lever pressing at RR10 and RR20 compared to RR5. Secondly, the breakpoint in a progressive ratio task was not different between lean and obese mice, suggesting that both groups learned to escalate their responses for the reward and mice were in similar motivational states. Thirdly, when lever presses during last day of training were matched, we observed a significant devaluation in lean, but not obese mice, suggesting that impairment in devaluation in obese mice is not due to underlying differences in the strength of learning. Finally, devaluation was restored in obese mice with enhanced lOFC GABAergic tone, suggesting that these associations had been initially made. Therefore, the inability of obese mice to devalue is unlikely due to a failure in associative learning.

An alternative explanation for impaired devaluation in obese mice is that they have different sensitivity to sensory feedback and/or satiety processing. Indeed, obesity-prone rats have decreased taste sensitivity compared to obesity-resistant rats^40^. Giving lean mice a sucrose-reinforced devaluation test, to remind them of the sensory or satiety properties of the outcome, was sufficient to induce devaluation. However, obese mice continued to press for sucrose in the devalued state, suggesting that either they are insensitive to the sensory properties of the outcome or that they have impaired goal-directed behaviour. To further dissociate this, we used a flavour preconditioning experiment where mice were pre-fed with an isocaloric cherry-flavoured sucrose solution or unflavoured sucrose solution prior to the devaluation test. Lean mice devalued to the preconditioned flavour, whereas obese mice did not, suggesting that general satiety is not differentially influencing the task. Consistent with this, both groups consumed less of the preconditioned (devalued) flavour compared to the valued flavour in the post-test, suggesting that both groups were sated during the preconditioning period. Furthermore, because mice consumed similar amounts in the preconditioning period, it suggests that both lean and obese mice can discriminate between flavours and value them equally, and that the water restriction period did not differentially influence their motivational state. Thus, the impairment in devaluation in obese mice is not due to differences in satiety processing.

We used an instrumental procedure to assess whether disinhibition of the lOFC influences satiety-induced devaluation of lean mice. The literature on OFC involvement in instrumental devaluation is somewhat contradictory. For example, excitotoxic lesions of the lOFC in rats did not affect instrumental performance on an outcome devaluation task, whereas these lesions disrupted Pavlovian outcome expectancies, such that rats were unable to update their performance in response to a changed Pavlovian contingency^41,42^. However, other studies using inhibitory DREADDs have demonstrated inhibition of the lOFC or the vlOFC impairs actionoutcome associations when instrumental contingencies have already been learned or have been reversed^11,12,43^. It is possible that disinhibition of the lOFC reflects an impairment in generalizing the devaluation setting and test. However, when obese mice were pre-fed within the same setting as the devaluation test, they did not devalue the expected outcome in comparison to lean mice with fully functioning OFCs. We also observed that when the lever contingency changed, lean mice learned the new contingency, whereas obese mice did not update their actions, consistent with impairment in vlOFC function^43^. Therefore, while lean mice can encode and recall the identity of the expected outcome, obese mice are impaired in representing, updating, or using the response-outcome associations. Taken together, the inability of obese mice to devalue rewards is not due to altered motivational state, disrupted satiety, or poor associative learning, but rather is most likely due to the inability use the value of the outcome to guide behaviour.

Our results support the hypothesis that devaluation requires sufficient inhibitory tone in the lOFC. Using chemogenetics, inhibition of lOFC GABAergic neurons of lean mice disrupts both satiety-induced devaluation as well as devaluation by conditioned taste avoidance. Importantly, inactivation of the OFC does not alter the palatability of food rewards^16^ or the amount of sucrose consumed during the prefeed, suggesting that the impairment in devaluation is likely related to altered goal-directed behaviour. Indeed, parvalbumin-containing (PV+) interneurons in the OFC facilitate cognitive flexibility as mice with reduced PV+ expression have impaired reversal learning^44^. Diet can influence perineuronal net expression around PV+ interneurons of the OFC^45^, and this could potentially influence their synaptic transmission^46^. Therefore, PV+ interneurons are a likely target for obesity-induced changes in OFC function and future work should investigate the interneuron subtype and mechanisms of altered synaptic transmission influenced by obesity.

### Augmenting GABA tone restores devaluation in obese mice

Given that intact lOFC GABAergic function is required for devaluation and boosting GABAergic function can restore the appropriate firing rate of pyramidal neurons of obese mice, we hypothesized that we could restore goal-directed behaviour in obese mice by increasing GABAergic function. Indeed, we found that enhancement of GABAergic tone with a GAT-1 reuptake inhibitor restores devaluation in obese mice and modestly improves that of lean mice. Similarly, optogenetic activation of GABAergic neurons restores devaluation in obese mice. These results are not due to altered locomotor activity as neither intra-OFC NNC-711 nor optogenetic stimulation of GABAergic neurons influenced distance travelled in an open field. Interestingly, optogenetic stimulation of lOFC GABAergic neurons was done prior to behavioural testing. The stimulation protocol mobilized vesicular GABAergic release demonstrated by a decrease in amplitude of oIPSCs compared to baseline lasting throughout the 10 min test session. Furthermore, we propose that NNC-711 effects on increasing tonic GABA currents could underlie reduced disinhibition in the lOFC of obese mice leading to a restoration of goal-directed behaviour. It is possible that the modest changes in behaviour observed in lean mice are because they already have appropriate tonic GABA control of pyramidal neuronal excitability and NNC-711-induced changes in tonic GABA minimally reduced the excitability, whereas the effects of NNC-711 are more profound in obese mice as it restores the influence of tonic GABA on the excitability of lOFC neurons. Given that decreased OFC function^11,12^ and obesogenic diet manipulations^47^ can promote habit formation, obesity-induced alterations in lOFC neurophysiology may underlie increased habits and increasing GABAergic tone in the lOFC restores goal-directed control.

Taken together, we causally demonstrate that GABAergic synaptic transmission in the lOFC underlies devaluation, and that increasing GABAergic tone in the lOFC of obese mice restores goal-directed behaviour. These data propose that obesity-induced changes in lOFC function impedes one’s ability to update actions based on current information and suggests that obesity-induced perturbations in lOFC functioning may be an underlying mechanism that contributes to habit-like behaviour, which presents an additional challenge for those maintaining food restrictive diets. Sex differences in central responses to obesogenic diets are well established^48^. Given that male mice were used in this study, it will be important to establish if sex differences in OFC function in obesity in future studies. Currently, there are five approved drug therapies for long-term weight management and only two have demonstrated minimal weight loss efficacy. Thus, the development of new therapies is of critical importance and our findings that inhibitory drive in the orbital regions of the frontal lobes impacts reward processes provide a novel putative target for potential therapeutics.

## Acknowledgements

The authors would like to acknowledge the Hotchkiss Brain Institute optogenetic core facility and the advanced microscopy facility for their technical support. This research was performed at the University of Calgary which is located on the unceded traditional territories of the people of the Treaty 7 region in Southern Alberta, which includes the Blackfoot Confederacy (including the Siksika, Piikuni, Kainai First Nations), the Tsuut’ina, and the Stoney Nakoda (including the Chiniki, Bearspaw, and Wesley First Nations). The City of Calgary is also home to Metis Nation of Alberta, Region III.

## Author Contributions

L.T.S, C.B. performed electrophysiological experiments, L.T.S., L.N, A.K.J., T.K., M. E., T. T., performed behavioural experiments, L.T.S., M.Q. performed immunohistochemistry experiments, L.T.S, L.N, C.B., M.A., S.L.B. analyzed the data, S.B.F. provide med associates code, L.T.S., L.N., S.L.B wrote the manuscript, all authors edited.

## Methods

### Animals

All protocols were in accordance with the ethical guidelines established by the Canadian Council for Animal Care and were approved by the University of Calgary Animal Care Committee. Adult (Postnatal day P60) male C57BL6 mice were obtained from Charles Rivers Laboratories or from the Clara Christie Centre for Mouse Genomics (University of Calgary). To target inhibitory neurons, adult P60 male Vgat-ires-cre mice (VGAT^cre^; aka Slc32a1tm2(cre)Lowl/J) obtained from Jackson Laboratory (strain: 016962) and bred in the Clara Christie Centre for Mouse Genomics. Animals were group housed (3-5 animals per cage) for the duration of the dietary manipulations. Mice were singly caged for operant training, as light food restriction was required.

### Diets

The control and high-fat diet were obtained from Research diets (New Brunswick, NJ). The control diet (D12450J) was composed of 20% protein, 70% carbohydrate and 10% fat from calories (3.82kcal/g). The high-fat diet (D12492) was composed of 20% protein, 20% carbohydrate, and 60% fat from calories (5.21 kcal/g). Both diets were matched for vitamins, minerals, and sucrose. Animals were maintained on these diets for a minimum of 12 weeks starting in adulthood (postnatal day 60-90) before training and testing (Extended Data Figure 1a).

### Glucose tolerance test

Lean and obese mice either given a high fat or low-fat diet ad libitum or food restricted and were fasted overnight. A baseline blood sample was collected from the tail vein (a small cut 1-2mm from the end of the tail, time 0 in timeline in Extended Data Figure 1b). Animals were then administered a 20% glucose solution (20% D-glucose in 0.09% saline, 2g of glucose per kilogram of body weight) and blood was collected at 11 different time points (15, 30, 45, 60, 75, 90, 105, 120, 150, 180, 210 minutes post-injection). Blood glucose levels were measured with an Accu-Chek Aviva blood glucose meter.

### Devaluation by satiety

Both lean and obese mice were mildly food restricted and maintained at 85% of their original weight throughout training and testing. Instrumental responding for a 30% sucrose solution was performed in sound- and light-attenuated Med-associate chambers (St-Albans, Vermont) equipped with a retractable active lever. A cue light was positioned above the lever and was illuminated when the lever was active. Chambers were illuminated with a house light, which signalled the beginning of the session. With the appropriate number of lever presses, a 30% liquid sucrose (dissolved in dH_2_O) was delivered into the cup via a syringe connected to a pump. All training consisted of 1h sessions. To train animals to lever press, we shaped behaviour by baiting the lever with sucrose during a fixed ratio (FR) 1 schedule of reinforcement.

To escalate responding, we switched to a random ratio (RR) 5, 10, and 20 schedules of reinforcement. During RR, rewards are delivered, on average, every 5, 10, or 20 lever presses, but not every 5, 10, or 20 times. Devaluation by satiety occurred over two testing days and the “valued” and “devalued” conditions were counterbalanced for every diet and manipulation. Water was removed prior to behavioural testing (zeitgeber time (ZT) 9). At ZT 21, a bottle of either H_2_0 (“valued”) or 30% sucrose (in dH_2_0, “devalued”) was introduced into the cage and the animals were allowed to drink freely for 3 hours. In one cohort, access to either water or sucrose occurred within the operant chambers, behavioural testing occurred at ZT0. The test period consisted of a 10 min unreinforced (no sucrose reward delivery) testing period. One cohort of lean and obese animals had a 10 min reinforced session (with sucrose delivery). In one cohort, mice were given either 25% sucrose and 5% cherry Kool-Aid (“valued” and matched for calories) or 30% sucrose (in dH20, “devalued”). Mice were allowed to drink freely for 1 hour. After test mice were given a 1 -hour choice test where both sucrose and cherry bottles were available in their home cages. A minimum of 5 lever presses on the valued and devalued days (total lever presses) was used as criteria to be included in the study. Further, if there were 0 lever presses in the valued condition, mice were excluded due to a lack of engagement in the task. In the satiety-induced devaluation data, 3 high fat and 1 low fat mice were excluded. In the restoration of sensitivity to devaluation by NNC711 experiment, 4 lean mice and 3 obese mice were excluded. Furthermore, there were 4 missed cannula placements, and these mice were excluded from the data.

### Progressive ratio

A subset of animals used in the satiety devaluation procedure was re-trained to lever press on a RR10 schedule. Some animals were trained to lever press on an FR10 schedule. The performance on the progressive ratio schedule was identical and therefore animals with different training (RR10 vs. FR10) were grouped together. They were then tested on a progressive ratio (PR) schedule. Under this schedule, the number of responses required to obtain each successive sucrose reward was determined by the following equation: Response ratio = [5e^(injection number × 0.2)^]-5, to produce the following sequence of required lever presses: 1, 2, 4, 6, 9, 12, 15, 25, 32, 40, 50, 62, etc. The daily PR sessions were terminated when 1 hour elapsed without sucrose deliver and, as a result, generally lasted 3 hours or less. The maximal number of presses emitted to attain the final ratio was defined as the breakpoint^49^.

### Contingency degradation

A subset of animals used in the devaluation procedure were re-trained to undergo contingency degradation. In this experiment, animals were maintained on a RR20 schedule of reinforcement, as above. During testing, the reinforcement contingency was altered such that sucrose delivery was explicitly not contingent on lever presses. Instead, the sucrose was set to be delivered at a 20 second variable interval schedule. A house light and a cue light were illuminated above the lever for the entirety of the session to signal session duration. Testing occurred over 2 consecutive days (1h sessions; 1 RR20 day (non-degraded (ND) days) + 1 contingency degradation day (CD)). The number of lever presses and the number of rewards earned were recorded.

### Devaluation by conditioned taste avoidance

To test for conditioned taste avoidance (CTA), lean adult C57BL/6 mice underwent a 6-day taste avoidance-conditioning paradigm. Three days prior to conditioning, mice were briefly exposed to grape and orange Kool-Aid^TM^ flavoured gelatine (Knox Gelatine) in their home cages to reduce food neophobia. To minimize stress, animals were brought to the testing room and remained in the room for 1 h before conditioning. During conditioning, each animal consumed a flavour of gelatine that was either paired with sickness inducing LiCl (40mL/kg of 0.24M LiCl, i.p.) or vehicle (40mL/kg 0.9% saline, i.p) over 3 conditioning days.

Each animal received the paired injection with one flavour and the unpaired injection with different flavour every second day for a total of 6 consecutive days of injections. Flavours paired with LiCI were counterbalanced. The two different gelatine exposures were administered in distinct environmental contexts. One context consisted of a smooth cage bottom, a paper house, with gelatine in a square plastic weigh boat. The second context had white paper towel on the cage bottom and gelatine was delivered in a plastic circle weigh boat. After 1h access to the flavoured gelatine, the remaining food was removed and weighed, and mice were immediately injected with either VEH or LiCl. Mice were then placed back into their conditioning cage for 1h before being moved to their home cage. Test days were counterbalanced and consisted of exposure to either orange of grape flavour on day 1 and orange and grape flavour on day 2. In the diet induced obesity experiments, after CTA, mice were randomized to either a high-fat or a low-fat diet for 12 weeks. Mice were then re-exposed to grape or orange gelatine for 1 h on separate days and gelatine consumption was measured. The “valued” state was exposure to the unpaired gelatine flavour, whereas the “devalued” state was exposure to the LiCl-paired gelatine flavour.

### Chemoqenetic, optogenetic and pharmacological targeting of inhibitory neurons

To target inhibitory neurons of the lOFC, we used VGAT^cre^ mice. All surgical procedures occurred under isoflurane anaesthesia. Animals were placed in a stereotaxic instrument (David Kopf Instruments) and administered ketoprofen (10mg/kg, subcutaneous) for pain management. A small incision on the skull was made and bregma was identified. All measurements were made relative to bregma. Virus injections were made using a Drummond Nanoinjector and precise delivery of the virus was performed via glass pipettes. To inhibit the GABAergic neurons of the lOFC, we transduced either a viral vector containing the inhibitory G-protein coupled Designer Receptor Activated Exclusively by a Designer Drug (DREADD) (pAAV8-hsyn-DIO-hM4D(Gi)-mCherry, Addgene) or the control virus (pAAV8-hSyn-DIO-mCherry, Addgene) to the GABAergic neurons of the lOFC (coordinates: anterior posterior (AP):2.6mm, medial lateral (ML): ±1.3mm, dorsal ventral (DV): −1.445 (100nl) and −1.945 (100nl)). We waited 3-5 weeks for transgene expression. For the selective satiety experiments, animals were trained to lever press for 30% sucrose as described above. To activate the DREADD receptor, animals received an IP injection of clozapine N-oxide (CNO; Abcam, dissolved in 0 9% saline, 2mg/kg body weight) 1h prior to behavioural testing at ZT 23. For the CTA experiment, animals were trained as above and received an IP injection of CNO (2 mg/kg) 30 minutes prior to behavioural testing (valued and devalued condition).

For optogenetic activation of GABAergic neurons, we transduced a viral vector containing Channel Rhodopsin 2 (ChR2, AAV2-EF1a-DIO-hChR2-EYFP, Canadian Neurophotonics platform) or the control virus (AAV2-EF1a-DIO-EYFP, Canadian Neurophotonics platform) to the GABAergic neurons of the lOFC (coordinates: AP: 2.6mm, ML: ±1.3mm, DV: −1.445 (55.2nl) and −1.945 (55.2nl)). The location of virus expression was performed posthoc. VGAT-Ires-Cre mice were placed on diets for at least 12 weeks prior to viral transduction. We waited a minimum of 3 weeks for transgene expression and mice continued to train during this time. Two-ferrule flat-tip cannulas (fibre length: 2.5mm, Pitch length: 2.6mm, Doric Lenses) were implanted in the lOFC to allow for delivery of light to the lOFC and activation of ChR2 (coordinates: AP: 2.6mm, ML: ±1.3mm, DV: −1.445). All training and testing occurred as previously described (selective satiety). Optical fibres inserted into the cannula were connected to lasers (473 nm and 589 nm from Laser Glow Technologies) via patch cords. Prior to testing for selective satiety, all mice were habituated (minimum of 3 days) to the insertion of the optical fibre and connection to patch cords prior to performance on RR20. During testing, patch cords were connected to either the 473 nm or 589 nm lasers in the home cage and light was delivered (5Hz for one second, every 10 s for 5 min). Mice were then immediately placed in the operant chambers for testing.

For intracranial cannulations, mice were anesthetized with isoflurane, injected with subcutaneous ketoprofen (10mg/kg) and saline (0.5-1 mL, 0.9%), and secured in the stereotaxic frame. Bilateral 23-gauge cannula (Plastics one) were implanted into the lOFC and secured with vetbond and dental acrylic. Stereotaxic coordinates for cannulae were (in mm) AP: +2.6 (from Bregma), ML: ±1.3, and DV: −1.44 (from dura). NNC-711 (10 μg, 0.2 μl over 3 min) was microinjected into the lOFC 30 min prior to the test during the last 30 min of the sucrose/water pre-feed.

### Electrophysiology

To examine whether diet-induced obesity alters the activity or inhibitory synaptic transmission of lOFC pyramidal neurons, we performed electrophysiology recordings in slice preparations from lean and obese mice. Animals were anesthetised with isoflurane and transcardially perfused with an ice-cold sucrose solution containing (in mM) 50 sucrose, 26.2 NaHCO3, 1.25 glucose, 4.9 MgCl2, 3 kynurenic acid, 0.1 CaCl2 and 1.32 ascorbic acid in bicarbonate-buffered artificial cerebrospinal fluid solution (aCSF in mM: 126 NaCl, 1.6 KCl, 1.1 NaH2PO4, 1.4 MgCl2, 2.4 CaCl2, 26 NaHCO3, 11 glucose). Mice were quickly decapitated, brains were extracted and 250 μm coronal sections containing the lOFC were prepared using a vibratome (Leica, Nussloch, Germany) in the same sucrose solution and then incubated in a holding chamber containing heated (32°C) aCSF for a minimum of 45 minutes. Sections were then transferred to the recording chamber and super fused with aCSF maintained at 32°C. lOFC cells were visualized on an upright microscope using “Dodt-type” gradient contrast infrared optics (Dodt et al., 2002) and whole-cell recordings were made using a MultiClamp 700B amplifier (Axon Instruments, Union City, CA) and collected with pClamp10. Pyramidal neurons in layer II/III were identified by morphological characteristics and were recorded approximately 100-300 μm above the inflection point of the rhinal sulcus.

For current clamp experiments (cellular activity), recording electrodes (3-5 MΩ) were filled with (in mM) 130 potassium-D-gluconate, 10 KCl, 10 HEPES, 0.5 EGTA, 10 sodium creatine phosphate, 4 Mg-ATP and 0.3 Na2GTP. After breaking into the cell, membrane resistance was recorded in voltage clamp. Cells were then switched into current clamp mode and the membrane potential for each neuron was set to −70 mV by current injection via the patch amplifier. A current step protocol consisting of 21 steps (0-500pA, 25pA increments, 400ms in duration, 3 seconds apart) was applied and the number of action potentials at each step was recorded. For some experiments, the current step protocol was recorded 10 min after application of picrotoxin and compared to the baseline recording. For optogenetic experiments, the current step protocol was performed prior to and following optogenetic activation (5Hz x 5 pulses for 40 seconds of Light Emitting Diode (LED)) through the 40x objective). For the NNC-711 experiment, baseline evoked firing was examined using a current step protocol. Bath application of either NNC-711 in aCSF (Tocris, 10 μM) or aCSF occurred for 10 min and the current step protocol was applied a second time.

For voltage clamp experiments (miniature inhibitory postsynaptic currents (mIPSCs), evoked inhibitory post synaptic currents (eIPSCs), tonic GABA currents, and paired pulse ratio (PPR)), recording electrodes were filled with a cesium chloride solution (CsCl) internal solution consisting of (in mM) 140 CsCl, 10 HEPES, 0.2 EGTA, 1 MgCl2, 2 MgATP, 0.3 NaGTP, 5 QX-314-Cl. DNQX (10 μM), strychnine (1 μM), DPCPX (1 μM), CGP-35348 (1 μM) and tetrodotoxin (TTX; 500 nM) were added to the aCSF to isolate GABA_A_ mIPSCs recorded at −70 mV. Evoked inhibitory post synaptic currents (eIPSCs), and paired pulse ratio (PPR) were done in the presence of DNQX (10 μM), strychnine (1 μM), DPCPX (1 μM), CGP-35348 (1 μM) and APV (10μM). IPSCs were filtered at 2 kHz, digitized at 10 kHz and collected on-line using pCLAMP 10 software. For tonic GABA experiments gabazine (1 00μM) was added to aCSF bath which included DNQX (10 μM), strychnine (1 μM), DPCPX (1 μM), CGP-35348 (1 μM) and APV (10 μM)) and change in holding potential and RMS noise was obtained by a Gaussian fit to an all points histogram over a 5 second interval^50^. In some experiments NNC-711 (10 μM) was washed onto the slice. The resting membrane potential as defined as the potential generated across the cell membrane by the difference in charge from internal and external solutions was calculated 2 minutes after achieving whole cell configuration by the amplifier. The junction potential of +4 mV for CsCl internal or +16.2 mV for KGluconate internal solution was not corrected. Series resistance (6-20 MΩ) and input resistance were monitored on-line with a 5-mV depolarizing step (50 ms). Recordings exhibiting a >20% change in series resistance were discarded. GABA_A_ mIPSCs were selected for amplitude (>12 pA), rise time (<4 ms), and decay time (<10 ms). Analysis of miniature events was performed blind to the experimental condition.

### Immunohistochemistry

Following behavioural experiments, VGAT^cre^ mice were deeply anesthetized with isoflurane and transcardially perfused with phosphate buffered saline (PBS) and then with 4% paraformaldehyde (PFA). Brains were dissected and postfixed in 4% PFA at 4°C overnight, then switched to 30% sucrose and coronal frozen sections were cut at 30 μm using a cryostat. Sections were blocked in normal goat serum (optogenetics experiment) or normal donkey serum (DREADD experiment) before incubation in primary antibody for one hour (Optogenetics experiment: Chicken anti GFP, Aves Labs (cat# GFP-1020) 1:1000, DREADD experiment: Rabbit anti RFP, Rockland (cat#600-401-379), 1:2000) and secondary antibody for one hour (Optogenetics experiment: Alexa Fluor 488 goat anti chicken, Invitrogen (cat# A11039), 1:400, DREADD experiment: Alexa Fluor 594 donkey anti rabbit, Invitrogen (cat#A21207), 1:400). Slices were mounted with histology mountain media (Fluoshield^TM^ with DAPI (Sigma). All images were obtained on an Olympus Virtual Slide Microscopy VS120-L100-W with a 10x objective (Olympus Canada Inc. Ontario, Canada).

### Data analysis and Statistics

Data were analyzed using MS excel. Except for mIPSC analysis which was performed blind to the experimenter, all other data collection and analysis were not performed blind to the conditions of the experiment as it was easy to detect obese vs lean mice. All statistical analyses were performed in GraphPad Prism 9.4.1 (GraphPad, US). When comparing two groups and a within animal design a two-way repeated measures ANOVA was used followed by Sidak’s multiple comparisons post hoc. When comparing two groups with different mice across variables a two-way ANOVA or two-way ANOVA mixed effects model was used depending on if the number of values were matched followed by a Sidak’s multiple comparison post hoc test. A three-way ANOVA or a three-way ANOVA mixed effect model (depending on if values were matched) followed by a Holms’ Sidaks post hoc was used for all data that had three different variables (For example: virus (mCherry vs hM4D(Gi) x drug (vehicle vs CNO) x value (valued vs devalued)). For the training data a simple linear regression test was used to compare the slopes of learning along with a twoway RM ANOVA followed by a Dunnett’s post hoc test was used to compare the RR5 days to RR10 and RR20. Data from electrophysiology frequency-current plots were analyzed using a three-way RM-ANOVA. To determine the mean excitability slope, R^2^ value of individual cells determined by linear regression whereby x = current step and y= number of action potentials. We used paired or unpaired two-sided t-tests for 2 group comparisons. All significance was set at P<0.05. A Shapiro-Wilk test was used to assess for normal distribution of data. All data are expressed as mean ± standard error of the mean (SEM). Individual responses are plotted over averaged responses. Experimental designs and samples sizes aimed at minimizing usage and distress of animals and were sufficient for detecting robust effect sizes. No statistical methods were used to pre-determine sample sizes but our sample sizes are similar to those reported in previous publications^11,21^. For detailed statistics information, see supplementary table 1.

### Data Availability

https://doi.org/10.6084/m9.figshare.21330999

