## Extended data for "Disinhibition of the orbitofrontal cortex biases decision making in obesity"

**Supplemental information**

#### Extended Data Figure legends

**Extended data Figure 1: Body weights and glucose tolerance in lean and obese food restricted or ad libitum fed mice.**

a) Mice fed a low-fat diet on ad libitum (lean ad libitum n=8), or food restricted (lean food restricted n=9) feeding schedules weighed less than mice fed a high fat diet on ad libitum (obese ad libitum n=9) and food restricted (obese food restricted n=8). Two-way RM ANOVA: Feeding schedule effect: F (1, 29) = 1.682, p=0.2048, Diet effect: F (1, 29) = 92.08, P<0.0001****, Diet and feeding schedule interaction: F (1, 29) = 2.771, p=0.1068. We had an a priori hypothesis that mice fed a high fat diet would gain more body weight. Sidak's multiple comparisons comparing food restricted in lean and obese mice P <0.0001**** and comparing lean and obese ad libitum p <0.0001****. Data are presented as mean values +/- SEM.

b) Mice fed a high fat diet (obese ad libitum n=8 and obese food restricted n=8) had reduced insulin sensitivity than mice fed a low-fat diet (lean ad libitum n=8 and lean food restricted n=9) as indicated by blood glucose (nM/L) concentrations following an intraperitoneal injection of 20% D-glucose solution. Three-way RM ANOVA mixed effects model: time x diet x feeding schedule interaction, F (11, 153) = 1.962, p=0.0359*, diet x feeding schedule interaction: F (1, 153) = 0.4945, p=0.4830. time x feeding schedule interaction: F (11, 153) = 2.935, p=0.4830, time x diet interaction F (11, 153) = 10.72, p<0.0001****, feeding schedule effect, F (1, 15) = 1.910, p=0.1872, diet effect, F (1, 153) = 27.63, p<0.0001****, time effect F (11, 165) = 90.76, p<0.0001****. Data are presented as mean values +/- SEM.

c) Mice fed a high fat diet (obese ad libitum n=8 and obese food restricted n=8) had reduced baseline insulin sensitivity than mice fed a low-fat diet (lean ad libitum n=8 and lean food restricted n=9) indicated at time 0 before 20% glucose injection. Two-way RM ANOVA: Schedule effect: F (1, 29) = 0.1033, p=0.7503, Diet effect F (1, 29) = 10.73, p=0.0027**, Schedule and diet interaction: F (1, 29) = 0.05216, p=0.8210. Data are presented as mean values +/- SEM.

d) Mice fed a high fat diet (obese ad libitum n=8 and obese food restricted n=8) have reduced insulin sensitivity than mice fed a low-fat diet (lean ad libitum n=8 and lean food restricted n=9) indicated by area under the curve. Two-way RM ANOVA: Schedule effect: F (1, 29) = 2.152, p=0.1531, Diet effect: F (1, 29) = 27.62, p=<0.0001****, Schedule and diet interaction: F (1, 29) = 0.5155, p=0.4785. We had an a priori hypothesis that mice fed a high fat diet would have prolonged blood glucose levels. Sidak's multiple comparison test comparing lean and obese mice with food restricted p= 0.0058** and ad libitum p= 0.0005*** feeding schedules. Data are presented as mean values +/- SEM.

**Extended data Figure 2: Satiety-induced devaluation and context generalization.**

a) Lean (n=60) and obese (n=60) mice were trained to lever press (RR5, RR10, and RR20 training) for sucrose prior to behavioural testing on the devaluation task (included cohorts from selective satiety, context generalization and flavour preconditioning experiments). Two-way RM ANOVA: diet effect F (1, 118) = 17.04, P<0.0001****, Schedule of reinforcement effect (RR5 vs. RR10 vs. RR20): F (1.310, 154.6) = 18.26, P<0.0001****, Diet x schedule of reinforcement interaction: F (2, 236) = 7.112, p=0.0010. A Dunnett's multiple comparison test compared RR10 and RR20 schedule of reinforcement to RR5 was performed to see if mice were escalating their lever presses from RR5. Lean RR5 vs RR10 P<0.0001****, Lean RR5 vs RR20 p= 0.0006***, Obese RR5 vs RR10 p= 0.0073**, Obese RR5 vs RR20 p=0.0046**. There is no significant difference in slopes. Simple linear regression: F = 3.678. DFn = 1, DFd = 356, p=0.0559, but there was a significant difference in the slope intercepts F = 40.26. DFn = 1, DFd = 357, P<0.0001****.

b) Satiety-induced devaluation procedure.

c) In the devaluation by selective satiety task, mice with long term exposure to a 60% high-fat (obese n=31) diet gained more weight than mice with long term exposure to a 10% low-fat (lean n=35) diet. Unpaired T-test: t (64) =10.88, p<0.0001****. Data are presented as mean values +/- SEM.

d) There was no difference in the consumption of sucrose (ml) of lean (n=35) and obese (n=31) during the 3-hour pre-feeding period prior to devaluation (devalued condition). Unpaired t-test: $t_{(64)}$ =0.6408, p=0.5239. Data are presented as mean values +/- SEM.

e) Lean (n=35) mice devalued the outcome as indicated by decreased lever presses in the devalued compared to the valued condition. Obese mice (n=31) were not sensitive to devaluation as indicated by comparable lever presses in the valued and devalued conditions. RM two-way ANOVA: devaluation effect, F (1,64) =7.8, p=0.0067**, diet group x devaluation interaction, F (1,64) =5.8, p=0.0187*, Sidak's multiple comparisons test, lean: p=0.0006***, obese: p=0.9561. Data are presented as mean values +/- SEM.

f) Obesity decreased the revaluation index ((lever presses valued – lever presses devalued)/ (lever presses valued + lever presses devalued)). Unpaired t-test: lean (n=35), obese (n=31), t (64) =4.109, p=0.0001***. Data are presented as mean values +/- SEM.

g) Total number of lever presses (valued + devalued) of obese mice (n=31) compared to lean mice (n=35) during the devaluation task. Unpaired t-test: t (64) =1.886, p=0.0638. Data are presented as mean values +/- SEM.

h) In the context generalization satiety-induced devaluation task, mice with long term exposure to a 60% high-fat (obese n=16) diet gained more weight than mice with long term exposure to a 10% low-fat (lean n=12) diet. Unpaired T-test: t (26) =10.18, p<0.0001****. Data are presented as mean values +/- SEM.

i) There was no difference in the consumption of sucrose (ml) of lean (n=12) and obese (n=16) mice during the prefeed. Unpaired T-test t (26) =0.3258, p= 0.7472. Data are presented as mean values +/- SEM.

j) In the sensory-specific satiety devaluation task, mice with long term exposure to a 60% high-fat (obese n=13) diet gained more weight than mice with long term exposure to a 10% low-fat (lean n=12) diet. Unpaired T-test: t (23) =6.068, p<0.0001****.

**Extended data Figure 3: Sucrose reinforced devaluation and progressive ratio.**

a) Devaluation procedure.

b) When sucrose is delivered during the devaluation test, lean mice (n=8) devalued the sucrose reward (reduced lever presses in the devalued condition), while obese mice (n=13) did not (similar number of lever presses during valued and devalued condition). Two-way RM ANOVA: devaluation effect: F (1, 19) = 5.283, p=0.0331, diet effect: F (1, 19) = 0.9864, p=0.3331, diet x devaluation interaction: F (1, 19) = 2.541, p=0.1274. A priori hypothesis was that sated lean mice would devalue the sucrose reward. Sidak's multiple comparisons test: lean valued vs. devalued p=0.0454*, obese valued vs. devalued p=0.8193. Data are presented as mean values +/- SEM.

c) Revaluation index of lean (n=8) and obese (n=13) mice in the devaluation task. Unpaired t-test: $t_{(19)}$ =2.481, p=0.0226*. Data are presented as mean values +/- SEM.

d) Total lever presses (valued + devalued) were not altered by diet (lean (n = 8) vs. obese (n = 13)). Unpaired t-test $t_{(19)}$ =0.9932, p = 0.3331. Data are presented as mean values +/- SEM.

e) Weights of lean (n=15) and obese (n=11) mice in the progressive ratio test. Unpaired T-test: t (24) =13.25, p<0.0001****. Data are presented as mean values +/- SEM.

f) Number of lever presses of lean (n=15) and obese (n=11) mice during progressive ratio. Unpaired t-test: t (24) =1.521, p=0.1412. Data are presented as mean values +/- SEM.

g) Number of sucrose reinforcers received by lean (n=15) and obese (n=11) mice during progressive ratio. Unpaired t-test: t (24) =1.282, p=0.2122. Data are presented as mean values +/- SEM.

h) Breakpoint obtained by lean (n=15) and obese (n=11) mice during progressive ratio. Unpaired t-test: t (24) =1.234, p=0.2293. Data are presented as mean values +/- SEM.

i) Lean (n=15) but not obese mice (n=16) learn the contingency change when rewards given regardless of lever presses. Two-way ANOVA: Diet effect F (1, 29) = 1.575, p=0.2195, Contingency effect: F (1, 29) = 19.97, p=0.0001 ***, Diet x contingency interaction F (1, 29) = 1.734, p=0.1982. We had an a priori hypothesis that lean mice would devalue and obese mice would not. Sidak's multiple comparison comparing lean non degraded (ND) vs contingency degraded (CD) day p= 0.0007 ***, obese non degraded and contingency changed day p= 0.0612. Data are presented as mean values +/- SEM.

**Extended data Figure 4: Obese mice have continued impairment of devaluation 7d after diet removal.**

a) Weights of mice before and after 7 days removal of diet following exposure to either the low-fat (n=9) or high-fat diet (n=8). RM Two-way ANOVA: Diet effect F (1, 15) = 37.03, p<0.0001****, Time effect F (1, 15) = 13.08, p=0.0025**, Diet x time interaction F (1, 15) = 4.772, p=0.0452. Sidak's multiple comparisons test: lean: pre and post diet removal p<0.0001****, obese: pre and post diet removal. p<0.0001****. Data are presented as mean values +/- SEM.

b) Lean (n=9) and obese (n=8) mice were trained to lever press (RR5, RR10, and RR20 training) for sucrose prior to behavioural testing on the devaluation task. Two-way RM ANOVA: Diet effect F (1, 15) = 16.28, p=0.0011**, Training effect F (2, 30) = 2.892, p=0.0710, Diet x training interaction F (2, 30) = 0.3876, p=0.6820. Simple linear regression: Slopes are equal F (1,47) = 0.5650, p=0.4560 and Y intercepts are different F (1,48) = 26.70, P<0.0001 ****. Data are presented as mean values +/- SEM.

c) There was a difference in the consumption of sucrose (ml) of lean (n=9) and obese (n=8) during the 3-hour pre-feeding period prior to devaluation (devalued condition). Unpaired t-test: $t_{(1f)}$ =2.341, p= 0.0335*. Data are presented as mean values +/- SEM.

d) Lever presses during devaluation of lean (n=9) and obese mice (n=8). RM two-way ANOVA: devaluation effect, F (1, 15) = 4.178, p=0.0589, diet effect: F (1, 15) = 5.038, p=0.0403 *, diet group x devaluation interaction, F (1, 15) = 11.97, p=0.0035. Sidaks multiple comparisons lean valued vs devalued p= 0.0023, obese valued vs devalued p=0.5726. Data are presented as mean values +/- SEM.

e) Obesity decreased the revaluation index ((lever presses valued – lever presses devalued)/ (lever presses valued + lever presses devalued)). Unpaired t-test: lean (n=9), obese (n=8), $t_{(15)}$ =2.478, p= 0.0256*. Data are presented as mean values +/- SEM.

f) Total number of lever presses (valued + devalued) of obese mice (n=8) compared to lean mice (n=9) during the devaluation task. Unpaired t-test: $t_{(15)}$ =2.245, p= 0.0403*. Data are presented as mean values +/- SEM.

**Extended data Figure 5: Pyramidal neuron excitability in lean and obese mice on ad libitum or food restricted feeding schedules.**

a) Representative recordings of action potentials observed at 100pA, 300pA and 500pA current steps from lOFC pyramidal neurons of lean and obese mice on ad libitum or food restricted feeding schedules.

b) Obese mice fed high fat diet on ad libitum (n=3) and food restricted (n=3) weight more than lean mice on ad libitum (n=3) and food restricted (n=3) feeding schedule. RM Two-way ANOVA: Diet effect F (1, 8) = 34.16, p=0.0004***, food schedule effect F (1, 8) = 12.93, p=0.0070**, Diet x food schedule interaction F (1, 8) = 0.2985, p=0.5997. We had an a priori that obese mice would weigh more than lean mice regardless of food schedule. Sidak's multiple comparisons test: ad libitum lean vs obese p= 0.0039**, food restricted lean vs obese 0.0113*. Data are presented as mean values +/- SEM.

c) Diet-induced obesity increased the excitability of lOFC pyramidal neurons as indicated by frequency-current (F-I) plots of number of action potentials at current injections from 0pA to 500pA of lean food restricted (n=14 cells/3 mice), lean ad libitum (12 cells/3 mice) and obese food restricted (n=14 cells/3 mice) and obese ad libitum (n=12 cells/3 mice). RM 3-way ANOVA: current step effect F (20, 520) = 509.4, p <0.0001****, diet effect: F (1, 436) = 12.28, p=0.0005***, food schedule effect: F (1, 26) = 4.450, p= 0.0447*, current x diet interaction: F (20, 436) = 13.45, p <0.0001****, current x food schedule interaction: F (20, 436) = 1.792, p= 0.0193*, diet x food schedule interaction: F (1, 436) = 0.1139, p= 0.7359, current x diet x food schedule interaction F (20, 436) = 0.9759, p= 0.4906. Data are presented as mean values +/- SEM.

d) Diet induced obesity (obese food restricted n=14 cells/3mice, obese ad libitum n=12 cells/3mice) increases the firing slope compared to lean mice (lean food restricted n=14 cells/3mice, lean ad libitum n=12 cells/3 mice) in both food restricted and ad libitum feeding schedules. Mean excitability slope (slope of linear regression from individual cells, x=current step y=number of action potentials) of lOFC pyramidal neurons. Two-way RM ANOVA: Diet effect: F (1, 48) = 20.85, P<0.0001, food schedule effect: F (1, 48) = 0.9912, p=0.3244, diet x food schedule interaction: F (1, 48) = 1.106, p=0.2983. Based on previous excitability data we had an a priori hypothesis that obese mice would have hyperexcitable pyramidal neurons. Sidaks multiple comparisons ad libitum lean vs obese p= 0.0007***, food restricted lean vs obese p=0.0254*. Data are presented as mean values +/- SEM.

e) The resting membrane potential in lean (lean food restricted n=14 cells/3mice, lean ad libitum n=12 cells/3 mice) and obese mice (obese food restricted n=14 cells/3 mice, obese ad libitum n=12 cells/3 mice) was not altered based on food schedule. Two-way RM ANOVA: Diet effect: F (1, 48) = 6.144, p=0.0168*, Feeding schedule effect: F (1, 48) = 0.04941, p=0.8250, Diet x feeding schedule interaction: F (1, 48) = 0.2927, p=0.5910. Data are presented as mean values +/- SEM.

**Extended data Figure 6: Action potential characteristics of lOFC neurons from lean and obese mice.**

a) Capacitance is similar in no PTX lean (n=10 cells/3 mice), lean PTX (n=11 cells/3 mice) no PTX obese (n=8 cells/3 mice) and obese PTX (n=9 cells/3 mice) mice. Two-way ANOVA: Diet effect: F (1, 34) = 0.3277, p=0.5708, Picrotoxin effect: F (1, 34) = 3.141, p=0.0853, Diet x picrotoxin interaction: F (1, 34) = 4.573, p=0.0397*. Data are presented as mean values +/- SEM.

b) Resting membrane is similar in no PTX lean (n=10 cells/3 mice), lean PTX (n=11 cells/3 mice) no PTX obese (n=8cells/3mice) and obese PTX (n=9cells/5mice) mice. Two-way ANOVA: Diet effect F (1, 34) = 0.6887, p=0.4124, Picrotoxin effect: F (1, 34) = 0.007646, p=0.9308, Diet x picrotoxin interaction F (1, 34) = 0.7791, p=0.3836. Data are presented as mean values +/- SEM.

c) Input resistance is increased in obese (n=8 cells/3 mice) compared to lean mice (n = 10 cells/3 mice), but no difference in PTX (lean PTX (n=11 cells/3 mice), obese PTX (n=9 cells/5 mice)). Two-way ANOVA: Diet effect: F (1, 34) = 5.360, p=0.0268*, Picrotoxin effect: F (1, 34) = 2.079, p=0.1585, Diet x picrotoxin interaction: F (1, 34) = 5.340, p=0.0270 *. Sidak's multiple comparisons: No PTX lean vs obese mice, p= 0.0062**, PTX lean vs obese, P>0.9999. Data are presented as mean values +/- SEM.

d) Rheobase is decreased in obese (n =8 cells/3 mice) compared to lean mice (n = 10 cells/3 mice), but no difference in PTX (lean PTX (n=11 cells/3 mice), obese PTX (n=9c ells/5 mice)). Two-way ANOVA: Diet effect F (1, 34) = 13.82, p=0.0007***, Picrotoxin effect: F (1, 34) = 4.043, p=0.0523, Diet x picrotoxin interaction: F (1, 34) = 4.174, p=0.0489*. Sidaks multiple comparisons: No PTX lean vs obese mice p= 0.0007***, PTX lean vs obese p=0.41. Data are presented as mean values +/- SEM.

e) Action potential height is similar in no PTX lean (n=10 cells/3 mice), lean PTX (n=11 cells/3 mice) no PTX obese (n=8 cells/3 mice) and obese PTX (n=9 cells/5 mice) mice. Two-way ANOVA: Diet effect: F (1, 34) = 6.050, p=0.0191*, Picrotoxin effect: F (1, 34) = 2.359, p=0.1338, Diet x picrotoxin interaction: F (1, 34) = 0.6723, p=0.4180. Data are presented as mean values +/- SEM.

f) Action potential width is similar in no PTX lean (n=10 cells/3 mice), lean PTX (n=11 cells/3 mice) no PTX obese (n=8 cells/3 mice) and obese PTX (n=9 cells/5 mice) mice. Two-way ANOVA: Diet effect: F (1, 34) = 0.02203, p=0.8829, Picrotoxin effect: F (1, 34) = 1.201, p=0.2807, Diet x picrotoxin interaction: F (1, 34) = 2.864, p=0.0997. Data are presented as mean values +/- SEM.

g) Threshold to fire is similar in no PTX lean (n=10 cells/3 mice), lean PTX (n=11 cells/3 mice) no PTX obese (n=8 cells/3 mice) and obese PTX (n=9 cells/5 mice) mice. Two-way ANOVA, Diet effect: F (1, 34) = 0.04215, p=0.8386, Picrotoxin effect: F (1, 34) = 1.252, p=0.2711, Diet x picrotoxin interaction: F (1, 34) = 0.2539, p=0.6176. Data are presented as mean values +/- SEM.

h) After hyperpolarization height is similar in no PTX lean (n=10 cells/3 mice), lean PTX (n=11 cells/3 mice) no PTX obese (n=8 cells/3 mice) and obese PTX (n=9 cells/5 mice) mice. Two-way ANOVA: Diet effect: F (1, 34) = 0.5353, p=0.4694, Picrotoxin effect: F (1, 34) = 0.04374, p=0.8356, Diet x picrotoxin interaction: F (1, 34) = 1.468, p=0.2340. Data are presented as mean values +/- SEM.

i) After hyperpolarization width is different in no PTX lean (n=10 cells/3 mice), lean PTX (n=11 cells/3 mice) no PTX obese (n=8 cells/3 mice) and obese PTX (n=9 cells/5 mice) mice. Two-way

ANOVA: Diet effect: F (1, 34) = 7.140, p=0.0115 *, Picrotoxin effect: F (1, 34) = 3.777, p=0.0603, Diet x picrotoxin interaction: F (1, 34) = 5.648, p=0.0232 *, Sidaks multiple comparisons: No PTX lean vs obese mice p= 0.0028**, PTX lean vs obese, p=0.9715. Data are presented as mean values +/- SEM.

**Extended data Figure 7: Inactivation of GABAergic neurons (via hM4d(Gi)) impairs satiety-induced devaluation and conditioned taste aversion.**

a) Body weights of mCherry (n=7) and hM4d(Gi) (n=9) are similar: Unpaired t-test: $t_{(14)}$ =0.1395, p=0.8910. Data are presented as mean values +/- SEM.

b) Lean mCherry (n=7) and hM4d(Gi) (n=9) were trained to lever press on a RR5, RR10 and RR20 schedule. The learning slope is not different between lean and obese mice. Simple linear regression of slopes: F (1,44) = 0.02496, p=0.8752. There is no difference in the intercepts: F (1,45) = 0.01782, p=0.8944. Data are presented as mean values +/- SEM.

c) Lean mCherry (n=7) and hM4d(Gi) (n=9) drink the same amount of sucrose in the pre consumption test. Two-way ANOVA: virus effect: F (1, 14) = 2.549, p=0.1327, Drug effect: F (1, 14) = 3.064, p=0.1019, Drug and diet interaction: F (1, 14) = 0.001013, p=0.9751. Data are presented as mean values +/- SEM.

d) Body weights of mCherry (n=12) and hM4d(Gi) (n=13) are the same: Unpaired t-test: $t_{(23)}$ =0.7069, p=0.4867. Data are presented as mean values +/- SEM.

e) Lean mCherry (n=12) and hM4d(Gi) (n=13) underwent pairing where a novel gelatine flavour was paired with LiCl inducing sickness, or vehicle. 3-way RM ANOVA: Conditioning day effect: F (1.924, 44.25) = 6.360, p=0.0041**, virus effect: F (1, 23) = 2.344, p=0.1394, value effect F (1.000, 23.00) = 0.3603, p=0.5542, day x virus interaction: F (2, 46) = 0.1939, p=0.8244, day x value interaction: F (1.890, 43.46) = 0.4546, p=0.6267, virus x value interaction: F (1, 23) = 0.002321, p=0.9620, day x virus x value F (2, 46) = 1.432, p=0.2493. Data are presented as mean values +/- SEM.

**Extended Figure 8: NNC-711 in the lOFC restores devaluation in obesity.**

a) Weights of lean and obese mice following long-term exposure to either the 10% low fat (lean n=14) or 60% high fat (obese n=16) diet in the devaluation by selective satiety task. Unpaired T-test: $t_{(28)}$ =7.818, p<0.0001****. Data are presented as mean values +/- SEM.

b) Number of lever presses performed on the last two days of RR5, RR10, RR20 training prior to behavioural testing for selective satiety devaluation of lean (n=14) and obese (n=16) mice. Two-way RM ANOVA: Diet effect: F (1, 28) = 24.80, P<0.0001****, training schedule effect: F (1.441, 40.35) = 11.45, p=0.0005***, diet x training schedule interaction: F (2, 56) = 3.103, p=0.0527. We expected that both lean and obese mice would escalate their lever presses. Dunnett's multiple comparison: lean RR5 vs RR10 p= 0.4182, lean RR5 vs RR20 p= 0.0171*, obese RR5 vs RR10 p= 0.0554, obese RR5 vs RR20 p= 0.0102*. Data are presented as mean values +/- SEM.

c) Sucrose presconsumption of lean vehicle mice (n=12), lean NNC mice (n=12), and obese vehicle mice (n=14) and obese NNC mice (n=15). Two-way RM ANOVA mixed effects model: Interaction: F (1, 21) = 0.03347, p= 0.8566, diet effect: F (1, 28) = 0.5022, p=0.4844, Drug effect: F (1, 21) = 0.007126, p=0.9335. Data are presented as mean values +/- SEM.

d) Total distance travelled (m) of lean (n=7) and obese (n=6) mice in a one-hour open field test immediately following lOFC infusions of either vehicle or NNC-711 (NNC-711 10$\mu$M). Two-way

RM ANOVA: Interaction: $F (1, 11) = 0.1175$, $p = 0.7383$, vehicle vs. NNC-711 effect: $F (1, 11) = 0.7986$, $p=0.3906$, diet effect: $F (1, 11) = 16.76$, $p=0.0018$*. Based on previous literature and because there was a diet effect, we expected that obese mice would travel less distance than lean mice. Sidaks multiple comparison test: vehicle lean vs obese $p= 0.0172$*, NNC-711 lean vs obese $p=0.0061$**. Data are presented as mean values +/- SEM.

**Extended Figure 9: Optogenetic activation of lOFC inhibitory neurons restores devaluation in obesity.**

a) Electrophysiology validation of ChR2 expressed in GABAergic neurons of the lOFC. Example traces of current evoked (pA) from a lOFC $GABA^{ChR2}$ neuron to a train of light pulses delivered at 5Hz for 4 ms (5 mW).

b) Electrophysiology validation of ChR2 expressed in GABAergic neurons of the lOFC. Example traces of current evoked (pA) from a lOFC $GABA^{ChR2}$ neuron to a single 500 ms pulse in voltage clamp. This was reliably repeated in several different brain slices from $VGAT^{cre}$ mice.

c) $VGAT^{cre}$ mice exposed to the high-fat diet (n = 7) weighed more than lean mice (n = 8) prior to behavioural testing for devaluation by selective satiety. Unpaired t-test: $t_{(13)} =7.531$, $p<0.0001$****. Data are presented as mean values +/- SEM.

d) Lean (n=8) and obese (n=7) $VGAT^{cre}$ mice were trained to lever press (RR5, RR10, and RR20 training) for sucrose prior to behavioural testing on the devaluation task. Two-way ANOVA mixed effects model: training schedule effect: $F (1.504, 19.55) = 9.122$, $p=0.0031$**, Diet effect: $F (1, 13) = 32.23$, $P<0.0001$****, Schedule of reinforcement and diet interaction: $F (2, 26) = 4.331$, $p=0.0238$ *. Dunnett's multiple comparisons: Lean RR5 vs RR10 $p= 0.9403$, lean RR5 vs RR20 $p= 0.0662$, obese RR5 vs RR20 $p= 0.0109$*, obese RR5 vs RR20 $p= 0.0108$*. Data are presented as mean values +/- SEM.

e) Lean (lean inactive 589 nm n=8, lean active 473 nm n=8) and obese (obese inactive 589nM n=6, obese active 473 nm n=7) mice consumed similar amounts of sucrose prior to the selective satiety procedure. Two-way ANOVA mixed effects model: Diet effect $F (1, 13) = 0.02776$, $p=0.8702$, Stimulation effect $F (1, 11) = 1.468$, $p=0.2510$, diet x stimulation interaction $F (1, 11) = 1.468$, 0.2510. Data are presented as mean values +/- SEM.

f) Total distance travelled (m) of lean (n=6) and obese (n=3) mice in a one-hour open field test immediately following photo-stimulation (589 nm inactive or 473 nm active) of lOFC GABAergic neurons. Two-way RM ANOVA: Interaction: $F (1, 7) = 0.01176$, $p=0.9167$, Diet effect: $F (1, 7) = 3.399$, $p=0.1078$ photostimulation effect: $F (1, 7) = 0.8146$, $p=0.3967$. Data are presented as mean values +/- SEM.

#### Body weights and glucose tolerance

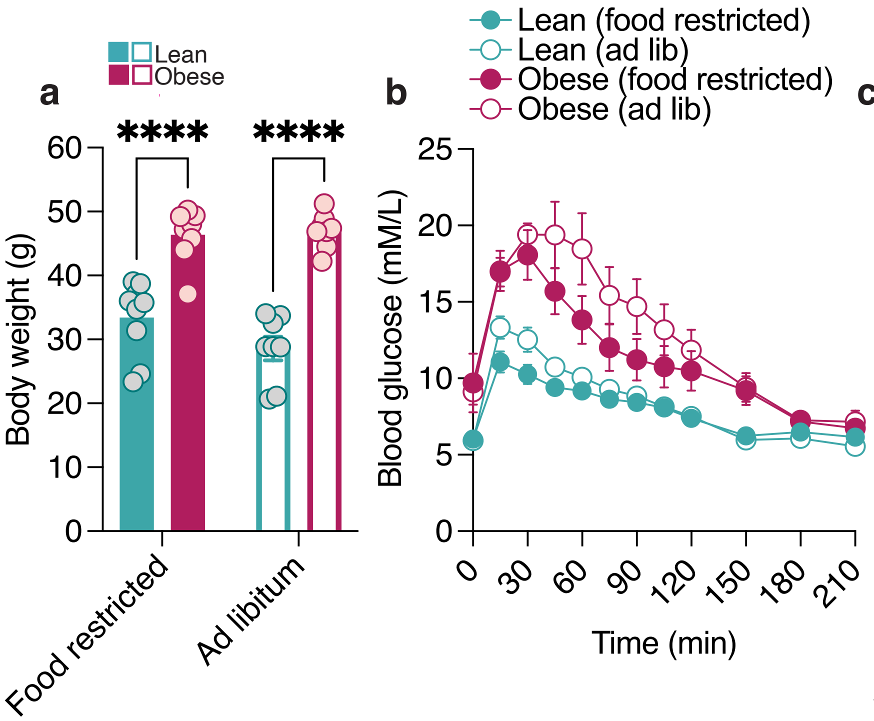

Extended Data Figure 1

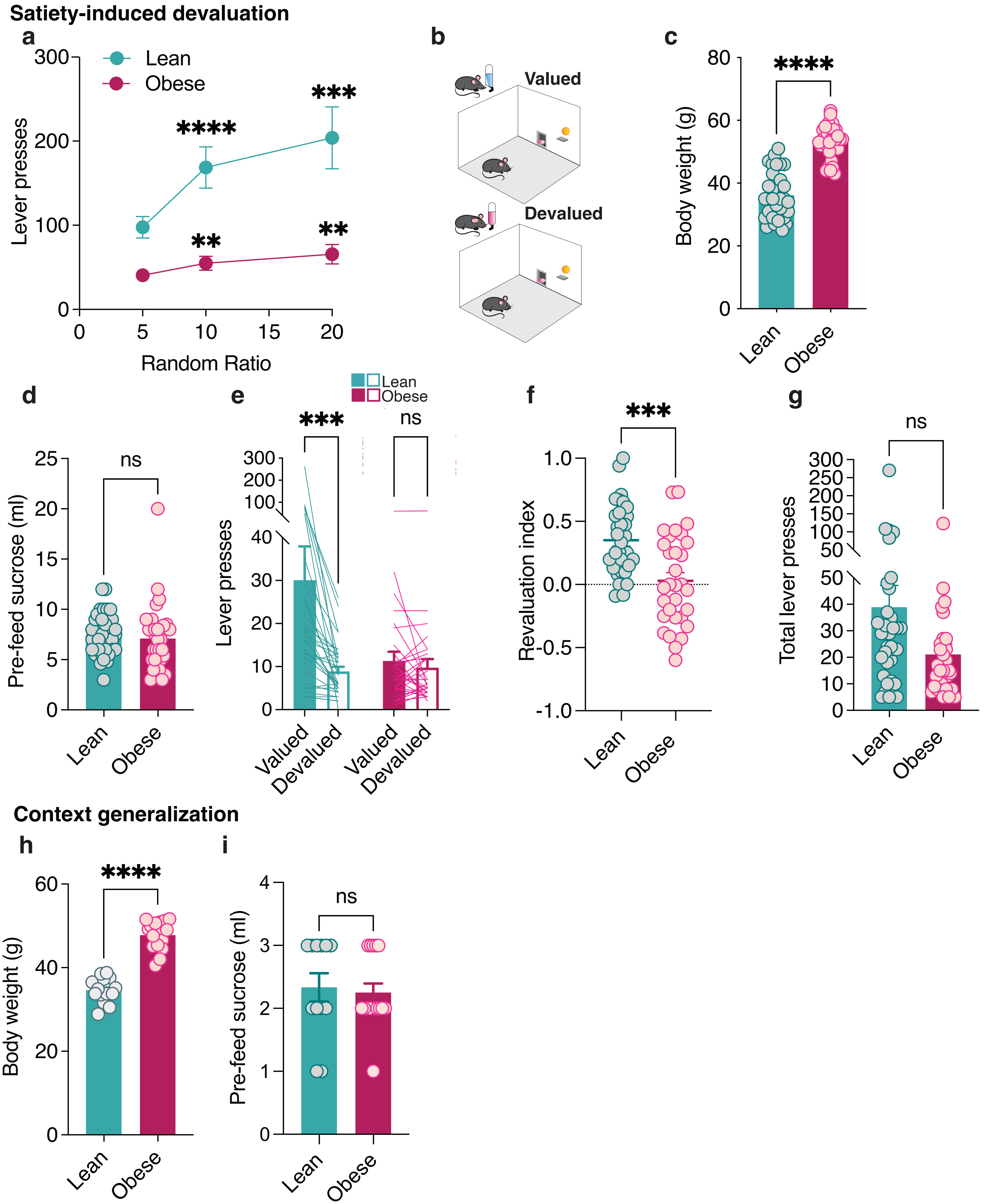

Extended Data Figure 2

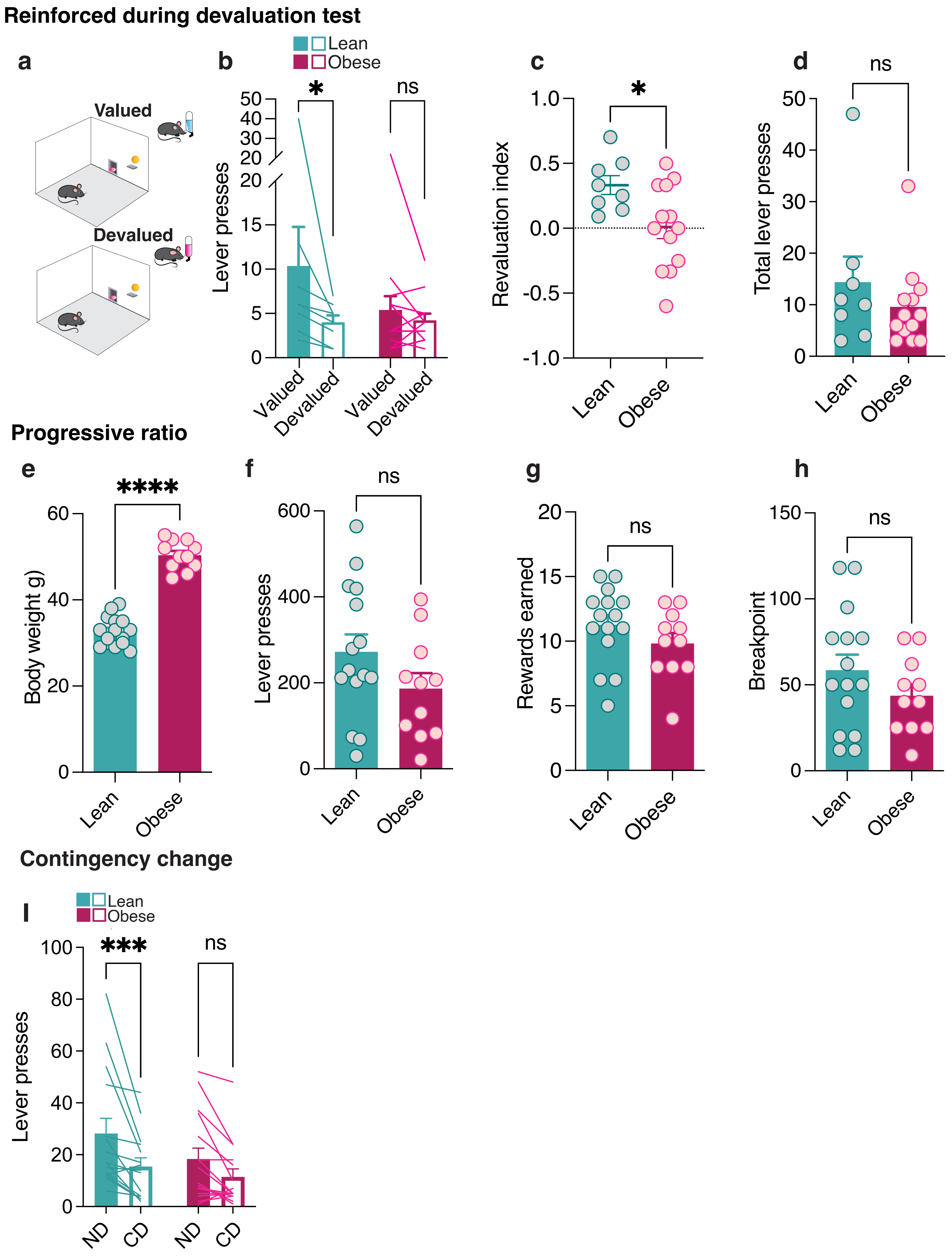

Extended Data Figure 3

**Action potential characteristics**

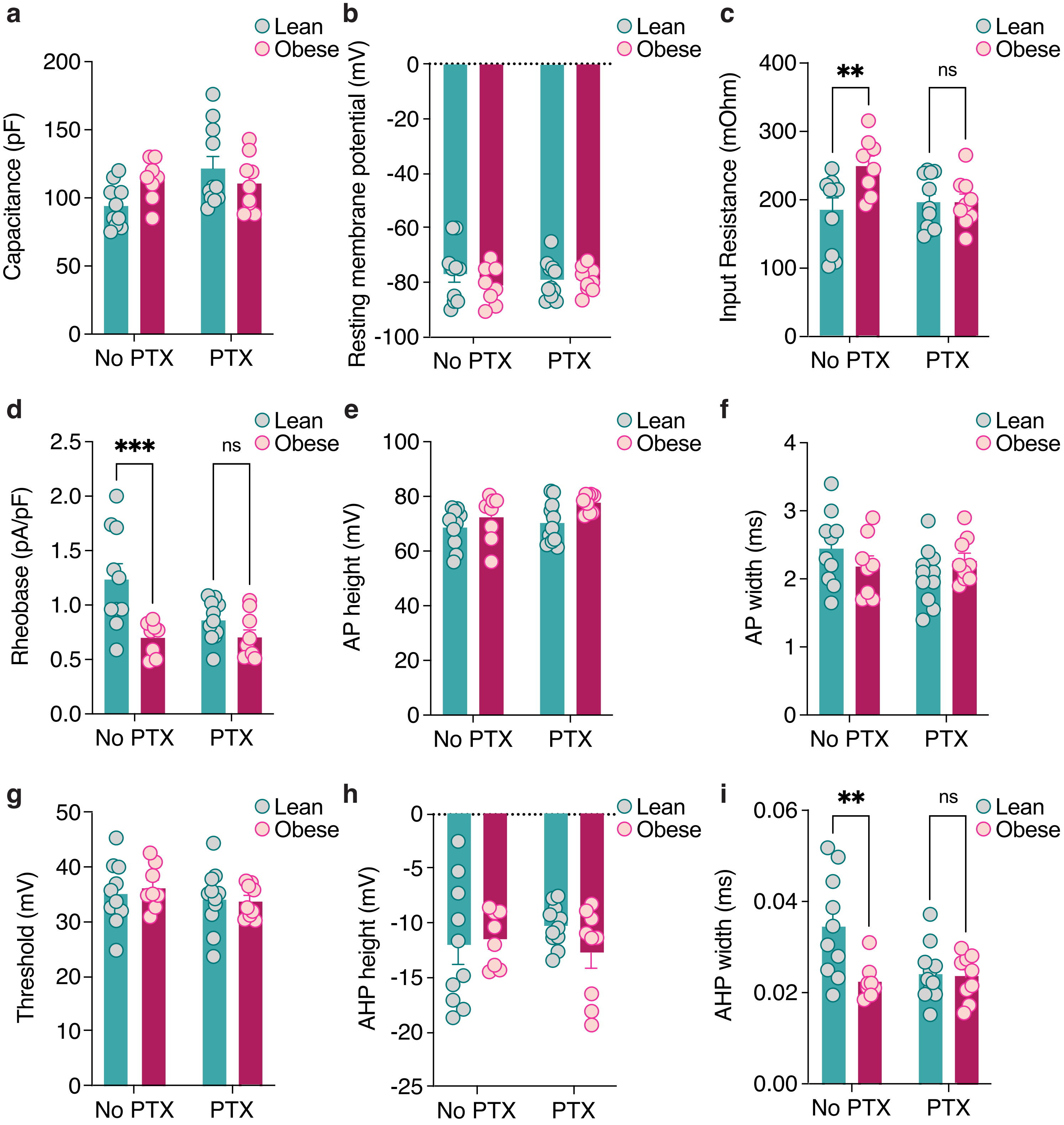

Extended Data Figure 6

### Pyramidal neuron excitability on ad libitum or food restricted feeding schedules

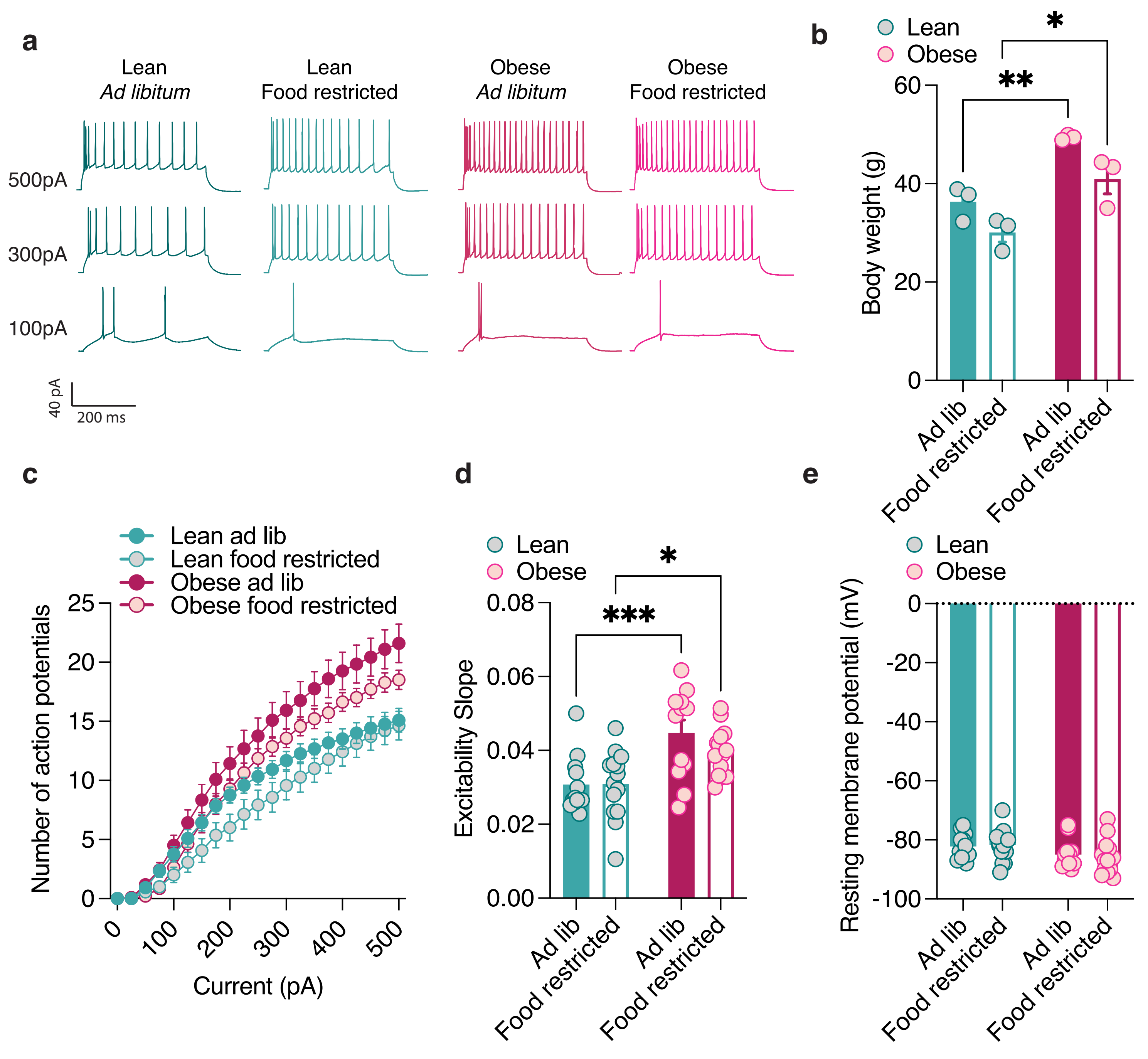

Extended Data Figure 5

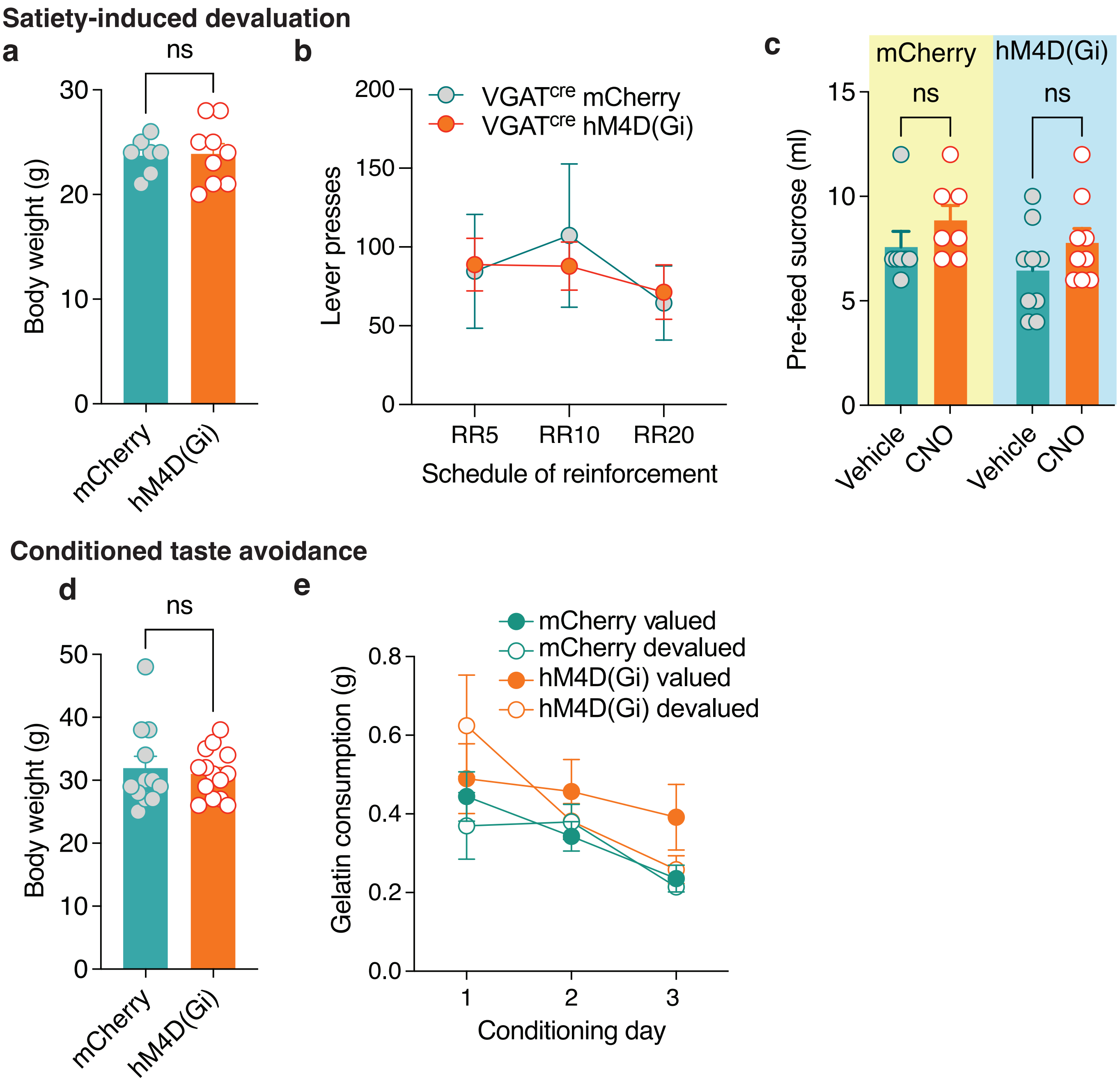

Extended Data Figure 7

7 day HFD removal -  devaluation

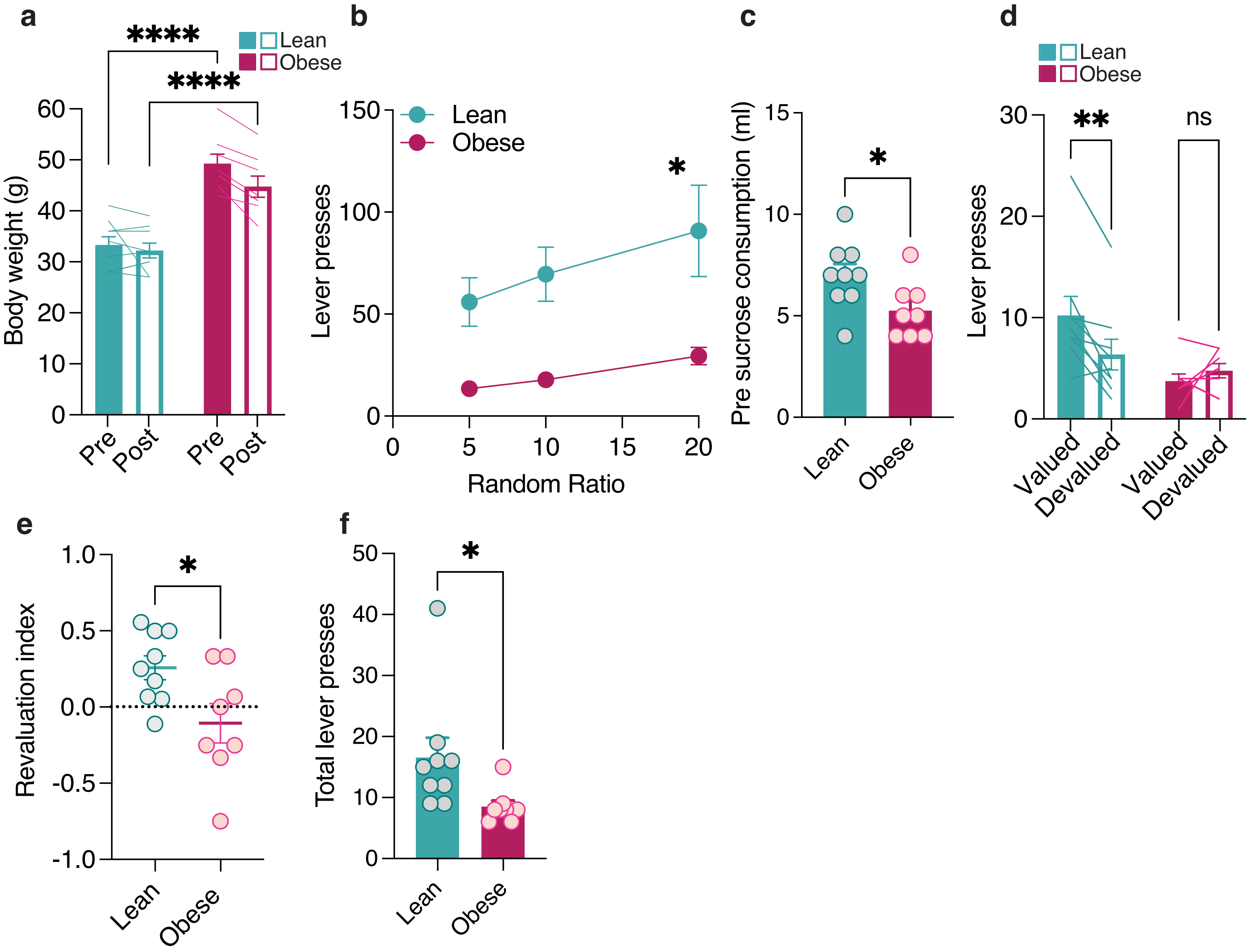

Extended data 4

**lOFC NNC-711: satiety-induced devaluation**

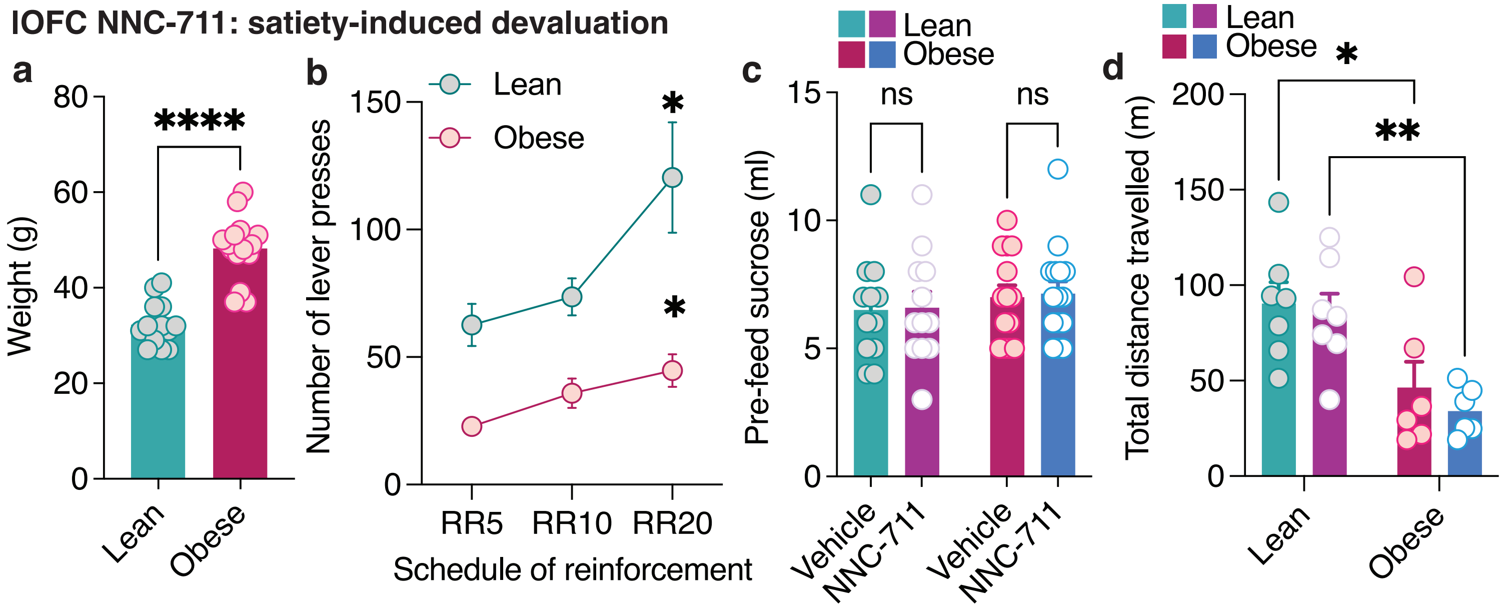

Extended Data Figure 8

**473 nM photostimulation of GABAergic neurons**

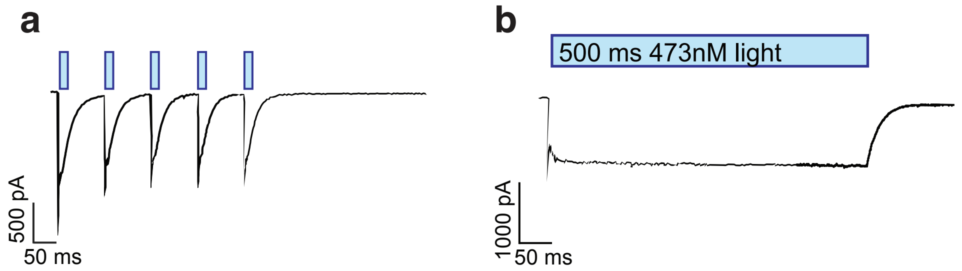

**lOFC optogenetics: satiety-induced devaluation**

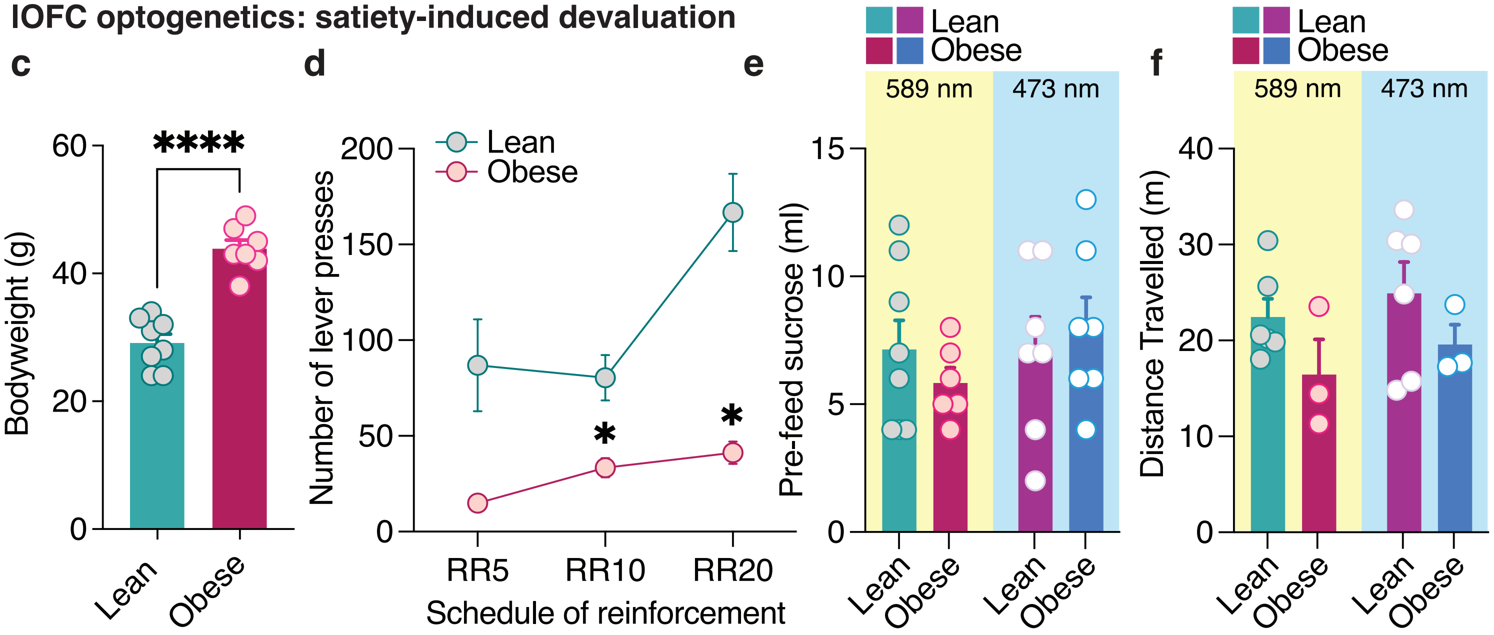

Extended Data Figure 9

Supplemental Table 1. Statistics table.

| V | Test Used | | n | | | Descriptive statistics (Average, Variance) | P - Value | | Degrees of freedom & F/t/z/R | | Multiple comparisons |
|---|---|---|---|---|---|---|---|---|---|---|---|
| Figure # | Test | Section & paragraph # | Exact devaluation | Defined | Section & paragraph # | Reported | Exact devaluation | Section & paragraph # | Value | Section & paragraph # | |
| 1b | two-way RM ANOVA followed by a Sidak's multiple comparisons test | fig legend | lean n=15 mice<br><br>obese n=20 mice | # animals | fig legend | error bars are mean +/- SEM | diet x devaluation interaction p=0.0203<br><br>diet effect p=0.0970<br><br>devaluation effect p=0.0118 | fig legend | diet x devaluation interaction F (1, 33) = 5.949<br><br>diet effect F (1, 33) = 2.919<br><br>devaluation effect F (1, 33) = 7.098 | fig legend | lean p=0.0038<br><br>obese p=0.9816 |
| 1c | Unpaired T-test | fig legend | lean n=15 mice<br><br>obese n=20 mice | # animals | fig legend | error bars are mean +/- SEM | p=0.0262 | fig legend | t=2.327, df=33 | fig legend | |
| 1d | Unpaired T-test | fig legend | lean n=15 mice<br><br>obese n=20 mice | # animals | fig legend | error bars are mean +/- SEM | p=0.0970 | fig legend | t=1.709, df=33 | fig legend | |
| 1f | two-way RM ANOVA followed by a Sidak's multiple comparisons test | fig legend | lean n=12 mice<br><br>obese n=16 mice | # animals | fig legend | error bars are mean +/- SEM | diet x devaluation interaction p=0.0292<br><br>diet effect p=0.1335<br><br>devaluation effect p=0.0497 | fig legend | diet x devaluation interaction F (1, 26) =5.330<br><br>diet effect F (1, 26) =2.399<br><br>devaluation effect F (1, 26) =4.239 | fig legend | lean p=0.0336<br><br>obese p=0.8123 |
| 1g | Unpaired T-test | fig legend | lean n=12 mice<br><br>obese n=16 mice | # animals | fig legend | error bars are mean +/- SEM | p=0.0302 | fig legend | t=2.293, df=26 | fig legend | |
| 1h | Unpaired T-test | fig legend | lean n=12 mice<br><br>obese n=16 mice | # animals | fig legend | error bars are mean +/- SEM | p=0.0497 | fig legend | t=2.059, df=26 | fig legend | |
| 1j | two-way RM ANOVA followed by a Sidak's multiple comparisons test | fig legend | lean n=12 mice<br><br>obese n=13 mice | # animals | fig legend | error bars are mean +/- SEM | diet x devaluation interaction p=0.0106<br><br>diet effect p=0.0343<br><br>devaluation effect p=0.0074 | fig legend | diet x devaluation interaction F (1, 23) = 7.740<br><br>diet effect F (1, 23) = 5.066<br><br>devaluation effect F (1, 23) = 8.642 | fig legend | lean p=0.0012<br><br>obese p=0.9920 |
| 1k | Unpaired T-test | fig legend | lean n=12 mice<br><br>obese n=13 mice | # animals | fig legend | error bars are mean +/- SEM | p=0.0174 | fig legend | t=2.562, df=23 | fig legend | |

Supplemental Table 1. Statistics table.

| | | | | | | | | | | | |
|---|---|---|---|---|---|---|---|---|---|---|---|
| 1l | Unpaired T-test | fig legend | lean n=12 mice<br><br>obese n=13 mice | # animals | fig legend | error bars are mean +/- SEM | p=0.0343 | fig legend | t=2.251, df=23 | fig legend | |
| 1m | two-way RM ANOVA | fig legend | lean n=12 mice<br><br>obese n=13 mice | # animals | fig legend | error bars are mean +/- SEM | diet x consumption interaction p=0.7707<br><br>diet effect p=0.4773<br><br>consumption effect p=0.7707 | fig legend | diet x consumption interaction F (1, 23) = 0.08699<br><br>diet effect F (1, 23) = 0.5220<br><br>consumption effect F (1, 23) = 0.08699 | fig legend | |
| 1n | two-way RM ANOVA followed by a Sidak's multiple comparisons test | fig legend | lean n=12 mice<br><br>obese n=13 mice | # animals | fig legend | error bars are mean +/- SEM | diet x devaluation interaction p=0.9788<br><br>diet effect p=0.0566<br><br>devaluation effect p=0.0018 | fig legend | diet x devaluation interaction F (1, 23) = 0.0007210<br><br>diet effect F (1, 23) = 4.028<br><br>devaluation effect F (1, 23) = 12.37 | fig legend | lean p=0.0471<br><br>obese p=0.0349 |
| 1o | two-way RM ANOVA followed by a Sidak's multiple comparisons test | fig legend | lean n=12 mice<br><br>obese n=13 mice | # animals | fig legend | error bars are mean +/- SEM | diet x devaluation interaction p=0.8188<br><br>diet effect p=0.0006<br><br>devaluation effect p=0.0001 | fig legend | diet x devaluation interaction F (1, 23) = 0.05368<br><br>diet effect F (1, 23) = 15.72<br><br>devaluation effect F (1, 23) = 21.94 | fig legend | lean p=0.0104<br><br>obese p=0.0034 |
| 2b | two-way RM ANOVA followed by a Sidak's multiple comparisons test | fig legend | lean n=10 mice<br><br>obese n=12 mice | # animals | fig legend | error bars are mean +/- SEM | diet x time interaction p<0.0001<br><br>diet effect p<0.0001<br><br>time effect p<0.0001 | fig legend | diet x time interaction F (1, 20) = 49.20<br><br>diet effect F (1, 20) = 75.01<br><br>time effect F (1, 20) = 179.3 | fig legend | lean p=0.0007<br><br>obese p<0.0001 |
| 2c | 3-way RM ANOVA followed by a Tukey's multiple comparison test | fig legend | lean n=10 mice<br><br>obese n=12 mice | # animals | fig legend | error bars are mean +/- SEM | day x diet x devaluation interaction p=0.2593<br><br>diet x devaluation interaction p=0.5735<br><br>day x devaluation interaction p=0.0006<br><br>day x diet interaction p=0.8001 | fig legend | day x diet x devaluation interaction F (2, 40) = 1.396<br><br>diet x devaluation interaction F (1, 20) = 0.3276<br><br>day x devaluation interaction F (1.391, 27.82) = 12.33 | fig legend | Pairing day 3 lean valued vs devalued p=0.0379<br><br>Pairing day 3 obese valued vs devalued p=0.0015 |

Supplemental Table 1. Statistics table.

| | | | | | | | | | | | |
|---|---|---|---|---|---|---|---|---|---|---|---|
| | | | | | | | devaluation effect<br>p<0.0001<br><br>diet effect<br>p=0.8831<br><br>day effect<br>p=0.0003 | | day x diet interaction<br>F (2, 40) = 0.2243<br><br>devaluation effect<br>F (1.000, 20.00) = 24.42<br><br>diet effect<br>F (1, 20) = 0.02218<br><br>day effect<br>F (1.832, 36.63) = 10.84 | | |
| 2d | two-way RM ANOVA followed by a Sidak's multiple comparisons test | fig legend | lean n=10 mice<br><br>obese n=12 mice | # animals | fig legend | error bars are mean +/- SEM | diet x devaluation interaction<br>p=0.3401<br><br>diet effect<br>p=0.7297<br><br>devaluation effect<br>p<0.0001 | fig legend | diet x devaluation interaction<br>F (1, 20) = 0.9552<br><br>diet effect<br>F (1, 20) = 0.1227<br><br>devaluation effect<br>F (1, 20) = 60.98 | fig legend | lean<br>p=0.0003<br><br>obese<br>p<0.0001 |
| 2e | Unpaired t-test | fig legend | lean n=10 mice<br><br>obese n=12 mice | # animals | fig legend | error bars are mean +/- SEM | p=0.5060 | fig legend | t=0.6773, df=20 | fig legend | |
| 2f | Unpaired t-test | fig legend | lean n=10 mice<br><br>obese n=12 mice | # animals | fig legend | error bars are mean +/- SEM | p=0.7297 | fig legend | t=0.3503, df=20 | fig legend | |
| 2g | two-way RM ANOVA followed by a Sidak's multiple comparisons test | fig legend | lean n=10 mice<br><br>obese n=12 mice | # animals | fig legend | error bars are mean +/- SEM | diet x devaluation interaction<br>p=0.0004<br><br>diet effect<br>p=0.8229<br><br>devaluation effect<br>p=0.0096 | fig legend | diet x devaluation interaction<br>F (1, 20) = 17.92<br><br>diet effect<br>F (1, 20) = 0.05144<br><br>devaluation effect<br>F (1, 20) = 8.210 | fig legend | lean<br>p=0.0002<br><br>obese<br>p=0.5410 |
| 2h | Unpaired t-test | fig legend | lean n=10 mice<br><br>obese n=12 mice | # animals | fig legend | error bars are mean +/- SEM | p=0.0001 | fig legend | t=4.681, df=20 | fig legend | |
| 2i | Unpaired t-test | fig legend | lean n=10 mice<br><br>obese n=12 mice | # animals | fig legend | error bars are mean +/- SEM | p=0.8229 | fig legend | t=0.2268, df=20 | fig legend | |
| 3b | Mixed effects analysis | fig legend | lean n=10/3<br><br>lean PTX n=11/3<br><br>obese n=8/3<br><br>obese PTX n=9/5 | # cells / # animals | fig legend | error bars are mean +/- SEM | current x drug x diet interaction<br>p<0.0001<br><br>drug x diet interaction<br>p=0.0023<br><br>current x diet interaction<br>p=0.1156<br><br>current x drug interaction<br>p=0.0071 | fig legend | current x drug x diet interaction<br>F (20, 318) = 9.668<br><br>drug x diet interaction<br>F (1, 18) = 12.56<br><br>current x diet interaction<br>F (20, 360) = 1.406 | fig legend | |

Supplemental Table 1. Statistics table.

| | | | | | | | | | | | |
|---|---|---|---|---|---|---|---|---|---|---|---|
| | | | | | | | diet effect p=0.0521<br><br>drug effect p=0.0653<br><br>current effect <0.0001 | | current x drug interaction F (20, 360) = 1.406<br><br>diet effect F (1, 18) = 4.325<br><br>drug effect F (0.09746, 1.754) = 7.656<br><br>current effect F (20.00, 360.0) = 779.8 | | |
| 3c | two-way ANOVA followed by a Sidak's multiple comparisons test | fig legend | lean n=10/3<br><br>lean PTX n=11/3<br><br>obese n=8/3<br><br>obese PTX n=9/5 | # cells / # animals | fig legend | error bars are mean +/- SEM | drug x diet interaction p=0.0004<br><br>drug effect p=0.0114<br><br>diet effect p=0.3806 | fig legend | drug x diet interaction F (1, 34) = 15.14<br><br>drug effect F (1, 34) = 7.156<br><br>diet effect F (1, 34) = 0.7891 | fig legend | lean: no PTX vs. PTX p=0.0001<br><br>obese: no PTX vs. PTX: obese p=0.6627<br><br>lean no PTX vs obese no PTX p=0.0139 |
| 3e | Unpaired T-test | fig legend | lean n=10/6<br><br>obese n=9/3 | # cells / # animals | fig legend | error bars are mean +/- SEM | p=0.0091 | fig legend | t=2.940, df=17 | fig legend | |
| 3f | Unpaired T-test | fig legend | lean n=10/6<br><br>obese n=9/4 | # cells / # animals | fig legend | error bars are mean +/- SEM | p=0.0975 | fig legend | t=1.754, df=17 | fig legend | |
| 3h | Unpaired T-test | fig legend | lean n=11/4<br><br>obese n=11/4 | # cells / # animals | fig legend | error bars are mean +/- SEM | p=0.0375 | fig legend | t=2.229, df=20 | fig legend | |
| 3j | Unpaired T-test | fig legend | lean n=9/4<br><br>obese n=10/4 | # cells / # animals | fig legend | error bars are mean +/- SEM | p=0.0313 | fig legend | t=2.348, df=17 | fig legend | |
| 3k | Unpaired T-test | fig legend | lean n=9/4<br><br>obese n=10/4 | # cells / # animals | fig legend | error bars are mean +/- SEM | p=0.0420 | fig legend | t=2.199, df=17 | fig legend | |
| 4d | Paired T test | fig legend | n=6/2 | # cells / # animals | fig legend | error bars are mean +/- SEM | p=0.0189 | fig legend | t=3.418, df=5 | fig legend | |
| 4e | Three-way RM ANOVA followed by a Holm-Sidak's multiple comparison test | fig legend | mCherry n=7<br><br>hM4D(Gi) n=9 | # animals | fig legend | error bars are mean +/- SEM | devaluation x virus x drug interaction p=0.1223<br><br>virus x drug interaction p=0.9023<br><br>devaluation x drug interaction p=0.1374<br><br>devaluation x virus interaction p=0.0673<br><br>drug effect p=0.0754<br><br>virus effect p=0.9610<br><br>devaluation effect p=0.0008 | fig legend | devaluation x virus x drug interaction F (1, 14) = 2.705<br><br>virus x drug interaction F (1, 14) = 0.01563<br><br>devaluation x drug interaction F (1, 14) = 2.483<br><br>devaluation x virus interaction F (1, 14) = 3.933<br><br>drug effect F (1, 14) = 3.687<br><br>virus effect | fig legend | mCherry vehicle: valued vs. devalued p=0.0153<br><br>mCherry CNO: valued vs. devalued p=0.0153<br><br>hM4D(Gi) vehicle: valued vs. devalued p=0.0153<br><br>hM4D(Gi) CNO: valued vs. devalued p=0.7508 |

Supplemental Table 1. Statistics table.

| | | | | | | | | | | | |
|---|---|---|---|---|---|---|---|---|---|---|---|
| | | | | | | | | | F (1, 14) = 0.002476<br><br>devaluation effect<br>F (1, 14) = 17.96 | | |
| 4f | two-way RM ANOVA followed by a Sidak's multiple comparisons test | fig legend | mCherry n=7<br><br>hM4D(Gi) n=9 | # animals | fig legend | error bars are mean +/- SEM | virus vs drug interaction p=0.0070<br><br>virus effect p=0.0008<br><br>drug effect p=0.0003 | fig legend | virus vs drug interaction F (1, 14) = 9.984<br><br>virus effect F (1, 14) = 18.07<br><br>drug effect F (1, 14) = 22.30 | fig legend | mCherry vehicle vs CNO p=0.5311<br><br>hM4D(Gi) vehicle vs CNO p<0.0001 |
| 4g | two-way RM ANOVA followed by a Sidak's multiple comparisons test | fig legend | mCherry n=7<br><br>hM4D(Gi) n=9 | # animals | fig legend | error bars are mean +/- SEM | virus vs drug interaction p=0.9023<br><br>virus effect p=0.9610<br><br>drug effect p=0.0754 | fig legend | virus x drug interaction F (1, 14) = 0.01563<br><br>virus effect F (1, 14) = 0.002476<br><br>drug effect F (1, 14) = 3.687 | fig legend | mCherry vehicle vs CNO p=0.4394<br><br>hM4D(Gi) vehicle vs CNO p=0.2679 |
| 4h | Three-way RM ANOVA followed by a Holm-Sidak's multiple comparisons test | fig legend | mCherry n=12<br><br>hM4D(Gi) n=13 | # animals | fig legend | error bars are mean +/- SEM | devaluation x virus x drug interaction p=0.0406<br><br>virus x drug interaction p=0.5552<br><br>devaluation x drug interaction p=0.0442<br><br>devaluation x virus interaction p=0.0110<br><br>drug effect p=0.0096<br><br>virus effect p=0.7288<br><br>devaluation effect p=0.0001 | fig legend | devaluation x virus x drug interaction F (1, 23) = 4.709<br><br>virus x drug interaction F (1, 23) = 0.3585<br><br>devaluation x drug interaction F (1.000, 23.00) = 4.533<br><br>devaluation x virus interaction F (1, 23) = 7.653<br><br>drug effect F (1.000, 23.00) = 7.989<br><br>virus effect F (1, 23) = 0.1232<br><br>devaluation effect F (1.000, 23.00) = 20.85 | fig legend | mCherry vehicle: valued vs devalued p=0.0009<br><br>mCherry CNO: valued vs devalued p=0.0180<br><br>hM4D(Gi) vehicle: valued vs. devalued p=0.0019<br><br>hM4D(Gi) CNO: valued vs. devalued p=0.5752 |
| 4i | two-way ANOVA | fig legend | mCherry n=12<br><br>hM4D(Gi) n=13 | # animals | fig legend | error bars are mean +/- SEM | virus vs drug interaction p=0.0256<br><br>virus effect p=0.0008<br><br>drug effect p<0.0001 | fig legend | virus vs drug interaction F (1, 23) = 5.700<br><br>virus effect F (1, 23) = 15.04<br><br>drug effect F (1, 23) = 26.44 | fig legend | mCherry vehicle vs CNO p=0.1327<br><br>hM4D(Gi) vehicle vs CNO p<0.0001 |

Supplemental Table 1. Statistics table.

| | | | | | | | | | | | |
|---|---|---|---|---|---|---|---|---|---|---|---|
| 4j | two-way ANOVA | fig legend | mCherry n=12<br><br>hM4D(Gi) n=13 | # animals | fig legend | error bars are mean +/- SEM | virus vs drug interaction p=0.5552<br><br>virus effect p=0.7288<br><br>drug effect p=0.0096 | fig legend | virus vs drug interaction F (1, 23) = 0.3585<br><br>virus effect F (1, 23) = 0.1232<br><br>drug effect F (1, 23) = 7.989 | fig legend | mCherry vehicle vs CNO p=0.0519<br><br>hM4D(Gi) vehicle vs CNO p=0.2283 |
| 5b | Mixed effects analysis | fig legend | lean vehicle n=7/4<br><br>lean NNC-711 n=7/4<br><br>obese vehicle n=10/4<br><br>obese NNC-711 n=10/6 | # cells / # animals | fig legend | error bars are mean +/- SEM | current x drug x diet interaction p=0.0253<br><br>drug x diet interaction p=0.1196<br><br>current x diet interaction p=0.0190<br><br>current x drug interaction p=0.0127<br><br>diet effect p=0.0715<br><br>drug effect p=0.0171<br><br>current effect p<0.0001 | fig legend | current x drug x diet interaction F (20, 278) = 1.756<br><br>drug x diet interaction F (1, 278) = 2.438<br><br>current x diet interaction F (20, 278) = 1.816<br><br>current x drug interaction F (1.192, 16.56) = 7.192<br><br>diet effect F (1, 278) = 3.272<br><br>drug effect F (1.000, 16.00) = 7.076<br><br>current effect F (2.154, 34.46) = 181.3 | fig legend | |
| 5c | two-way ANOVA | fig legend | lean vehicle n=7/4<br><br>lean NNC-711 n=7/4<br><br>obese vehicle n=10/4<br><br>obese NNC-711 n=10/6 | # cells / # animals | fig legend | error bars are mean +/- SEM | drug x diet interaction p=0.1282<br><br>drug effect p=0.0031<br><br>diet effect p=0.1552 | fig legend | drug x diet interaction F (1, 30) = 2.447<br><br>drug effect F (1, 30) = 10.32<br><br>diet effect F (1, 30) = 2.126 | fig legend | Lean: vehicle vs NNC-711 p=0.4977<br><br>Obese: vehicle vs NNC-711 p=0.0016 |
| 5e | two-way ANOVA | fig legend | lean vehicle n=5/4<br><br>lean NNC-711 n=6/6<br><br>obese vehicle n=5/23<br><br>obese NNC-711 n=8/5 | # cells / # animals | fig legend | error bars are mean +/- SEM | drug x diet interaction p=0.1836<br><br>drug effect p=0.1016<br><br>diet effect p=0.0172 | fig legend | drug x diet interaction F (1, 20) = 1.898<br><br>drug effect F (1, 20) = 2.944<br><br>diet effect F (1, 20) = 6.743 | fig legend | vehicle lean vs obese p=0.0330<br><br>NNC-711 lean vs obese p=0.5892 |
| 5f | two-way ANOVA | fig legend | lean vehicle n=5/4<br><br>lean NNC-711 n=6/6 | # cells / # animals | fig legend | error bars are mean +/- SEM | drug x diet interaction p=0.2202<br><br>drug effect p=0.8707 | fig legend | drug x diet interaction F (1, 20) = 1.602<br><br>drug effect | fig legend | |

Supplemental Table 1. Statistics table.

| | | | | | | | | | | | |
|---|---|---|---|---|---|---|---|---|---|---|---|
| | | | obese vehicle<br>n=5/23<br><br>obese NNC-711<br>n=8/5 | | | | diet effect<br>p=0.8219 | | F (1, 20) = 0.02718<br><br>diet effect<br>F (1, 20) = 0.05205 | | |
| 5h | | fig legend | | # cells / # animals | fig legend | error bars are mean +/- SEM | | fig legend | | fig legend | |
| 5i | Unpaired T-test | fig legend | lean n=7/6<br><br>obese n=8/6 | # cells / # animals | fig legend | error bars are mean +/- SEM | p=0.0023 | fig legend | t=3.786, df=13 | fig legend | |
| 5j | two-way ANOVA followed by Holms-Sidak's multiple comparisons | fig legend | lean GBZ<br>n=8/4<br><br>lean GBZ + NNC-711<br>n=7/4<br><br>obese GBZ<br>n=9/5<br><br>obese GBZ + NNC-711<br>n=7/4 | # cells / # animals | fig legend | error bars are mean +/- SEM | drug x diet interaction<br>p=0.7144<br><br>drug effect<br>p=0.0013<br><br>diet effect<br>p=0.0095 | fig legend | drug x diet interaction<br>F (1, 27) = 0.1367<br><br>drug effect<br>F (1, 27) = 12.90<br><br>diet effect<br>F (1, 27) = 7.786 | fig legend | Lean: GBZ vs NNC+ GBZ<br>p=0.0202<br><br>obese: GBZ vs NNC+ GBZ<br>p=0.0288 |
| 5k | two-way ANOVA followed by Holms-Sidak's multiple comparisons | fig legend | lean GBZ<br>n=8/4<br><br>lean GBZ + NNC-711<br>n=7/4<br><br>obese GBZ<br>n=9/5<br><br>obese GBZ + NNC-711<br>n=7/4 | # cells / # animals | fig legend | error bars are mean +/- SEM | drug x diet interaction<br>p=0.0898<br><br>drug effect<br>p=0.0011<br><br>diet effect<br>p=0.0643 | fig legend | drug x diet interaction<br>F (1, 27) = 3.095<br><br>drug effect<br>F (1, 27) = 13.34<br><br>diet effect<br>F (1, 27) = 3.721 | fig legend | Lean: GBZ vs NNC+ GBZ<br>p=0.0016<br><br>obese: GBZ vs NNC+ GBZ<br>p=0.1860 |
| 6c | three-way ANOVA mixed effects model followed by Holm Sidak's multiple comparisons | fig legend | lean vehicle<br>n=12<br><br>lean NNC-711<br>n= 12<br><br>obese vehicle<br>n=14<br><br>obese NNC-711<br>n=15 | # animals | fig legend | error bars are mean +/- SEM | devaluation x diet x drug interaction<br>p=0.6634<br><br>diet x drug interaction<br>p=0.9273<br><br>devaluation x drug interaction<br>p=0.0211<br><br>devaluation x diet interaction<br>p=0.0020<br><br>drug effect<br>p=0.3446<br><br>diet effect<br>p=0.0586<br><br>devaluation effect<br>p<0.0001 | fig legend | devaluation x diet x drug interaction<br>F (1, 14) = 0.1977<br><br>diet x drug interaction<br>F (1, 14) = 0.008637<br><br>devaluation x drug interaction<br>F (1.000, 14.00) = 6.739<br><br>devaluation x diet interaction<br>F (1, 14) = 14.34<br><br>drug effect<br>F (1.000, 28.00) = 0.9240<br><br>diet effect<br>F (1, 14) = 4.241<br><br>devaluation effect<br>F (1.000, 28.00) = 26.07 | fig legend | lean vehicle valued vs. devalued<br>p=0.0142<br><br>lean NNC-711 valued vs. devalued<br>p=0.0204<br><br>obese vehicle valued vs. devalued<br>p=0.1309<br><br>obese NNC-711 valued vs. devalued<br>p=0.0201 |
| 6d | Two-way ANOVA mixed effects model followed by a Sidak's | fig legend | lean vehicle<br>n=12<br><br>lean NNC-711<br>n= 12 | # animals | fig legend | error bars are mean +/- SEM | diet x drug interaction<br>p=0.0565<br><br>drug effect<br>p=0.0047 | fig legend | diet x drug interaction<br>F (1, 21) = 4.074<br><br>drug effect | fig legend | Lean vehicle vs NNC-711<br>P=0.6948<br>Obese vehicle vs NNC-711<br>P=0.0018 |

Supplemental Table 1. Statistics table.

| | | | | | | | | | | | |
|---|---|---|---|---|---|---|---|---|---|---|---|
| | multiple comparisons test | | obese vehicle n=14<br><br>obese NNC-711 n=15 | | | | diet effect p=0.0008 | | F (1, 21) = 10.02<br><br>diet effect F (1, 28) = 14.23 | | |
| 6e | Two-way ANOVA mixed effects model | fig legend | lean vehicle n=12<br><br>lean NNC-711 n= 12<br><br>obese vehicle n=14<br><br>obese NNC-711 n=15 | # animals | fig legend | error bars are mean +/- SEM | diet x drug interaction p= 0.9194<br><br>drug effect p=0.3119<br><br>diet effect p=0.0484* | fig legend | diet x drug interaction F (1, 21) = 0.01049<br><br>drug effect F (1, 21) = 1.074<br><br>diet effect F (1, 28) = 4.262 | fig legend | |
| 7b | three-way RM ANOVA | fig legend | lean n=9/4<br><br>obese n=7/4 | # cells / # animals | fig legend | error bars are mean +/- SEM | current x stim x diet interaction p=0.5513<br><br>stim x diet interaction p=0.3317<br><br>current x diet interaction p=0.0784<br><br>current x stim interaction p<0.0001<br><br>diet effect p=0.0667<br><br>stim effect p=0.0005<br><br>current effect p<0.0001 | fig legend | current x stim x diet interaction F (20, 280) = 0.9284<br><br>stim x diet interaction F (1, 14) = 1.011<br>current x diet interaction F (20, 280) = 1.505<br><br>current x stim interaction F (2.535, 35.49) = 10.52<br><br>diet effect F (1, 14) = 3.954<br><br>stim effect F (1.000, 14.00) = 19.88<br><br>current effect F (4.154, 58.16) = 102.5 | fig legend | |
| 7c | two-way RM ANOVA followed by a Sidak's multiple comparisons test | fig legend | lean n=9/4<br><br>obese n=7/4 | # cells / # animals | fig legend | error bars are mean +/- SEM | diet x stimulation interaction p=0.1655<br><br>diet effect p=0.2571<br><br>stimulation effect p<0.0001 | fig legend | diet x stimulation interaction F (1, 14) = 2.141<br><br>diet effect F (1, 14) = 1.396<br><br>stimulation effect F (1, 14) = 33.58 | fig legend | lean baseline vs. stim p=0.0110<br><br>obese baseline vs. stim p=0.0005 |
| 7e | two-way RM ANOVA followed by a Sidak's multiple comparisons test | fig legend | lean n=7/4<br><br>obese n=11/5 | # cells / # animals | fig legend | error bars are mean +/- SEM | diet x stimulation interaction p=0.7657<br><br>diet effect p=0.0049<br><br>stimulation effect p=0.8894 | fig legend | diet x stimulation interaction F (1, 16) = 0.09190<br><br>diet effect F (1, 16) = 10.66<br><br>stimulation effect F (1, 16) = 0.01998 | fig legend | baseline lean vs obese p=0.0095<br><br>stimulation lean vs obese p=0.0060 |

Supplemental Table 1. Statistics table.

| | | | | | | | | | | | |
|---|---|---|---|---|---|---|---|---|---|---|---|
| 7f | two-way RM ANOVA | fig legend | lean n=7/4<br><br>obese n=11/5 | # cells / # animals | fig legend | error bars are mean +/- SEM | diet x stimulation interaction p=0.5285<br><br>diet effect p=0.0272<br><br>stimulation effect p=0.2726 | fig legend | diet x stimulation interaction F (1, 16) = 0.4151<br><br>diet effect F (1, 16) = 5.911<br>stimulation effect F (1, 16) = 1.291 | fig legend | |
| 7h | two-way RM ANOVA followed by a Sidak's multiple comparisons test | fig legend | lean n=8/6<br><br>obese n=7/4 | # cells / # animals | fig legend | error bars are mean +/- SEM | diet x stimulation interaction p=0.1609<br><br>diet effect p=0.0725<br><br>stimulation effect p=0.0024 | fig legend | diet x stimulation interaction F (1, 13) = 2.211<br><br>diet effect F (1, 13) = 3.819<br><br>stimulation effect F (1, 13) = 14.12 | fig legend | baseline lean vs obese p=0.0462<br><br>stimulation lean vs obese p=0.4828 |
| 7i | Unpaired T-test | fig legend | lean n=8/6<br><br>obese n=7/4 | # cells / # animals | fig legend | error bars are mean +/- SEM | p= 0.0299 | fig legend | t=2.438, df=13 | fig legend | |
| 8d | three-way ANOVA mixed effects model followed by Holm Sidak's multiple comparisons | fig legend | Lean 589nM n=8<br><br>lean 473nM n=7<br><br>obese 589nM n=6<br><br>obese 473nM n=7 | # animals | fig legend | error bars are mean +/- SEM | devaluation x diet x stim interaction p=0.1399<br><br>diet x stim interaction p=0.5306<br><br>devaluation x stim interaction p=0.1818<br><br>devaluation x diet interaction p=0.0427<br><br>stim effect p=0.5421<br><br>diet effect p=0.2384<br><br>devaluation effect p=0.0004 | fig legend | devaluation x diet x stim interaction F (1, 9) = 2.621<br><br>diet x stim interaction F (1, 9) = 0.4254<br><br>devaluation x stim interaction F (1, 9) = 2.094<br><br>devaluation x diet interaction F (1, 9) = 5.564<br><br>stim effect F (1, 13) = 0.3920<br><br>diet effect F (1, 9) = 1.594<br><br>devaluation effect F (1, 13) = 21.95 | fig legend | lean 589nM stimulation valued vs. devalued: p=0.0149<br><br>lean 473nM stimulation valued vs. devalued: p=0.0215<br><br>obese 589nM stimulation valued vs. devalued: p=0.7385<br><br>obese 473nM stimulation valued vs. devalued: p=0.0474 |
| 8e | two-way ANOVA mixed effects followed by Sidak's multiple comparison | fig legend | lean 589 n=8<br><br>lean 473 n=7<br><br>obese 589 n=6<br><br>obese 473 n=7 | # animals | fig legend | error bars are mean +/- SEM | diet x stimulation interaction p=0.0001<br><br>diet effect p=0.0621<br><br>stimulation effect p=0.0012 | fig legend | diet x stimulation interaction F (1, 24) = 20.29<br><br>diet effect F (1, 24) = 3.829<br><br>stimulation effect F (1, 24) = 13.61 | fig legend | 589nM stimulation lean vs obese p=0.0003<br><br>473nM stimulation lean vs obese p=0.1586 |

Supplemental Table 1. Statistics table.

| | | | | | | | | | | |
|---|---|---|---|---|---|---|---|---|---|---|
| 8f | two-way ANOVA mixed effects | fig legend | Lean 589nM n=8<br><br>lean 473nM n=7<br><br>obese 589nM n=6<br><br>obese 473nM n=7 | # animals | fig legend | error bars are mean +/- SEM | diet x stimulation interaction p=0.5277<br><br>diet effect p=0.2289<br><br>stimulation effect p=0.5440 | fig legend | diet x stimulation interaction F (1, 11) = 0.4254<br><br>diet effect F (1, 13) = 1.594<br><br>stimulation effect F (1, 11) = 0.3920 | fig legend |

Extended Data Statistics

| V | Test Used | | n | | | Descriptive statistics (Average, Variance) | p - Value | | Degrees of freedom & F/t/z/R | | Multiple comparisons |
|---|---|---|---|---|---|---|---|---|---|---|---|
| Figure # | Test | Section & paragraph # | Exact devaluation | Defined | Section & paragraph # | Reported | Exact devaluation | Section & paragraph # | Value | Section & paragraph # | |
| ED 1a | two-way ANOVA followed by Sidaks multiple comparison | fig legend | lean ad lib n=8<br><br>lean food restricted n=9<br><br>obese ad libitum n=8<br><br>obese food restricted n=8 | # animals | fig legend | error bars are mean +/- SEM | feeding schedule x diet interaction p=0.1068<br><br>feeding schedule effect p=0.2048<br><br>diet effect p<0.0001 | fig legend | feeding schedule x diet interaction F (1, 29) = 2.771<br><br>feeding schedule effect F (1, 29) = 1.682<br><br>diet effect F (1, 29) = 92.08 | fig legend | Food restricted lean vs obese p<0.0001<br><br>Ad libitum lean vs obese p<0.0001 |
| ED 1b | 3-Way ANOVA | fig legend | lean food restricted n=9<br><br>lean ad lib n=8<br><br>obese food restricted n=8<br><br>obese ad libitum n=8 | # animals | fig legend | error bars are mean +/- SEM | time x diet x feeding schedule interaction p=0.0359<br><br>diet x feeding schedule interaction p=0.4830<br><br>time x feeding schedule interaction p=0.0635<br><br>time x diet interaction p<0.0001<br><br>feeding schedule effect p=0.1872<br><br>diet effect p<0.0001<br><br>time effect p<0.0001 | fig legend | time x diet x feeding schedule interaction F(11, 153) = 1.962<br><br>diet x feeding schedule interaction F(1, 153) = 0.4945<br><br>time x feeding schedule interaction F(11, 153) = 2.935<br><br>time x diet interaction F (11, 153) = 10.72<br><br>feeding schedule effect F (1, 15) = 1.910<br><br>diet effect F (1, 153) = 27.63<br><br>time effect | fig legend | |

Supplemental Table 1. Statistics table.

| | | | | | | | | | | | |
|---|---|---|---|---|---|---|---|---|---|---|---|
| | | | | | | | | | F (11, 165) = 90.76 | | |
| ED 1c | two-way ANOVA followed by Sidaks multiple comparison | fig legend | lean food restricted n=9<br><br>lean ad lib n=8<br><br>obese food restricted n=8<br><br>obese ad libitum n=8 | # animals | fig legend | error bars are mean +/- SEM | feeding schedule x diet interaction p=0.8210<br><br>feeding schedule effect p=0.7503<br><br>diet effect p=0.0027 | fig legend | feeding schedule x diet interaction F (1, 29) = 0.05216<br><br>feeding schedule effect F (1, 29) = 0.1033<br><br>diet effect F (1, 29) = 10.73 | fig legend | |
| ED 1d | two-way ANOVA followed by Sidaks multiple comparison | fig legend | lean food restricted n=9<br><br>lean ad lib n=8<br><br>obese food restricted n=8<br><br>obese ad libitum n=8 | # animals | fig legend | error bars are mean +/- SEM | feeding schedule x diet interaction p=0.4785<br><br>feeding schedule effect p=0.1531<br><br>diet effect p<0.0001 | fig legend | feeding schedule x diet interaction F (1, 29) = 0.5155<br><br>feeding schedule effect F (1, 29) = 2.152<br><br>diet effect F (1, 29) = 27.62 | fig legend | Food restricted lean vs obese p=0.0058<br><br>Ad libitum lean vs obese p=0.0005 |
| ED 2a | two-way ANOVA followed by Dunnett's multiple comparison | fig legend | lean n=60<br><br>obese n=60 | # animals | fig legend | error bars are mean +/- SEM | training schedule x diet interaction p=0.0010<br><br>training schedule effect p<0.0001<br><br>diet effect p<0.0001 | fig legend | training schedule x diet interaction F (2, 236) = 7.112<br><br>training schedule effect F (1.310, 154.6) = 18.26<br><br>diet effect F (1, 118) = 17.04 | fig legend | lean RR5 vs RR10 p<0.0001<br><br>lean RR5 vs RR20 p=0.0006<br><br>obese RR5 vs RR10 p=0.0073<br><br>obese RR5 vs RR20 p=0.0046 |
| ED 2c | Unpaired T test | fig legend | lean n=35<br><br>obese n=31 | # animals | fig legend | error bars are mean +/- SEM | p<0.0001 | fig legend | t=10.88, df=64 | fig legend | |
| ED 2d | Unpaired T test | fig legend | lean n=35<br><br>obese n=31 | # animals | fig legend | error bars are mean +/- SEM | p=0.5239 | fig legend | t=0.6408, df=64 | fig legend | |
| ED 2e | two-way RM ANOVA followed by Sidaks multiple comparison | fig legend | lean n=35<br><br>obese n=31 | # animals | fig legend | error bars are mean +/- SEM | diet x value interaction p=0.0187<br><br>diet effect p=0.0638<br><br>value effect p=0.0067 | fig legend | diet x value interaction F (1, 64) = 5.827<br><br>diet effect F (1, 64) = 3.558<br><br>value effect F (1, 64) = 7.853 | fig legend | lean valued vs devalued p=0.0006<br><br>obese valued vs devalued P=0.9561 |
| ED 2f | Unpaired T test | fig legend | lean n=35<br><br>obese n=31 | # animals | fig legend | error bars are mean +/- SEM | p=0.0001 | fig legend | t=4.109, df=64 | fig legend | |
| ED 2g | Unpaired T test | fig legend | lean n=35 | # animals | fig legend | error bars are mean +/- SEM | P=0.0638 | fig legend | t=1.886, df=64 | fig legend | |

Supplemental Table 1. Statistics table.

| | | | | | | | | | | | |
|---|---|---|---|---|---|---|---|---|---|---|---|
| | | | obese n=31 | | | | | | | | |
| ED 2h | Unpaired T test | fig legend | lean n=12<br><br>obese n=16 | # animals | fig legend | error bars are mean +/- SEM | p<0.0001 | fig legend | t=10.18, df=26 | fig legend | |
| ED 2i | Unpaired T test | fig legend | lean n=12<br><br>obese n=16 | # animals | fig legend | error bars are mean +/- SEM | p=0.7472 | fig legend | t=0.3258, df=26 | fig legend | |
| ED 2j | Unpaired T test | fig legend | lean n=12<br><br>obese n=13 | # animals | fig legend | error bars are mean +/- SEM | p<0.0001 | fig legend | t=6.068, df=23 | fig legend | |
| ED 3b | two-way RM ANOVA followed by Sidaks multiple comparison | fig legend | lean n=8<br><br>obese n=13 | # animals | fig legend | error bars are mean +/- SEM | diet x value interaction p=0.1274<br><br>diet effect p=0.3331<br><br>value effect p=0.0331 | fig legend | diet x value interaction F (1, 19) = 2.541<br><br>diet effect F (1, 19) = 0.9864<br><br>value effect F (1, 19) = 5.283 | fig legend | lean valued vs devalued p=0.0454<br><br>obese valued vs devalued P=0.8193 |
| ED 3c | Unpaired T test | fig legend | lean n=8<br><br>obese n=13 | # animals | fig legend | error bars are mean +/- SEM | p=0.0226 | fig legend | t=2.481, df=19 | fig legend | |
| ED 3d | Unpaired T test | fig legend | lean n=8<br><br>obese n=13 | # animals | fig legend | error bars are mean +/- SEM | p=0.3331 | fig legend | t=0.9932, df=19 | fig legend | |
| ED 3e | Unpaired T test | fig legend | lean n=15<br><br>obese n=11 | # animals | fig legend | error bars are mean +/- SEM | p<0.0001 | fig legend | t=13.25, df=24 | fig legend | |
| ED 3f | Unpaired T test | fig legend | lean n=15<br><br>obese n=11 | # animals | fig legend | error bars are mean +/- SEM | p=0.1412 | fig legend | t=1.521, df=24 | fig legend | |
| ED 3g | Unpaired T test | fig legend | lean n=15<br><br>obese n=11 | # animals | fig legend | error bars are mean +/- SEM | p=0.2122 | fig legend | t=1.282, df=24 | fig legend | |
| ED 3h | Unpaired T test | fig legend | lean n=15<br><br>obese n=11 | # animals | fig legend | error bars are mean +/- SEM | p=0.2293 | fig legend | t=1.234, df=24 | fig legend | |
| ED 3i | two-way RM ANOVA followed by Sidaks multiple comparison | fig legend | lean n=15<br><br>obese n=16 | # animals | fig legend | error bars are mean +/- SEM | contingency x diet interaction p=0.1982<br><br>contingency effect p=0.0001<br><br>diet effect p=0.2195 | fig legend | contingency x diet interaction F (1, 29) = 1.734<br><br>contingency effect F (1, 29) = 19.97<br><br>diet effect F (1, 29) = 1.575 | fig legend | lean ND vs CD p=0.0007<br><br>obese ND vs CD p=0.0612 |
| ED 4a | two-way RM ANOVA followed by Sidaks multiple comparison | fig legend | lean n=8<br><br>obese n=9 | # animals | fig legend | error bars are mean +/- SEM | diet x value interaction p=0.9458<br><br>diet effect p=0.9410<br><br>value effect p<0.0001 | fig legend | diet x value interaction F (1, 15) = 0.004780<br><br>diet effect F (1, 15) = 0.005657<br><br>value effect F (1, 15) = 77.09 | fig legend | lean valued vs devalued p<0.0001<br><br>obese valued vs devalued p<0.0001 |
| ED 4b | Unpaired T test | fig legend | lean n=8<br><br>obese n=9 | # animals | fig legend | error bars are mean +/- SEM | p=0.9410 | fig legend | t=0.07521, df=15 | fig legend | |

Supplemental Table 1. Statistics table.

| | | | | | | | | | | | |
|---|---|---|---|---|---|---|---|---|---|---|---|
| ED 4c | Unpaired T test | fig legend | lean n=8<br><br>obese n=9 | # animals | fig legend | error bars are mean +/- SEM | p=0.0309 | fig legend | t=2.381, df=15 | fig legend | |
| ED 5a | two-way RM ANOVA followed by Sidaks multiple comparison | fig legend | lean n=9<br><br>obese n=8 | # animals | fig legend | error bars are mean +/- SEM | diet x time interaction p=0.0452<br><br>diet effect p<0.0001<br><br>time effect p=0.0025 | fig legend | diet x time interaction F (1, 15) = 4.772<br><br>diet effect F (1, 15) = 37.03<br><br>time effect F (1, 15) = 13.08 | fig legend | lean pre vs post p<0.0001<br><br>obese pre vs post p<0.0001 |
| ED 5b | two-way RM ANOVA followed by Dunnets multiple comparisons and simple linear regression | fig legend | lean n=9<br><br>obese n=8 | # animals | fig legend | error bars are mean +/- SEM | training schedule x diet interaction p=0.6820<br><br>training schedule effect p=0.0710<br><br>diet effect p=0.0011 | fig legend | training schedule x diet interaction F (2, 30) = 0.3876<br><br>training schedule effect F (2, 30) = 2.892<br><br>diet effect F (1, 15) = 16.28 | fig legend | lean RR5 vs RR10 p=0.5563<br><br>lean RR5 vs RR20 p=0.0448<br><br>obese RR5 vs RR10 p=0.9443<br><br>obese RR5 vs RR20 p=0.9443<br><br>slopes p=0.4560<br><br>y-intercept p<0001 |
| ED 5c | Unpaired T test | fig legend | lean n=9<br><br>obese n=8 | # animals | fig legend | error bars are mean +/- SEM | p=0.0335 | fig legend | t=2.341, df=15 | fig legend | |
| ED 5d | two-way RM ANOVA followed by Sidaks multiple comparison | fig legend | lean n=9<br><br>obese n=8 | # animals | fig legend | error bars are mean +/- SEM | diet x value interaction p=0.0035<br><br>diet effect p=0.0403<br><br>value effect p=0.0589 | fig legend | diet x value interaction F (1, 15) = 11.97<br><br>diet effect F (1, 15) = 5.038<br><br>value effect F (1, 15) = 4.178 | fig legend | lean valued vs devalued p=0.0023<br><br>obese valued vs devalued p=0.5726 |
| ED 5e | Unpaired T test | fig legend | lean n=9<br><br>obese n=8 | # animals | fig legend | error bars are mean +/- SEM | p=0.0256 | fig legend | t=2.478, df=15 | fig legend | |
| ED 5f | Unpaired T test | fig legend | lean n=9<br><br>obese n=8 | # animals | fig legend | error bars are mean +/- SEM | p=0.0403 | fig legend | t=2.245, df=15 | fig legend | |
| ED 6b | two-way ANOVA followed by Sidaks multiple comparison | fig legend | lean food restricted n=3<br><br>lean ad lib n=3<br><br>obese food restricted n=3<br><br>obese ad libitum | # animals | fig legend | error bars are mean +/- SEM | feeding schedule x diet interaction p=0.5997<br><br>feeding schedule effect p=0.0070<br><br>diet effect p=0.0070 | fig legend | feeding schedule x diet interaction F (1, 8) = 0.2985<br><br>feeding schedule effect F (1, 8) = 12.93<br><br>diet effect F (1, 8) = 34.16 | fig legend | Food restricted lean vs obese p=0.0039<br><br>Ad libitum lean vs obese p=0.0113 |

Supplemental Table 1. Statistics table.

| | | | | | | | | | | | |
|---|---|---|---|---|---|---|---|---|---|---|---|
| | | | n=3 | | | | | | | | |
| ED 6c | three-way RM ANOVA | fig legend | lean food restricted n=14/3<br><br>lean ad lib n=12/3<br><br>obese food restricted n=14/3<br><br>obese ad libitum n=12/3 | # cells / # animals | fig legend | error bars are mean +/- SEM | current x diet x food schedule interaction p=0.4906<br><br>diet x food schedule interaction p=0.7359<br><br>current x food schedule interaction p=0.0193<br><br>current x diet interaction p=<0.0001<br><br>food schedule effect p=0.0447<br><br>diet effect p=0.0005<br><br>current effect p<0.0001 | fig legend | current x diet x food schedule interaction F (20, 436) = 0.9759<br><br>diet x food schedule interaction F (1, 436) = 0.1139<br><br>current x food schedule interaction F (20, 436) = 1.792<br><br>current x diet interaction F (20, 436) = 13.45<br><br>food schedule effect F (1, 26) = 4.450<br><br>diet effect F (1, 436) = 12.28<br><br>current effect F (20, 520) = 509.4 | fig legend | |
| ED 6d | two-way RM ANOVA followed by Sidaks multiple comparison | fig legend | lean food restricted n=14/3<br><br>lean ad lib n=12/3<br><br>obese food restricted n=14/3<br><br>obese ad libitum n=12/3 | # cells / # animals | fig legend | error bars are mean +/- SEM | feeding schedule x diet interaction p=0.2983<br><br>feeding schedule effect p=0.3244<br><br>diet effect p<0.0001 | fig legend | feeding schedule x diet interaction F (1, 48) = 1.106<br><br>feeding schedule effect F (1, 48) = 0.9912<br><br>diet effect F (1, 48) = 20.85 | fig legend | Food restricted lean vs obese p=0.0254<br><br>Ad libitum lean vs obese p=0.0007 |
| ED 6e | two-way RM ANOVA | fig legend | lean food restricted n=14/3<br><br>lean ad lib n=12/3<br><br>obese food restricted n=14/3<br><br>obese ad libitum n=12/3 | # cells / # animals | fig legend | error bars are mean +/- SEM | feeding schedule x diet interaction p=0.5910<br><br>feeding schedule effect p=0.8250<br><br>diet effect p=0.0168 | fig legend | feeding schedule x diet interaction F (1, 48) = 0.2927<br><br>feeding schedule effect F (1, 48) = 0.04941<br><br>diet effect F (1, 48) = 6.144 | fig legend | |
| ED 7a | two-way ANOVA | fig legend | lean n=10/3<br><br>lean PTX n=11/3<br><br>obese n=8/3<br><br>obese PTX | # cells / # animals | fig legend | error bars are mean +/- SEM | drug x diet interaction p=0.0397<br><br>drug effect p=0.0853<br><br>diet effect | fig legend | drug x diet interaction F (1, 34) = 4.573<br><br>drug effect F (1, 34) = 3.141<br><br>diet effect F (1, 34) = 0.3277 | fig legend | |

Supplemental Table 1. Statistics table.

| | | | | | | | | | | | |
|---|---|---|---|---|---|---|---|---|---|---|---|
| | | | n=9/5 | | | | p=0.5708 | | | | |
| ED 7b | two-way ANOVA | fig legend | lean<br>n=10/3<br><br>lean PTX<br>n=11/3<br><br>obese<br>n=8/3<br><br>obese PTX<br>n=9/5 | # cells / # animals | fig legend | error bars are mean +/- SEM | drug x diet interaction<br>p=0.3836<br><br>drug effect<br>p=0.9308<br><br>diet effect<br>p=0.4124 | fig legend | drug x diet interaction<br>F (1, 34) = 0.7791<br><br>drug effect<br>F (1, 34) = 0.007646<br><br>diet effect<br>F (1, 34) = 0.6887 | fig legend | |
| ED 7c | two-way ANOVA followed by Sidaks multiple comparison | fig legend | lean<br>n=10/3<br><br>lean PTX<br>n=11/3<br><br>obese<br>n=8/3<br><br>obese PTX<br>n=9/5 | # cells / # animals | fig legend | error bars are mean +/- SEM | drug x diet interaction<br>p=0.0270<br><br>drug effect<br>p=0.1585<br><br>diet effect<br>p=0.0268 | fig legend | drug x diet interaction<br>F (1, 34) = 5.340<br><br>drug effect<br>F (1, 34) = 2.079<br><br>diet effect<br>F (1, 34) = 5.360 | fig legend | No PTX lean vs obese<br>p=0.0062<br><br>PTX lean vs obese<br>p>0.9999 |
| ED 7d | two-way ANOVA followed by Sidaks multiple comparison | fig legend | lean<br>n=10/3<br><br>lean PTX<br>n=11/3<br><br>obese<br>n=8/3<br><br>obese PTX<br>n=9/5 | # cells / # animals | fig legend | error bars are mean +/- SEM | drug x diet interaction<br>p=0.0489<br><br>drug effect<br>p=0.0523<br><br>diet effect<br>p=0.0007 | fig legend | drug x diet interaction<br>F (1, 34) = 4.174<br><br>drug effect<br>F (1, 34) = 4.043<br><br>diet effect<br>F (1, 34) = 13.82 | fig legend | No PTX lean vs obese<br>p=0.0007<br><br>PTX lean vs obese<br>p=0.4100 |
| ED 7e | two-way ANOVA | fig legend | lean<br>n=10/3<br><br>lean PTX<br>n=11/3<br><br>obese<br>n=8/3<br><br>obese PTX<br>n=9/5 | # cells / # animals | fig legend | error bars are mean +/- SEM | drug x diet interaction<br>p=0.4180<br><br>drug effect<br>p=0.1338<br><br>diet effect<br>p=0.0191 | fig legend | drug x diet interaction<br>F (1, 34) = 0.6723<br><br>drug effect<br>F (1, 34) = 2.359<br><br>diet effect<br>F (1, 34) = 6.050 | fig legend | |
| ED 7f | two-way ANOVA | fig legend | lean<br>n=10/3<br><br>lean PTX<br>n=11/3<br><br>obese<br>n=8/3<br><br>obese PTX<br>n=9/5 | # cells / # animals | fig legend | error bars are mean +/- SEM | drug x diet interaction<br>p=0.0997<br><br>drug effect<br>p=0.2807<br><br>diet effect<br>p=0.8829 | fig legend | drug x diet interaction<br>F (1, 34) = 2.864<br><br>drug effect<br>F (1, 34) = 1.201<br><br>diet effect<br>F (1, 34) = 0.02203 | fig legend | |
| ED 7g | two-way ANOVA | fig legend | lean<br>n=10/3<br><br>lean PTX<br>n=11/3<br><br>obese<br>n=8/3<br><br>obese PTX<br>n=9/5 | # cells / # animals | fig legend | error bars are mean +/- SEM | drug x diet interaction<br>p=0.6176<br><br>drug effect<br>p=0.2711<br><br>diet effect<br>p=0.2711 | fig legend | drug x diet interaction<br>F (1, 34) = 0.2539<br><br>drug effect<br>F (1, 34) = 1.252<br><br>diet effect<br>F (1, 34) = 0.04215 | fig legend | |

Supplemental Table 1. Statistics table.

| | | | | | | | | | | | |
|---|---|---|---|---|---|---|---|---|---|---|---|
| ED 7h | two-way ANOVA | fig legend | lean n=10/3<br><br>lean PTX n=11/3<br><br>obese n=8/3<br><br>obese PTX n=9/5 | # cells / # animals | fig legend | error bars are mean +/- SEM | drug x diet interaction p=0.2340<br><br>drug effect p=0.8356<br><br>diet effect p=0.4694 | fig legend | drug x diet interaction F (1, 34) = 1.468<br><br>drug effect F (1, 34) = 0.04374<br><br>diet effect F (1, 34) = 0.5353 | fig legend | |
| ED 7i | two-way ANOVA followed by Sidaks multiple comparison | fig legend | lean n=10/3<br><br>lean PTX n=11/3<br><br>obese n=8/3<br><br>obese PTX n=9/5 | # cells / # animals | fig legend | error bars are mean +/- SEM | drug x diet interaction p=0.0232<br><br>drug effect p=0.0603<br><br>diet effect p=0.0603 | fig legend | drug x diet interaction F (1, 34) = 5.648<br><br>drug effect F (1, 34) = 3.777<br><br>diet effect F (1, 34) = 7.140 | fig legend | No PTX lean vs obese p=0.0028<br><br>PTX lean vs obese p=0.9715 |
| ED 8a | Unpaired T-test | fig legend | mCherry n=7<br><br>hM4D(Gi) n=9 | # animals | fig legend | error bars are mean +/- SEM | p=0.8910 | fig legend | t=0.1395, df=14 | fig legend | |
| ED 8b | Simple linear regression of slopes and y intercepts | fig legend | mCherry n=7<br><br>hM4D(Gi) n=9 | # animals | fig legend | error bars are mean +/- SEM | slope p=0.8752<br><br>y-intercept p=0.8944 | fig legend | slope: F (1,44) = 0.02496<br><br>y-intercept: F (1,45) =0.01782 | fig legend | |
| ED 8c | two-way RM ANOVA | fig legend | mCherry n=7<br><br>hM4D(Gi) n=9 | # animals | fig legend | error bars are mean +/- SEM | virus vs drug interaction p=0.9751<br><br>virus effect p=0.1327<br><br>drug effect p=0.1019 | fig legend | virus vs drug interaction F (1, 14) = 0.001013<br><br>virus effect F (1, 14) = 2.549<br><br>drug effect F (1, 14) = 3.064 | fig legend | |
| ED 8d | Unpaired T-test | fig legend | mCherry n=12<br><br>hM4D(Gi) n=13 | # animals | fig legend | error bars are mean +/- SEM | p=0.4867 | fig legend | t=0.7069, df=23 | fig legend | |
| ED 8e | three-way RM ANOVA | fig legend | mCherry n=12<br><br>hM4D(Gi) n=13 | # animals | fig legend | error bars are mean +/- SEM | day x virus x value interaction p=0.2493<br><br>virus x value interaction p=0.9620<br><br>day x value interaction p=0.6267<br><br>day x virus interaction p=0.8244<br><br>value effect p=0.5542<br><br>virus effect p=0.1394<br><br>day effect | fig legend | day x virus x value interaction F (2, 46) = 1.432<br><br>virus x value interaction F (1, 23) = 0.002321<br><br>day x value interaction F (1.890, 43.46) = 0.4546<br><br>day x virus interaction F (2, 46) = 0.1939<br><br>value effect F (1.000, 23.00) = 0.3603 | fig legend | |

Supplemental Table 1. Statistics table.

| | | | | | | | | | | | |
|---|---|---|---|---|---|---|---|---|---|---|---|
| | | | | | | | p=0.0041 | | virus effect<br>F (1, 23) = 2.344<br><br>day effect<br>F (1.924, 44.25) = 6.360 | | |
| ED 9a | Unpaired T-test | fig legend | lean<br>n=14<br><br>obese<br>n=16 | # animals | fig legend | error bars are mean +/- SEM | p<0.0001 | fig legend | t=7.818, df=28 | fig legend | |
| ED 9b | two-way ANOVA followed by Dunnett's multiple comparison | fig legend | lean<br>n=14<br><br>obese<br>n=16 | # animals | fig legend | error bars are mean +/- SEM | training schedule x diet interaction<br>p=0.0527<br><br>training schedule effect<br>p=0.0005<br><br>diet effect<br>p<0.0001 | fig legend | training schedule x diet interaction<br>F (2, 56) = 3.103<br><br>training schedule effect<br>F (1.441, 40.35) = 11.45<br><br>diet effect<br>F (1, 28) = 24.80 | fig legend | lean RR5 vs RR10<br>p=0.4182<br><br>lean RR5 vs RR20<br>p=0.0171<br><br>obese RR5 vs RR10<br>p=0.0554<br><br>obese RR5 vs RR20<br>p=0.0102 |
| ED 9c | two-way ANOVA mixed effects model | fig legend | lean<br>n=14<br><br>obese<br>n=16 | # animals | fig legend | error bars are mean +/- SEM | diet x drug interaction<br>p=0.8566<br><br>drug effect<br>p=0.9335<br><br>diet effect<br>p=0.4844 | fig legend | diet x drug interaction<br>F (1, 21) = 0.03347<br><br>drug effect<br>F (1, 21) = 0.007126<br><br>diet effect<br>F (1, 28) = 0.5022 | fig legend | |
| ED 9d | two-way RM ANOVA followed by Sidaks multiple comparison | fig legend | lean<br>n=7<br><br>obese<br>n=6 | # animals | fig legend | error bars are mean +/- SEM | diet x drug interaction<br>p=0.7383<br><br>drug effect<br>p=0.3906<br><br>diet effect<br>p=0.0018 | fig legend | diet x drug interaction<br>F (1, 11) = 0.1175<br><br>drug effect<br>F (1, 11) = 16.76<br><br>diet effect<br>F (1, 11) = 0.7986 | fig legend | vehicle lean vs obese<br>p=0.0172<br><br>NNC7-11 vehicle vs obese<br>p=0.0061 |
| ED 10 c | Unpaired T-test | fig legend | lean n=8<br><br>obese<br>n=7 | # animals | fig legend | error bars are mean +/- SEM | p<0.0001 | fig legend | t=7.531, df=13 | fig legend | |
| ED 10 d | two-way RM ANOVA followed by Dunnett's multiple comparisons | fig legend | lean n=8<br><br>obese<br>n=7 | # animals | fig legend | error bars are mean +/- SEM | training schedule x diet interaction<br>p=0.0238<br><br>training schedule effect<br>p=0.0031<br><br>diet effect<br>p<0.0001 | fig legend | training schedule x diet interaction<br>F (2, 26) = 4.331<br><br>training schedule effect<br>F (1.504, 19.55) = 9.122<br><br>diet effect<br>F (1, 13) = 32.23 | fig legend | lean RR5 vs RR10<br>p=0.9403<br><br>lean RR5 vs RR20<br>p=0.0662<br><br>obese RR5 vs RR10<br>p=0.0109 |

Supplemental Table 1. Statistics table.

| | | | | | | | | | | | |
|---|---|---|---|---|---|---|---|---|---|---|---|
| | | | | | | | | | | | obese RR5 vs RR20 p=0.0108 |
| ED 10 e | Two way ANOVA mixed effects analysis | fig legend | lean 589 n=8<br><br>lean 473 n=7<br><br>obese 589 n=6<br><br>obese 473 n=7 | # animals | fig legend | error bars are mean +/- SEM | diet x stimulation interaction p=0.2557<br><br>stimulation effect p=0.2510<br><br>diet effect p=0.8702 | fig legend | diet x stimulation interaction F (1, 11) = 1.438<br><br>stimulation effect F (1, 11) = 1.468<br><br>diet effect F (1, 13) = 0.02776 | fig legend | |
| ED 10f | two-way RM ANOVA | fig legend | lean n=6<br><br>obese n=3 | # animals | fig legend | error bars are mean +/- SEM | diet x stimulation interaction p=0.9167<br><br>stimulation effect p=0.3967<br><br>diet effect p=0.1078 | fig legend | diet x stimulation interaction F (1, 7) = 0.01176<br><br>stimulation effect F (1, 7) = 0.8146<br><br>diet effect F (1, 7) = 3.399 | fig legend | |
